# Selective ion binding and uptake shape the microenvironment of biomolecular condensates

**DOI:** 10.1101/2024.12.24.630169

**Authors:** Iris B. A. Smokers, Enrico Lavagna, Rafael V. M. Freire, Matteo Paloni, Ilja K. Voets, Alessandro Barducci, Paul B. White, Mazdak Khajehpour, Evan Spruijt

**Affiliations:** Institute for Molecules and Materials, Radboud University, Heyendaalseweg 135, 6523 AJ Nijmegen, The Netherlands; CBS (Centre de Biologie Structurale), Univ Montpellier, CNRS, Inserm, Montpellier, France; Laboratory of Self-Organizing Soft Matter, Department of Chemical Engineering and Chemistry, Eindhoven University of Technology, P.O. Box 513, 5600 MB Eindhoven, the Netherlands; Institute for Complex Molecular Systems, Eindhoven University of Technology, P.O. Box 513, 5600 MB Eindhoven, the Netherlands; Thomas Young Centre and Department of Chemical Engineering, University College London, London WC1E 7JE, United Kingdom; Department of Chemistry, University of Manitoba, Winnipeg, Manitoba, R3T 2N2, Canada

**Keywords:** Biomolecular condensates, coacervates, ion binding, partitioning

## Abstract

Biomolecular condensates modulate various ion-dependent cellular processes and can regulate subcellular ion distributions by selective uptake of ions. However, the molecular grammar governing condensate-ion interactions is poorly understood. Here, we use NMR spectroscopy of ions and model condensate components to quantify and spatially resolve selective ion binding to condensates and show that these interactions follow the law of matching water affinities, resulting in strong binding between proteins and chaotropic anions, and between nucleic acids and kosmotropic cations. Ion uptake into condensates directly follows binding affinities, resulting in selective uptake of strong-binding ions, but exclusion of weak-binding ions. Ion binding further shapes the condensate microenvironment by altering the composition, viscosity and interface potential. Such changes can have profound effects on biochemical processes taking place inside condensates, as we show for RNA duplex formation. Our findings provide a new perspective on the role of condensate-ion interactions in cellular bio- and electrochemistry and may aid design of condensate-targeting therapeutics.

## Introduction

Biomolecular condensates are phase-separated compartments that regulate a variety of cellular processes, including ribosome biogenesis, stress response and protein aggregation.^1,2^ They are formed through multivalent interactions between proteins (and RNA), resulting in droplets that are enriched in biomolecules and have distinct local physicochemical environments. These environments can differ in hydrophobicity, viscosity and the presence of specific ions, and drastically alter biochemical processes.^3,4^ By uptake of divalent metal ions such Mg^2+^ and Cu^2+^, condensates regulate both ribozyme function^5,6^ and protein aggregation^2^, and could alter the reactivity of enzymes that require metal ions in their active site.^7–11^ In addition, the differential uptake of ions into condensates has recently been shown to modulate subcellular ion distributions, and may thereby play a role in regulating the electrochemistry of cells.^12–14^ Outside the cell, condensates can also be used to remove heavy metals and other ions from wastewater.^15–17^

Ions also modulate the stability of condensates. Condensate formation is often driven by charge-charge interactions, and counterion release makes their formation entropically favorable.^18–21^ This makes condensates susceptible to salt, as ions screen charges^22,23^ and can give rise to re-entrant phase separation.^24^ In addition, several marine species use condensates as underwater adhesives^25,26^ and extracellular hard tissues,^27,28^ and the exposure to sea water alters their physical state from liquid to gel-like. Furthermore, condensates are hypothesized to have played a role in the origins of life as a first generation of cell-like compartments – or protocells – in the salty prebiotic soup.^29–33^

These striking effects of ions on condensates have sparked a growing interest in measuring ion uptake in condensates.^14,23,42–48,34–41^ However, the molecular mechanisms behind ion interactions with condensates remain poorly understood. Binding of ions to specific regions in the condensate components could alter the conformation of biomolecules inside condensates, could lead to selective uptake of ions and charged cofactors and therapeutics, and could create distinct nanoscopic environments that foster specific reactions and alter condensate material properties. Therefore, insight into the molecular “grammar” of condensate-ion interactions is essential for a full understanding of condensate function and the potential of novel condensate-targeting therapeutics.^49–51^

In this work, we investigate the molecular mechanisms by which salt ions interact with the components of model condensates and how they affect the local environment inside the droplets. We investigate a range of ions with different properties: from weakly hydrated (chaotropes) to strongly hydrated (kosmotropes). Through NMR binding assays, we find that chaotropic anions and kosmotropic cations selectively bind to the condensate components, following the law of matching water affinities (LMWA), a trend we postulate is universal for all condensates. Molecular dynamics simulations and SAXS measurements further show that the binding of ions to the peptide is sequence specific and results in compaction of the peptide chain. Using a combination of ^1^H, ^7^Li, ^13^C, ^19^F, ^23^Na, ^25^Mg, ^31^P, ^35^Cl, ^39^K, ^81^Br and ^133^Cs-NMR spectroscopy, we can quantify the full composition of the condensate phase with a single technique, and show that the partitioning of ions directly follows their binding strength, resulting in preferential uptake of strong-binding ions but exclusion of weak-binding ions, in contrast to theoretical predictions.^52,53^

The binding and uptake of ions has a drastic effect on the properties of the condensates. Strong-binding ions remodel the phase diagram by effectively neutralizing charges and favouring other interactions such as π-π stacking, which can result in re-entrant phase separation. Selective ion binding further modulates the condensate viscosity and can flip the interface potential of condensates, which can impact their interaction with membranes and other intracellular structures. Lastly, we show that the altered microenvironment can affect biochemistry inside the condensates: kosmotropic cations can significantly stabilize RNA duplexes inside condensates whereas the same ions destabilize RNA duplexes in solution. Taken together, our results show that ions can affect the condensate composition and microenvironment in dramatic and nontrivial manners, and we provide molecular insight into how specific ions bind selectively to condensates to shape their properties.

## Results

### 1. Selective binding of ions to condensate components

Biomolecular condensates are composed of proteins that often contain significant disordered regions, and nucleic acids. To enable a systematic and high-resolution investigation of the specific binding of different salt ions to these condensate components, we used a peptide/nucleotide condensate model system, made with 1 mM protamine chloride / 25 mM sodium adenosine 5’-triphosphate (ATP) (Figure 1a).^54–58^ We prepared condensates with a series of different anions and cations (all 100 mM, charge-based, Figure 1a, Supplementary Figure 2) ranging from strongly kosmotropic (strongly hydrated) to strongly chaotropic (weakly hydrated, Figure 1i). Ions can be classified according to their hydration strength, because charge-charge interactions in water are mediated by water. This is one of the factors reflected in the Hofmeister series.^59^

**Figure 1:**
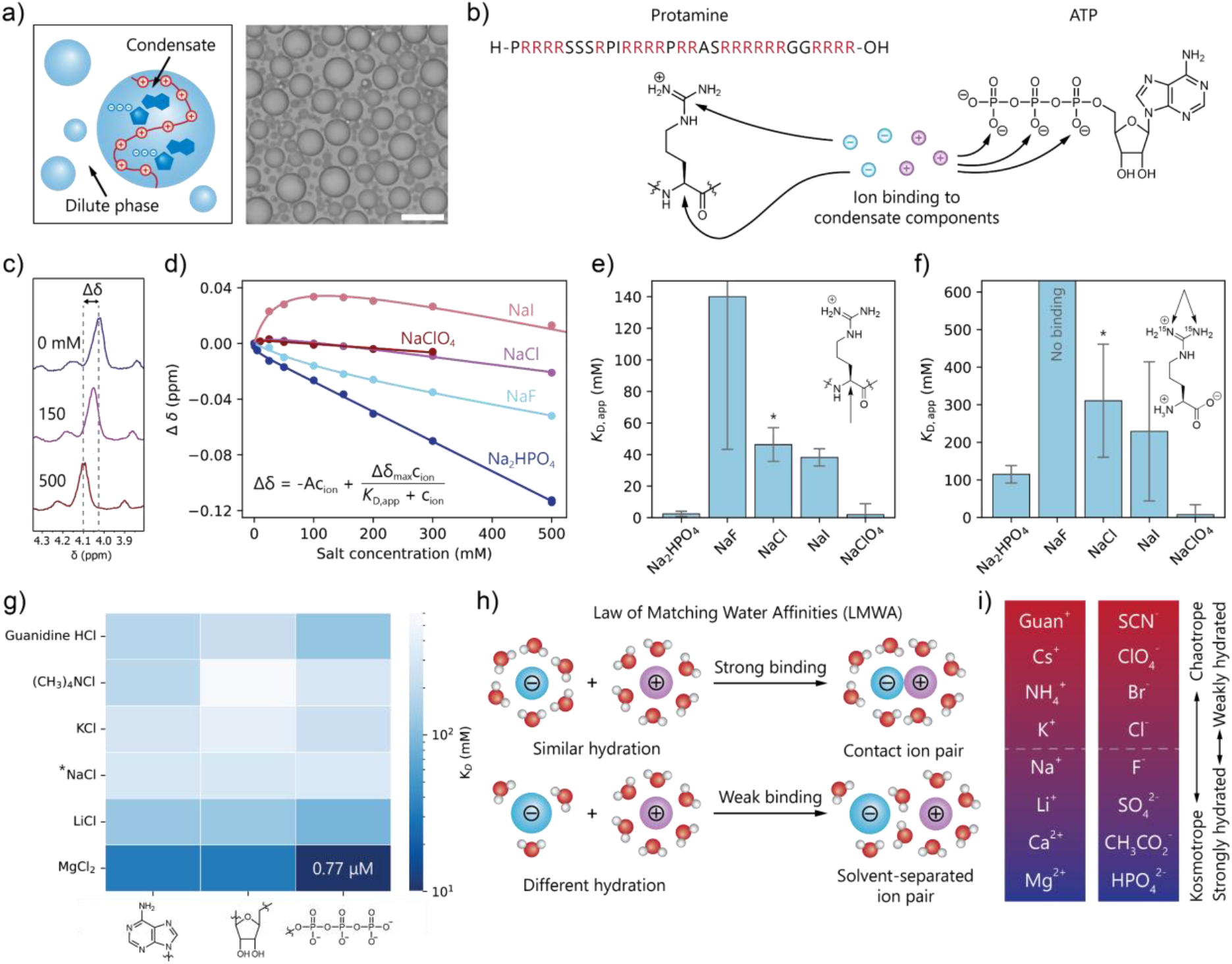
Ion binding to the condensate components protamine and ATP follows the law of matching water affinities (LMWA) and valency. **a)** The arginine-rich peptide protamine and ATP form model biomolecular condensates, scalebar = 20 μm. **b)** Through NMR-based binding assays we investigated the binding of ions to the condensate components protamine and ATP in absence of condensates. **c)** ^1^H-NMR chemical shifts of the arginine α-proton as a function of salt concentration at 5°C. **d)** The change in chemical shift (Δδ) as a function of salt concentration for different anions was fitted to a binding isotherm from Rembert *et al*.^60^ to obtain the apparent dissociation constant *K*_D,app_. The other binding curves can be found in Supplementary Information Section 3.2. **e)** Apparent dissociation constants of anions for the α-proton. Error bars represent the error of the fit. *Because the sample already contains 23.1 mM Cl^-^ at 0 mM added salt, this amount was added to the *K*_D,app_. **f)** Apparent dissociation constants of anions for the guanidinium groups in arginine. Chaotropic and divalent anions bind strongly to the arginines in protamine. Error bars represent the error of the fit. *Because the sample already contains 61.8 mM Cl^-^ at 0 mM added salt, this amount was added to the *K*_D,app_. **g)** Apparent dissociation constants of cations for the phosphate, ribose and nucleobase of ATP. Kosmotropic and divalent cations bind strongly to the phosphates on ATP.*Because the sample already contains 92.9 mM Na^+^ at 0 mM added salt, this amount was added to the *K*_D,app_. **h)** Binding of ions to the condensate components follows the law of matching water affinities (LMWA). **i)** Trends in hydration strength for cations and anions.

We quantified the interaction strength between the ions and the individual condensate components in dilute solution using an NMR-based binding assay (Figure 1b).^60^ We measured the change in ^1^H, ^15^N or ^1^P-NMR chemical shift as a function of salt concentration (Figure 1c,d) to determine ion binding to the α-proton and guanidinium moiety of the arginines in protamine, and to the phosphates, sugar and nucleobase in ATP. Equation 1 describes the effect of ions on the NMR chemical shift: a linear term accounts for the non-specific shift due to an increase in ionic strength and change in water activity, and a Langmuir isotherm accounts for specific ion binding:

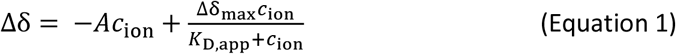

In this equation, Δδ is the change in chemical shift with respect to the sample without added salt, *A* is a constant, *c*_ion_ is the concentration of the salt ion of interest, Δδ_max_ is the maximum chemical shift change due to binding at saturation and *K*_D,app_ is the apparent dissociation constant. *A*, Δδ_max_ and *K*_D,app_ were obtained by fitting.

Using this method for sodium salts with different anions, we found that chaotropic anions such as ClO_4_^-^ bind surprisingly strongly to both the α-protons and guanidiniums of arginines, with low-mM apparent dissociation constants in both cases, while kosmotropic anions such as F^-^ do not bind or only bind weakly (Figure 1e,f). The strongly kosmotropic HPO_42-_, however, binds to both the α-proton and the guanidinium, indicating that valency and possibly hydrogen-bonding^61^ also play a role in binding strength. It is interesting to note that different ions have a different sign of the non-linear part of Δδ, which suggests that they have different mechanisms of binding (Supplementary Figure 4 & 6).

For ATP, instead, we observe that kosmotropic cations such as Mg^2+^ and Li^+^ bind strongly to the phosphates with sub-μM *K*_D,app_ for Mg^2+^ (Figure 1g, Supplementary Figure 9-10), while the weakly kosmotropic Na^+^, and chaotropic K^+^ and (CH_3_)_4_N^+^ have much weaker binding. The observed binding of Mg^2+^ and Li^+^ to the ribose and nucleobase is most likely due proximity effects of the phosphates. We also observe binding for the strongly chaotropic guanidinium, likely because of hydrogen-bonding interactions.^61^

The binding assay shows that chaotropic monovalent anions interact strongly with the arginines in protamine, which are also chaotropic, while kosmotropic cations interact strongly with the phosphates on ATP, which are also kosmotropic. These observations follow the law of matching water affinities (LMWA), as proposed by Collins.^62,63^ Because water has a high dielectric constant, long-range electrostatic forces disappear for ions in water. In these regimes, short-range effects such as the water affinity of ions take over. According to the LMWA, the water affinity, or hydration enthalpy, of ions determines their interaction strength with other ions. Oppositely charged ions with matching water affinities interact most strongly (Figure 1h): kosmotropes form contact ion pairs because of stronger electrostatic attraction due to their high charge-density, while chaotropes form contact ion pairs to release unfavourably bound hydration water.^63^ Interaction between a kosmotrope and a chaotrope, however, is less favourable and these combinations do not tend to form strong ion pairs.

Interestingly, all organic cations in nature (i.e. guanidinium and ammonium) are chaotropic, while all organic anions (i.e. phosphates, carboxylic acids and sulfates) are kosmotropic (Figure 1i). Therefore, any protein in the condensate with positive charges is expected to bind chaotropic anions such as ClO_4_^-^, I^-^ and Br^-^, and any negatively charged protein or nucleic acid is expected to bind kosmotropic cations such as Li^+^ and Mg^2+^. The valency of the ions also influences their binding to condensate components: multivalent, kosmotropic phosphates do bind to the chaotropic arginines in protamine.

### 2. Ion binding is sequence specific and compacts protamine

To obtain a deeper insight into the ion binding to condensate components, we ran all-atom explicit solvent Molecular Dynamics (MD) simulations of protamine with different anions. Radial distribution functions (Figure 2a, b) confirm that ClO_4_^-^ and Cl^-^ bind to the backbone α-protons and amide NHs, and to the guanidinium groups of arginines in protamine, with an apparent *K*_D_ that is in qualitative agreement with the NMR binding assays (Figure 2c, Supplementary Figure 11), while F^-^ shows no significant backbone binding.

**Figure 2:**
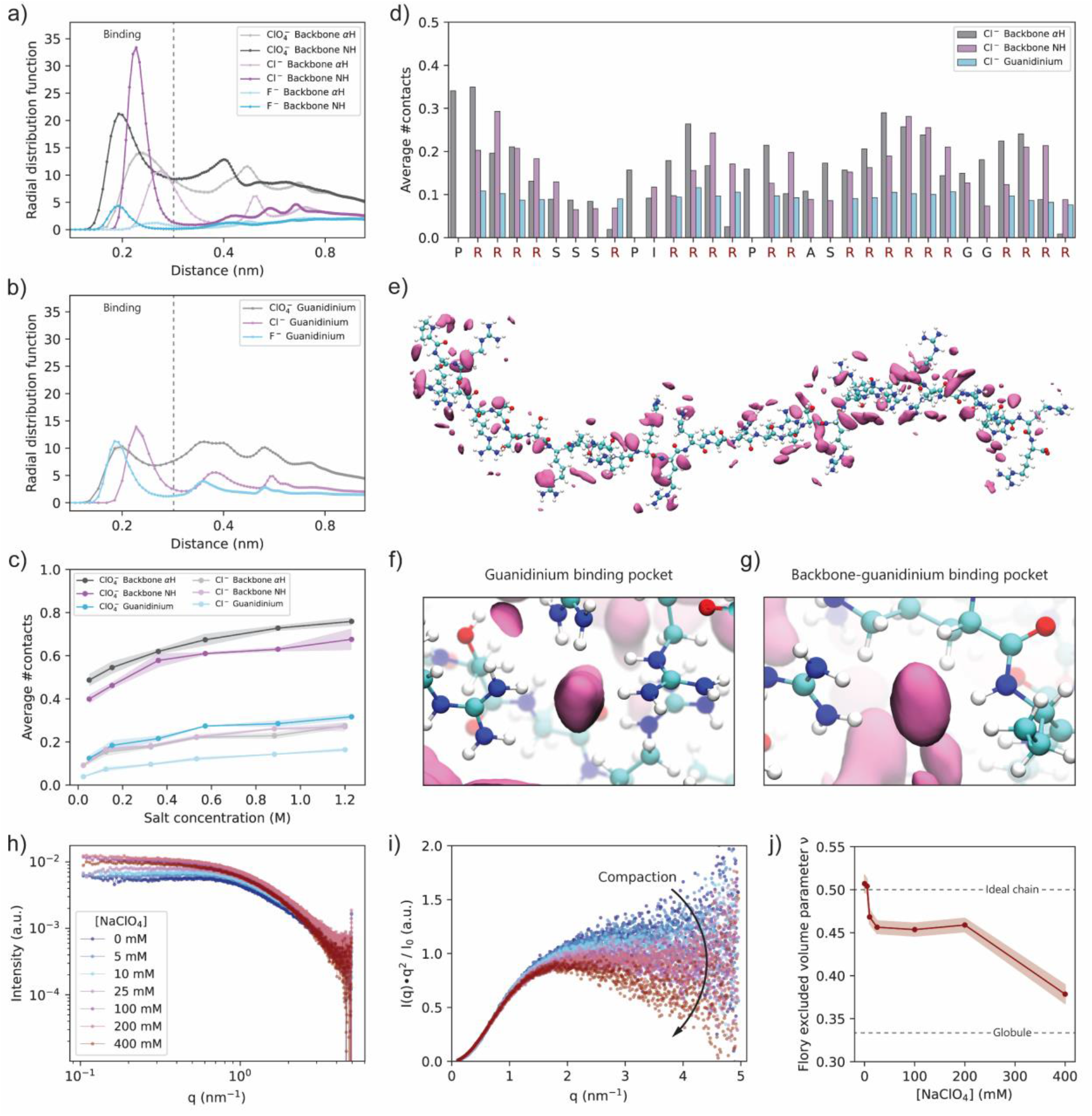
Ion binding to protamine is sequence specific and causes compaction of the peptide chain. **a,b)** Radial distribution functions (RDF) of NaF, NaCl and NaClO_4_^-^ with respect to **a)** the arginine backbone α-proton (αH) and NH and **b)** guanidinium as obtained from all-atom explicit solvent Molecular Dynamics (MD) simulations at 300 mM added salt concentration. Dotted lines represent the cut-off radius for binding, which differs slightly per type of ion. **c)** Average number of protamine hydrogen-ion contacts as a function of the ion concentration. **d)** Average number of hydrogen-ion contacts per residue in protamine for 300 mM NaCl. **e)** Snapshot of the whole protamine sequence with highlighted regions of high ion density, from a simulation with the protamine constrained in the open configuration at 500 mM added NaCl concentration. The mauve solids represent isosurfaces with isovalue 40x the bulk ion concentration. The peptide is represented with white hydrogen, cyan carbon, blue nitrogen and red oxygen atoms. **f, g)** Snapshots showing the binding pockets with high affinity for ions. Both are taken from a simulation with the protamine constrained in the open configuration at 500 mM added NaCl concentration. **h)** SAXS scattering curves for 1 mM protamine with different concentrations of NaClO^4^. **i)** Kratky representation of the SAXS data. The curves progressively bend downwards for q > 1.5 nm^-1^ for increasing NaClO^4^ concentrations, indicating a compaction of the protamine chain. **j)** The Flory excluded volume parameter ν, obtained from fitting the SAXS curves with a generalized Gaussian chain model, decreases slightly from 0 to 25 mM NaClO^4^ indicating compaction of the chain. The decrease at 400 mM is likely due to complexation of multiple protamines.

Analysis of the ion distribution per residue (Figure 2d) reveals that ions bind specifically to arginine-rich regions, where multivalent ‘binding pockets’ are temporally formed between neighbouring guanidinium groups (Figure 2e,f), and/or backbone NH and α-protons (Figure 2g). In arginine-poor regions, binding to the backbone is significantly reduced, showing that ion binding is sequence-specific and is a cooperative effect between the backbone and guanidinium protons.

Analysis by small angle X-ray scattering (SAXS) shows that the strong binding of chaotropic anions to protamine induces compaction of the peptide. Scattering curves of protamine for high NaClO_4_ concentrations exhibit a sharper decay at high *q* than for low NaClO_4_ concentrations (Figure 2h) and bend downwards from q > 1.5 nm^-1^ in the Kratky plot (Figure 2i). By contrast, scattering curves of protamine with NaF do not show any dependence on salt concentration (Supplementary Figure 13). The scattering curves were fitted with a generalized Gaussian chain model with Flory excluded volume parameter ν to quantify the degree of compaction (Figure 2j). These fits indeed demonstrate that the protamine behaves like an ideal Gaussian chain (ν = 0.5) at very low NaClO_4_ concentration, followed by moderate compaction by ClO_4_^-^ binding at intermediate NaClO_4_ concentrations (ν = 0.45), and strong compaction at high NaClO_4_ concentration (ν = 0.37), demarcating the potential onset of homotypic protamine clusters (vide infra).

### 3. Ion partitioning into condensates follows LMWA

Selective binding of salt ions to individual condensate components could result in differential partitioning of these ions into the condensates themselves. Therefore, we investigated the partitioning of a wide range of salt ions into the protamine/ATP condensates. Interestingly, many salt ions have an NMR-active nucleus, allowing us to measure their concentration simultaneously with all other condensate components with a combination of ^1^H, ^7^Li, ^13^C, ^19^F, ^23^Na, ^25^Mg, ^31^P, ^35^Cl, ^39^K, ^81^Br and ^133^Cs-NMR spectroscopy. Classical approaches to measure the ion content of condensates have the disadvantage that either separate techniques are required to measure the ions and other condensate components (e.g., ICP-MS in combination with HPLC)^14,34,42,45^ or that they cannot separately measure the anion and cation concentration (e.g., thermogravimetric analysis or tie-line analysis).^35,37,44^ The latter can lead to incorrect conclusions in cases where the anion and cation partition differently. Our NMR-based approach, however, allows us to individually measure all condensate components – the salt anion and cation, protamine, ATP and buffer – in both the condensate and dilute phase with a single technique for a wide variety of salts.

Using this method, we observe distinct partitioning behaviour for different ions (Figure 3c,f), which directly follows the binding strengths measured in Section 1. For the anions we observe that the chaotropic (ClO_4_^-^ and SCN^-^) and divalent anions (HPO_42-_), which bind strongly, are included, whereas kosmotropic anions (F^-^) are excluded. For the monovalent anions, the partitioning correlates linearly with their hydration strength (Figure 3d). The transition point between inclusion and exclusion of the investigated ions coincides with a Jones-Dole coefficient *B* = 0, which marks the transition between weakly and strongly hydrated ions,^64^ indicating that weakly hydrated anions in general are included, while strongly hydrated anions in general are excluded. Inclusion of anions in the condensates also changes the condensate composition, reflected by the ratio protamine:ATP (Figure 3e), to retain overall charge neutrality of the condensate (Supplementary Figure 47). To confirm that our bulk measurements represent the ion distribution inside individual condensate droplets, we used Raman microscopy to measure the concentration of SCN^-^ inside individual condensate droplets and observe that SCN^-^ indeed localizes to the condensate and is evenly distributed throughout the droplet (Supplementary Figure 49).

**Figure 3:**
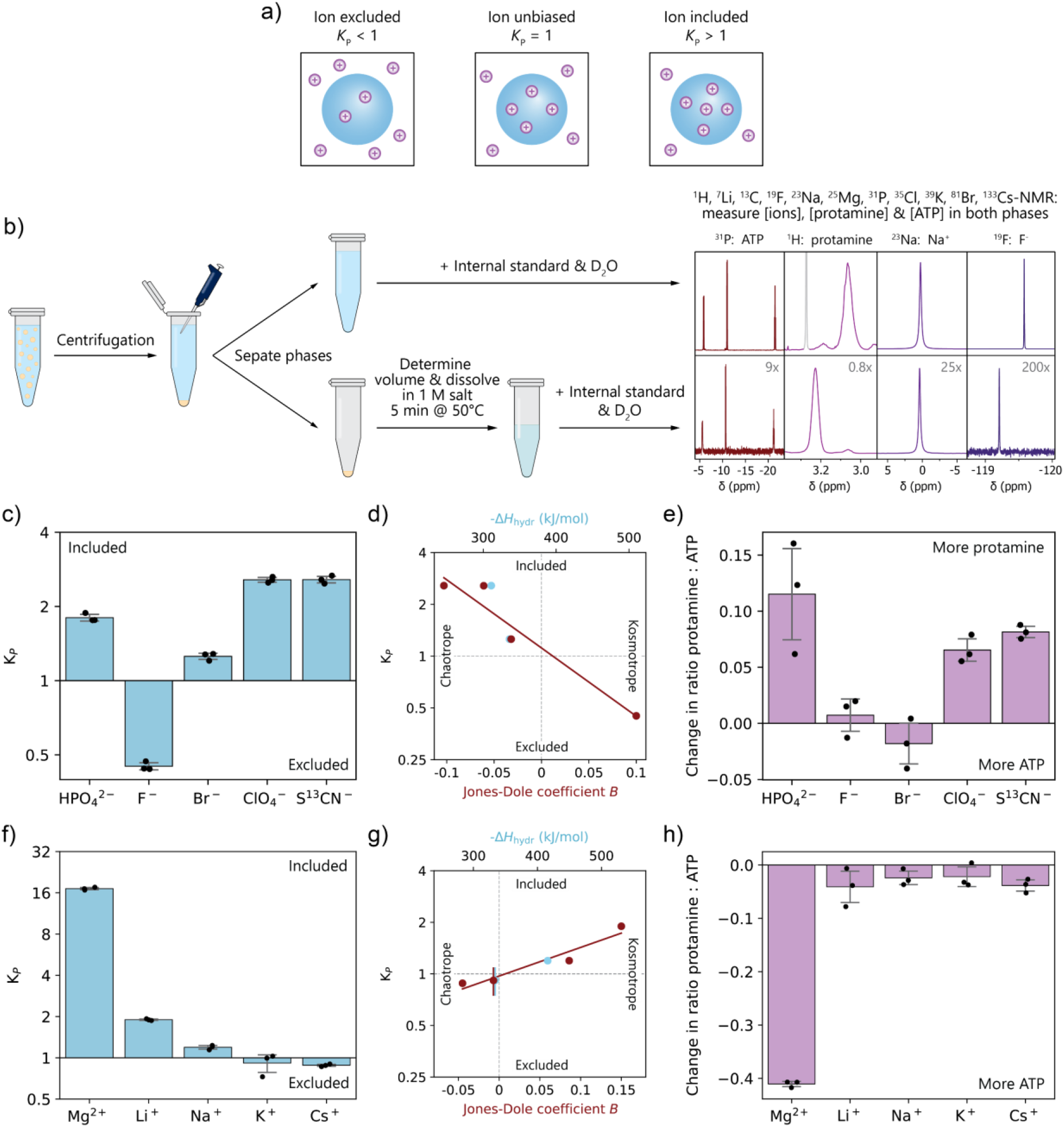
Protamine/ATP condensates selectively partition salt ions that bind to the condensate components. **a)** Ions can either be excluded, have no bias, or be included in the condensate phase. **b)** Graphical representation of the experimental procedure to measure ion- and condensate component partitioning by ^1^H, ^7^Li, ^13^C, ^19^F, ^23^Na, ^31^P, ^35^Cl, ^39^K, ^81^Br and ^133^Cs-NMR. The NMR spectra are for the dilute (top) and condensate (bottom) phase of protamine/ATP condensates with 100 mM NaF. **c)** Partitioning of 100 mM of different anions into protamine/ATP condensates. Chaotropic and divalent anions are included, while kosmotropic anions are excluded. **d)** Partitioning of monovalent anions as a function of Jones-Dole coefficient *B* and hydration enthalpy.^63,65^ **e)** Change in ratio of protamine:ATP inside the condensate phase upon ion partitioning to maintain overall charge balance of the condensates. **f)** Partitioning of 100 mM of different cations into protamine/ATP condensates. Kosmotropic and divalent cations are included, while chaotropic cations are excluded. **g)** Partitioning of monovalent cations as a function of Jones-Dole coefficient *B* and hydration enthalpy.^63,65^ **h)** Change in ratio of protamine:ATP inside the condensate phase upon ion partitioning to maintain overall charge balance of the condensates.

We observe a similar, but reversed, trend for the partitioning of cations (Figure 3f). Kosmotropic cations (Li^+^ and Mg^2+^) are included, while chaotropic cations (K^+^ and Cs^+^) are weakly excluded. Again, the transition between inclusion and exclusion coincides with *B* = 0 (Figure 3g), indicating that kosmotropic cations in general are included in condensates, while chaotropic cations in general are excluded. Like anions, cations also change the condensate composition (Figure 3h). In particular kosmotropic cations compete with protamine for interaction with ATP, and decrease the condensate protamine content.

Multivalent ions have stronger interaction with the condensate components, and therefore they show enhanced partitioning. However, water affinity and the corresponding LMWA still influence their extent of partitioning, as is shown by the different partitioning of HPO_4_^2-^ and Mg^2+^.

### 4. Selective ion binding remodels condensate phase diagrams

It is well-known that salt can modulate condensate stability by screening charges and lowering the entropic driving force for condensate formation.^22,38,66^ The critical salt concentration (CSC, typically NaCl) is therefore a conventional estimate of heterotypic condensate stability. However, the differential partitioning we found suggests that, unlike existing theories predict,^52,53^ different ions can influence the phase diagram and stability of condensates in very different ways.

We investigated the effect of selective ion binding on the stability and shape of the phase diagram of the protamine/ATP condensates. To this end, we used our NMR spectroscopy method to measure 5- or 6-dimensional phase diagrams containing the concentrations of protamine, ATP, Tris and 2-3 salt ions, allowing us to assess changes in the total condensate and dilute phase composition as a function of salt concentration.

The obtained phase diagrams for LiCl, NaF, NaCl and NaClO_4_ (Figure 4a-f, Supplementary Information Section 3.5.1) show marked differences. The tie-lines (Figure 4a-c) confirm our findings from Section 3 that F^-^ is excluded, while Li^+^, Cl^-^ and ClO_4_^-^ are included for all salt concentrations. For Cl^-^ and ClO_4_^-^, which both bind to protamine, the inclusion becomes stronger for higher salt concentrations, and this effect is stronger for the more strongly binding ClO_4_^-^.

**Figure 4:**
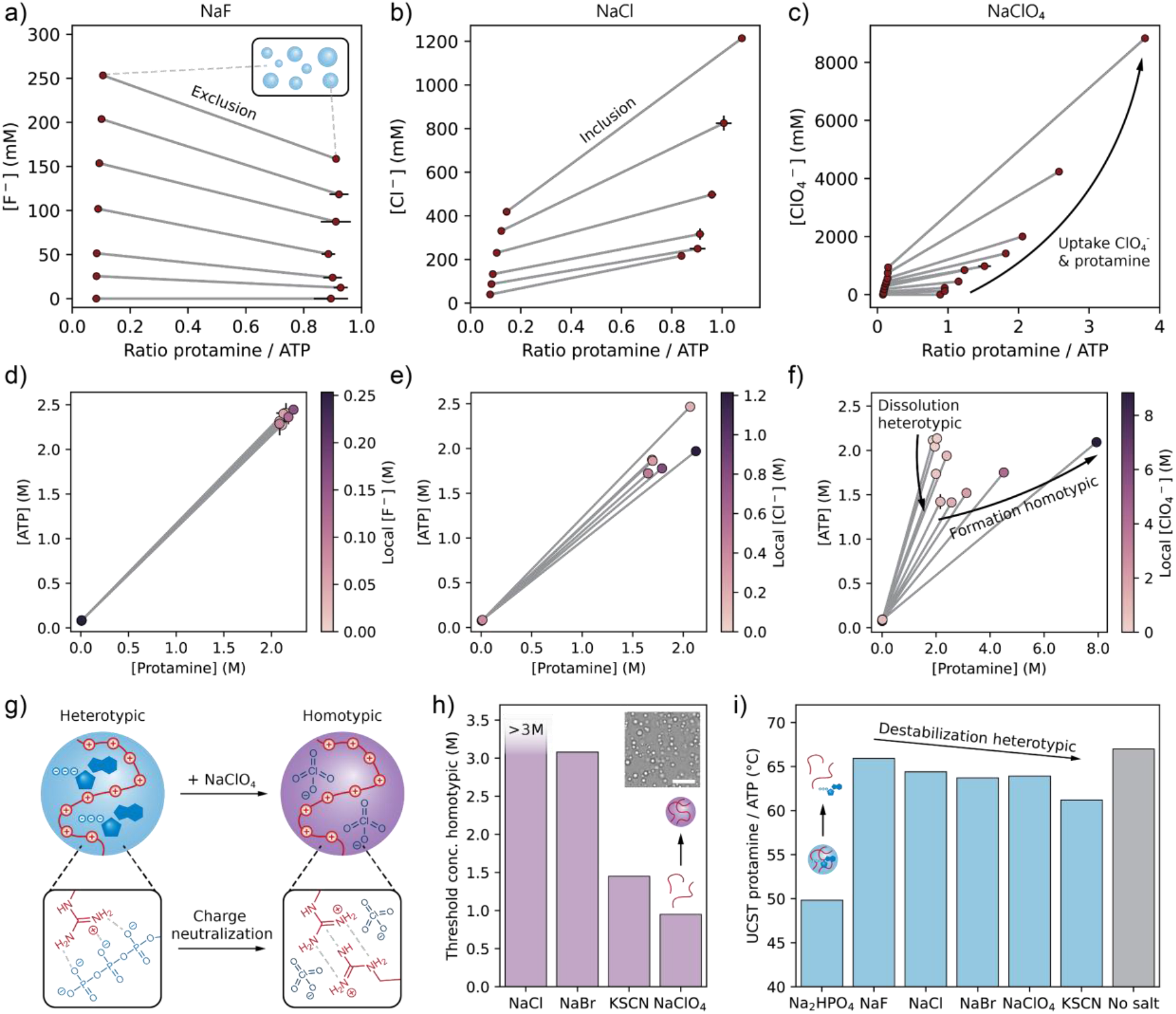
Selective ion binding remodels condensate phase diagrams. **a-c)** Phase diagrams of protamine/ATP with increasing concentrations of **a)** NaF, **b)** NaCl or **c)** NaClO_4_. Concentrations represent the charge concentrations of each species. Tie-lines are drawn in grey. **d-f)** [ATP] vs [protamine] representation of the same phase diagrams, showing the dissolution of heterotypic protamine/ATP condensates and subsequent formation of homotypic protamine/ClO_4_^-^ condensates for increasing concentrations of NaClO_4_. **g)** Schematic illustration of the transition from heterotypic protamine/ATP condensates to homotypic protamine condensates due to charge neutralization by ClO_4_^-^. **h)** Chaotropic anions lower the threshold concentration for formation of homotypic protamine condensates. Inset: Brightfield microscopy image of protamine/ClO_4_^-^ homotypic condensates. Scalebar is 10 μm. **i)** Upper critical solution temperature (UCST) for protamine/ATP condensates in presence of 100 mM of different salts. Chaotropic anions screen protamine’s charges and thereby destabilize heterotypic condensates.

For LiCl, NaF and NaCl, the condensates dissolve around 500 mM salt and we observe no change in the protamine:ATP ratio inside the condensates as a function of salt. For NaClO_4_, however, we instead observe a transition from heterotypic condensates that are rich in protamine and ATP, to homotypic protamine condensates that are neutralized by ClO_4_^-^ (Figure 4g). Arginine-rich peptides are known to form homotypic condensates at high salt concentrations, mediated by π-π stacking interactions between the arginine moieties.^24^ Due to the strong binding of ClO_4_^-^ to protamine, the multivalent ATP is gradually replaced by the monovalent ClO_4_^-^ (Figure 4c,f), while ClO_4_^-^-induced compaction (Figure 2h-j) increases the local protamine concentration. As a result, homotypic condensates are formed that now localize Cl^-^, Na^+^ and Tris more strongly than before (Supplementary Figure 52). Using a plate reader turbidity assay we determined the threshold salt concentration required to form homotypic protamine-anion condensates in absence of ATP, and observe that the ability of anions to promote homotypic condensate formation is highly dependent on their chaotropicity (Figure 4h), as strongly binding anions can more effectively neutralize the positive charges on protamine.

The effect of ions on condensates is, however, more nuanced than this. Anions that bind strongly to protamine may promote homotypic phase separation, but they destabilize charge-balanced heterotypic protamine/ATP condensates. In presence of 100 mM of anions, the upper critical solution temperature (UCST) is significantly reduced for strongly binding and divalent ions HPO_4_^2-^, ClO_4_^-^ & SCN^-^ (Figure 4i), while it is hardly affected by F^-^.

Interestingly, specific ion binding can also favour phase separation at a charge imbalance. When ATP is titrated into a solution of protamine and ions, there is a large excess of the positively charged protamine. Anions that strongly bind to protamine lower the excess of positive charge and thereby favour phase separation with ATP and lower the critical ATP concentration required for condensate formation ([ATP]_crit_) (Supplementary Figure 57). Cations that bind to ATP, however, increase the charge imbalance by neutralizing negative charge on ATP and thereby increase the [ATP]_crit_. Ions that do not bind to the condensate components generally also increase the [ATP]_crit_ due to nonspecific charge-screening effects.

These results show that specific ion binding modulates condensate phase diagrams in a multitude of ways. We observe similar trends for other heterotypic condensate systems (Supplementary Information Section 3.5.3), and for a homotypic peptide-based condensate (Supplementary Information Section 3.5.4).

### 5. Ion binding changes viscosity and interface potential of condensates

Crucially, selectively binding ions can also affect the physical properties and function of condensates. We measured the viscosity and interface potential (*ζ*-potential) of protamine/ATP condensates in the presence of different salts. Using Raster Image Correlation Spectroscopy (RICS), we observe that most salts lower the viscosity of the protamine/ATP condensates (Figure 5a), as expected.^67^ However, strongly binding ions, such as ClO_4_^-^ and Mg^2+^, increase the viscosity by neutralizing charges on the protamine and ATP, facilitating π-π and hydrogen-bonding interactions between protamines or ATPs, respectively, and compacting protamine. Interestingly, we observed previously (Figure 4i, Supplementary Figure 48) that these ions destabilize the protamine/ATP condensates at charge-balance, showing that there is no direct linear relation between condensate stability and viscosity.

**Figure 5:**
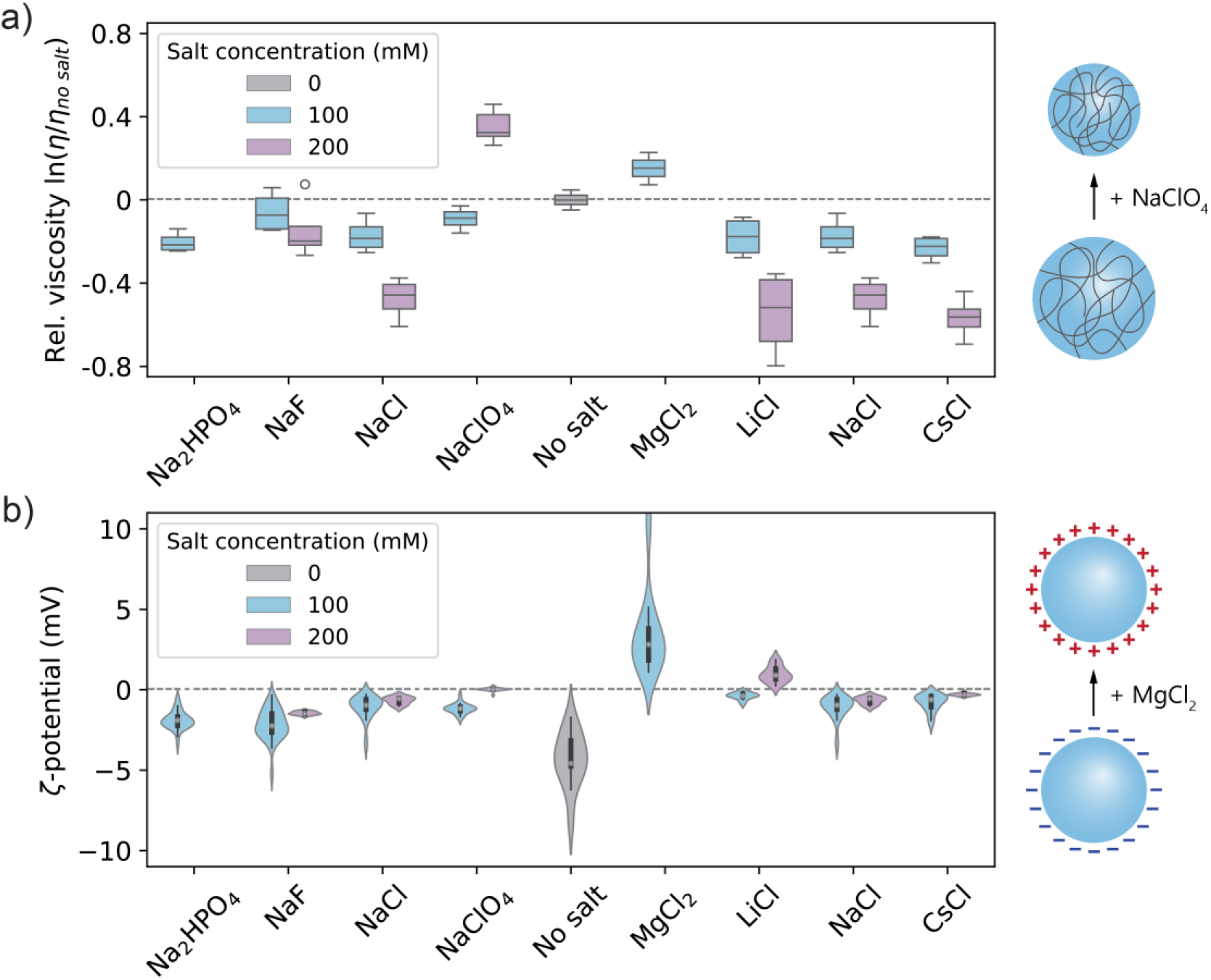
Ion binding and partitioning changes condensate viscosity and interface potential. **a)** Ions that bind strongly to the condensate components (NaClO_4_ and MgCl_2_) increase the condensate viscosity, even though they disfavour phase separation at charge balance. Outliers are represented as empty circles. **b)** Strongly binding cations (Mg_2+_ and Li_+_) can flip the interface (*ζ*-)potential of protamine/ATP condensates from negative to positive.

Using microelectrophoresis,^68^ we measured the *ζ*-potential of condensates with different salts. Protamine/ATP condensates naturally have a negative *ζ*-potential (−4.3 mV), and for most salts the *ζ*-potential remains negative (Figure 5b). However, addition of either 100 mM MgCl_2_ or 200 mM LiCl, flips the *ζ*-potential to positive (+3.5 or +1.0 mV, respectively), showing that binding of specific ions can also invert the interface potential. Importantly, this charge reversal contradicts the observed decrease in protamine:ATP ratio inside the condensates (Figure 3h), indicating that the interface potential is not always determined by the macromolecular components, but instead small ionic components may localize at the interface and dictate the interface potential.

### 6. Ion binding alters RNA duplex stability in condensates

Considering that the local environment inside condensates can have profound effects on biochemical processes inside,^3,4,69,70^ we investigated the stability of RNA- and DNA duplexes inside the protamine/ATP condensates. We used complementary decamer strands labelled with a Cy3/Cy5 Förster resonance energy transfer (FRET) pair, and analysed the FRET intensity inside the condensates and in buffer using confocal fluorescence microscopy (Figure 6a) for condensates with 100 mM of different salts.

**Figure 6:**
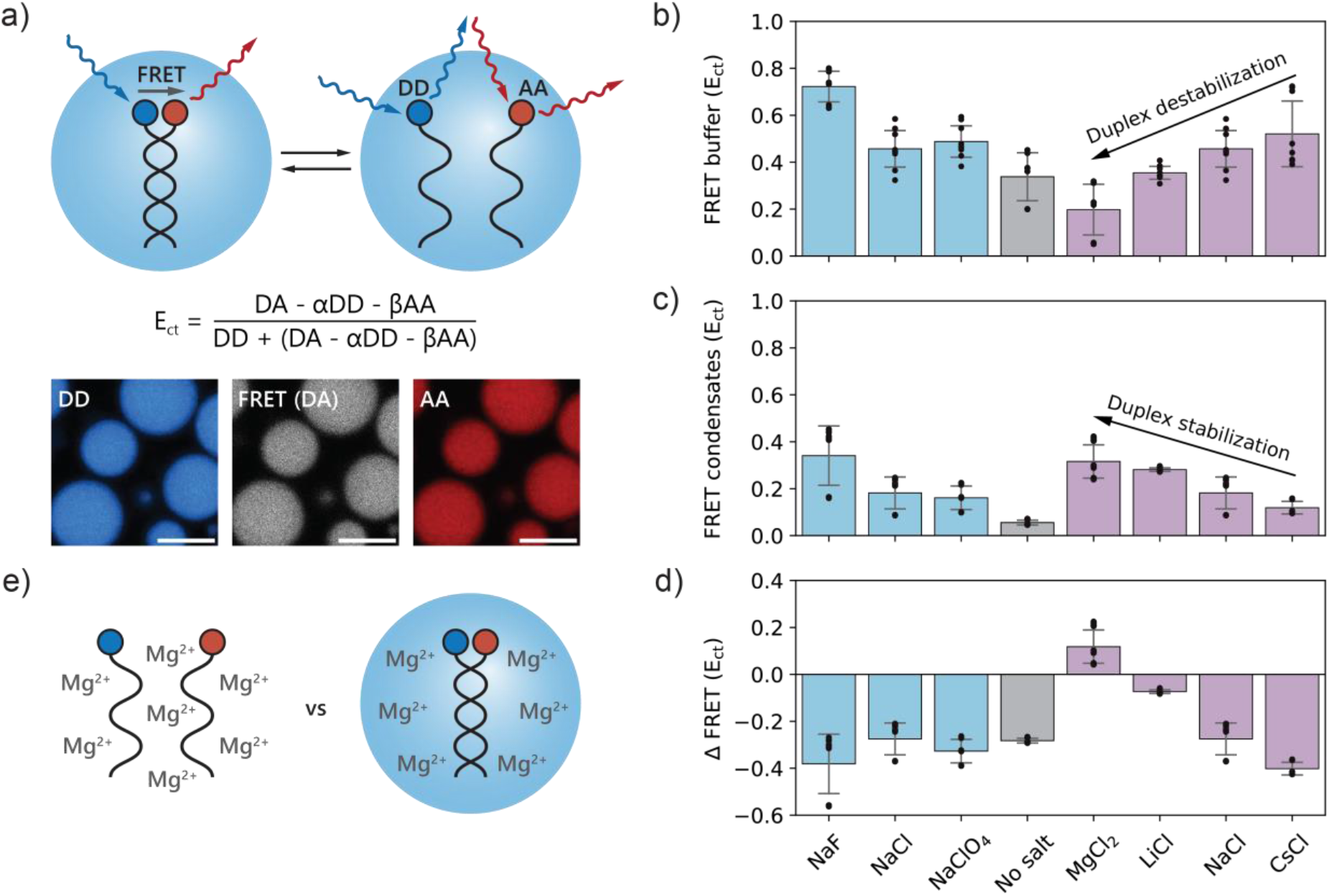
Ion binding and partitioning affects RNA duplex formation inside condensates. **a)** Schematic illustration of FRET analysis of the duplex stability of a Cy3/Cy5-FRET pair labelled set of complementary decamer RNAs. Fluorescence microscopy images of the Cy3 donor-donor excitation and emission (DD), Cy3 donor excitation and Cy5 acceptor emission (FRET, DA) and Cy5 acceptor-acceptor excitation and emission (AA) in protamine/ATP condensates with 100 mM NaCl. Scalebar is 10 μm. **b)** Corrected FRET efficiency of RNA in buffer. **c)** Corrected FRET efficiency inside protamine/ATP condensates with 100 mM salt. **d)** Change in FRET efficiency between the condensate and dilute phase (ΔFRET = FRET efficiency condensate – FRET efficiency buffer). **e)** Schematic illustration of the stabilization of RNA duplexes in condensates in presence of Mg_2+_.

In general, the condensate environment destabilizes RNA duplexes (Figure 6b-d), in agreement with previous reports.^71,72^ However, the addition of salt reduces this destabilization. For kosmotropic anions, such as F^-^, duplex stability is enhanced both in buffer and inside condensates, likely because it lowers the solubility of the free nucleobases in the single-stranded state. For cations, however, we observe a complete inversion of their effect on duplex stability. While the kosmotropic cations Mg^2+^ and Li^+^ destabilize the duplex in buffer, they favour duplex formation in condensates. According to LMWA, individual charge-charge interactions between RNA and protamine are relatively weak, and therefore the stronger-binding Mg^2+^ and Li^+^ may weaken the interaction with protamine and favour duplex formation. We also observe a significant increase in RNA partitioning in presence of MgCl_2_, which may be due to the positive *ζ*-potential of the droplets (Supplementary Figure 61). We observe similar trends for DNA (Supplementary Figure 62-63), showing that ion binding can fundamentally alter condensate properties, and possibly their function as passive helicases.

## Discussion

Through thorough NMR analysis, we obtained detailed molecular insight into ion binding and uptake in biomolecular condensates. Interactions between condensate components and salt ions follow the law of matching water affinities (LMWA), resulting in strong binding between chaotropic anions and cationic proteins, and between kosmotropic cations and nucleic acids or anionic proteins. This results in the strongest inclusion of kosmotropic cations (Li^+^, Mg^2+^, Ca^2+^ and to a lesser degree Na^+^), and chaotropic anions (ClO_4_^-^, I^-^, and to a lesser degree Br^-^ and Cl^-^), but exclusion of Cs^+^, K^+^ and F^-^. Ion binding shapes the condensate microenvironment by altering the composition, viscosity and interface potential. Such changes can have profound effects on biochemical processes taking place inside the condensates, as we show for RNA duplex formation.

The observation that charge-charge interactions are determined by the hydration strengths of the interacting species prompted us to rethink such interactions in the context of biomolecular condensates. The entropic rearrangement of water molecules during ion complexation plays a key role in charge-charge interactions, and therefore charge-charge interactions should not be considered as purely enthalpic, as is typically done. Specific ion binding may also help to explain the thermodynamic properties of charge-based condensate formation, which is typically entropy-driven^18– 21,73,74^ and sometimes even enthalpically unfavorable^18,21,74^ (see Extended Discussion for more detail; Supplementary Information Section 4).

Our findings also provide further insight into the driving forces behind partitioning of small molecules into condensates, and may explain how condensates can modulate subcellular ion distributions and regulate cellular electrochemistry through interphase potentials.^12,13^ They further show that solvent effects should be explicitly considered when investigating interactions between guest molecules and condensates, for instance in the context of wastewater treatment^15–17^ and delivery of small-molecule therapeutics.^49–51^

## Acknowledgements

The authors would like to thank Brent Visser for help with RICS analysis and setting up the Raman microscopy methods, Merlijn van Haren for help with *ζ*-potential analysis, prof. Paul Cremer for insightful discussions on ion binding, Ben Joosten for help with Raman microscopy, and Paul van der Ven for ICP-MS analysis. The authors acknowledge the European Synchrotron Radiation Facility (ESRF) for provision of synchrotron radiation facilities under proposal number MX-2671 and would like to thank Dr. Stephanie Hutin for assistance and support in using beamline BM29. This project was funded by the European Research Council (ERC) under the European Union’s Horizon 2020 research and innovation programme under grant agreement number 851963. E.L., M.P. and A.B. acknowledge funding from the Swiss National Science Foundation (SNSF) under grant CRSII5_193740 and of the French Agence Nationale de la Recherche (ANR) under grant ANR-21-CE30-0001. M.K. acknowledges funding from the Natural Sciences and Engineering Research Council of Canada (NSERC) RGPIN-2017-05935.

## Author contributions

I.B.A.S., E.S. and M.K. conceived the idea and designed the experiments. I.B.A.S. performed and analysed all NMR, CSC, partitioning, brightfield- and Raman microscopy, RICS, microelectrophoresis and FRET experiments, and constructed all phase diagrams. E.L. and M.P. performed and analysed molecular dynamics simulations. R.V.M.F. and I.K.V. performed and analysed SAXS measurements. P.B.W. helped to set up the NMR methods. I.B.A.S. wrote the original draft. I.B.A.S., E.L., R.V.M.F., I.K.V., M.K. and E.S. reviewed and edited the paper. E.S., A.B. and I.K.V. acquired funding. E.S., M.K., A.B. and I.K.V. supervised the project. All authors have read and agreed to the final version of the paper.

## References

(1) Banani, S. F.; Lee, H. O.; Hyman, A. A.; Rosen, M. K. Biomolecular Condensates: Organizers of Cellular Biochemistry. Nat. Rev. Mol. Cell Biol. 2017, 18 (5), 285–298. 10.1038/nrm.2017.7.

(2) Juliani do Amaral, M.; Mohapatra, S.; Passos, A. R.; Lopes da Silva, T. S.; Carvalho, R. S.; da Silva Almeida, M.; Pinheiro, A. S.; Wegmann, S.; Cordeiro, Y. Copper Drives Prion Protein Phase Separation and Modulates Aggregation. Sci. Adv. 2023, 9 (44), eadi7347. 10.1126/sciadv.adi7347.

(3) Smokers, I. B. A.; Visser, B. S.; Slootbeek, A. D.; Huck, W. T. S.; Spruijt, E. How Droplets Can Accelerate Reactions─Coacervate Protocells as Catalytic Microcompartments. Acc. Chem. Res. 2024, 57 (14), 1885–1895. 10.1021/acs.accounts.4c00114.

(4) O’Flynn, B. G.; Mittag, T. The Role of Liquid–Liquid Phase Separation in Regulating Enzyme Activity. Curr. Opin. Cell Biol. 2021, 69, 70–79. 10.1016/j.ceb.2020.12.012.

(5) Poudyal, R. R.; Guth-Metzler, R. M.; Veenis, A. J.; Frankel, E. A.; Keating, C. D.; Bevilacqua, P. C. Template-Directed RNA Polymerization and Enhanced Ribozyme Catalysis inside Membraneless Compartments Formed by Coacervates. Nat. Commun. 2019, 10 (1), 1–13. 10.1038/s41467-019-08353-4.

(6) Iglesias-Artola, J. M.; Drobot, B.; Kar, M.; Fritsch, A. W.; Mutschler, H.; Dora Tang, T. Y.; Kreysing, M. Charge-Density Reduction Promotes Ribozyme Activity in RNA–Peptide Coacervates via RNA Fluidization and Magnesium Partitioning. Nat. Chem. 2022, 14 (4), 407–416. 10.1038/s41557-022-00890-8.

(7) Koga, S.; Williams, D. S.; Perriman, A. W.; Mann, S. Peptide-Nucleotide Microdroplets as a Step towards a Membrane-Free Protocell Model. Nat. Chem. 2011, 3 (9), 720–724. 10.1038/nchem.1110.

(8) Peeples, W.; Rosen, M. K. Mechanistic Dissection of Increased Enzymatic Rate in a Phase-Separated Compartment. Nat. Chem. Biol. 2021, 17 (6), 693–702. 10.1038/s41589-021-00801-x.

(9) Saha, B.; Chatterjee, A.; Reja, A.; Das, D. Condensates of Short Peptides and ATP for the Temporal Regulation of Cytochrome: C Activity. Chem. Commun. 2019, 55 (94), 14194–14197. 10.1039/c9cc07358b.

(10) Crosby, J.; Treadwell, T.; Hammerton, M.; Vasilakis, K.; Crump, M. P.; Williams, D. S.; Mann, S. Stabilization and Enhanced Reactivity of Actinorhodin Polyketide Synthase Minimal Complex in Polymer–Nucleotide Coacervate Droplets. Chem. Commun. 2012, 48 (97), 11832–11834. 10.1039/c2cc36533b.

(11) Küffner, A. M.; Prodan, M.; Zuccarini, R.; Capasso Palmiero, U.; Faltova, L.; Arosio, P. Acceleration of an Enzymatic Reaction in Liquid Phase Separated Compartments Based on Intrinsically Disordered Protein Domains. ChemSystemsChem 2020, 2 (4), e2000001. 10.1002/syst.202000001.

(12) Dai, Y.; Wang, Z.-G.; Zare, R. N. Unlocking the Electrochemical Functions of Biomolecular Condensates. Nat. Chem. Biol. 2024, 20 (11), 1420–1433. 10.1038/s41589-024-01717-y.

(13) Dai, Y.; Zhou, Z.; Yu, W.; Ma, Y.; Kim, K.; Rivera, N.; Mohammed, J.; Lantelme, E.; Hsu-Kim, H.; Chilkoti, A.; You, L. Biomolecular Condensates Regulate Cellular Electrochemical Equilibria. Cell 2024, 187 (21), 5951–5966.e18. 10.1016/j.cell.2024.08.018.

(14) Posey, A. E.; Bremer, A.; Erkamp, N. A.; Pant, A.; Knowles, T. P. J.; Dai, Y.; Mittag, T.; Pappu, R. V.; Authors, C. Biomolecular Condensates Are Characterized by Interphase Electric Potentials. J. Am. Chem. Soc. 2024, 146, 2024.07.02.601783. 10.1101/2024.07.02.601783.

(15) Bediako, J. K.; Kang, J. H.; Yun, Y. S.; Choi, S. H. Facile Processing of Polyelectrolyte Complexes for Immobilization of Heavy Metal Ions in Wastewater. ACS Appl. Polym. Mater. 2022, 4 (4), 2346–2354. 10.1021/acsapm.1c01634.

(16) Zhou, L.; Shi, H.; Li, Z.; He, C. Recent Advances in Complex Coacervation Design from Macromolecular Assemblies and Emerging Applications. Macromol. Rapid Commun. 2020, 41 (21), 2000149. 10.1002/marc.202000149.

(17) Valley, B.; Jing, B.; Ferreira, M.; Zhu, Y. Rapid and Efficient Coacervate Extraction of Cationic Industrial Dyes from Wastewater. ACS Appl. Mater. Interfaces 2019, 11 (7), 7472–7478. 10.1021/acsami.8b21674.

(18) Chang, L. W.; Lytle, T. K.; Radhakrishna, M.; Madinya, J. J.; Vélez, J.; Sing, C. E.; Perry, S. L. Sequence and Entropy-Based Control of Complex Coacervates. Nat. Commun. 2017, 8 (1), 1–8. 10.1038/s41467-017-01249-1.

(19) Kayitmazer, A. B. Thermodynamics of Complex Coacervation. Adv. Colloid Interface Sci. 2017, 239, 169–177. 10.1016/j.cis.2016.07.006.

(20) Chen, S.; Wang, Z. G. Driving Force and Pathway in Polyelectrolyte Complex Coacervation. Proc. Natl. Acad. Sci. U. S. A. 2022, 119 (36), e2209975119. 10.1073/pnas.2209975119.

(21) Priftis, D.; Laugel, N.; Tirrell, M. Thermodynamic Characterization of Polypeptide Complex Coacervation. Langmuir 2012, 28 (45), 15947–15957. 10.1021/la302729r.

(22) Spruijt, E.; Sprakel, J.; Lemmers, M.; Cohen Stuart, M. A.; Van der Gucht, J. Relaxation Dynamics at Different Time Scales in Electrostatic Complexes: Time-Salt Superposition. Phys. Rev. Lett. 2010, 105, 208301. 10.1103/PhysRevLett.105.208301.

(23) Crabtree, M. D.; Holland, J.; Pillai, A. S.; Kompella, P. S.; Babl, L.; Turner, N. N.; Eaton, J. T.; Hochberg, G. K. A.; Aarts, D. G. A. L.; Redfield, C.; Baldwin, A. J.; Nott, T. J. Ion Binding with Charge Inversion Combined with Screening Modulates DEAD Box Helicase Phase Transitions. Cell Rep. 2023, 42 (11). 10.1016/j.celrep.2023.113375.

(24) Hong, Y.; Najafi, S.; Casey, T.; Shea, J. E.; Han, S. I.; Hwang, D. S. Hydrophobicity of Arginine Leads to Reentrant Liquid-Liquid Phase Separation Behaviors of Arginine-Rich Proteins. Nat. Commun. 2022, 13 (1), 1–15. 10.1038/s41467-022-35001-1.

(25) Guo, Q.; Zou, G.; Qian, X.; Chen, S.; Gao, H.; Yu, J. Hydrogen-Bonds Mediate Liquid-Liquid Phase Separation of Mussel Derived Adhesive Peptides. Nat. Commun. 2022, 13 (1), 1–10. 10.1038/s41467-022-33545-w.

(26) Wei, W.; Petrone, L.; Tan, Y.; Cai, H.; Israelachvili, J. N.; Miserez, A.; Waite, J. H. An Underwater Surface-Drying Peptide Inspired by a Mussel Adhesive Protein. Adv. Funct. Mater. 2016, 26 (20), 3496–3507. 10.1002/adfm.201600210.

(27) Gabryelczyk, B.; Cai, H.; Shi, X.; Sun, Y.; Swinkels, P. J. M.; Salentinig, S.; Pervushin, K.; Miserez, A. Hydrogen Bond Guidance and Aromatic Stacking Drive Liquid-Liquid Phase Separation of Intrinsically Disordered Histidine-Rich Peptides. Nat. Commun. 2019, 10 (1), 1–12. 10.1038/s41467-019-13469-8.

(28) Lim, J.; Kumar, A.; Low, K.; Verma, C. S.; Mu, Y.; Miserez, A.; Pervushin, K. Liquid-Liquid Phase Separation of Short Histidine- And Tyrosine-Rich Peptides: Sequence Specificity and Molecular Topology. J. Phys. Chem. B 2021, 125 (25), 6776–6790. 10.1021/acs.jpcb.0c11476.

(29) Van Haren, M. H. I.; Nakashima, K. K.; Spruijt, E. Coacervate-Based Protocells: Integration of Life-like Properties in a Droplet. J. Syst. Chem. 2020, 8, 107–120.

(30) Slootbeek, A. D.; van Haren, M. H. I.; Smokers, I. B. A.; Spruijt, E. Growth, Replication and Division Enable Evolution of Coacervate Protocells. Chem. Commun. 2022, 58 (80), 11183–11200. 10.1039/d2cc03541c.

(31) Ghosh, B.; Bose, R.; Tang, T. Y. D. Can Coacervation Unify Disparate Hypotheses in the Origin of Cellular Life? Curr. Opin. Colloid Interface Sci. 2021, 52, 101415. 10.1016/j.cocis.2020.101415.

(32) De Ronde, C. E. J.; Channer, D. M. D.; Faure, K.; Bray, C. J.; Spooner, E. T. C. Fluid Chemistry of Archean Seafloor Hydrothermal Vents: Implications for the Composition of circa 3.2 Ga Seawater. Geochim. Cosmochim. Acta 1997, 61 (19), 4025–4042. 10.1016/s0016-7037(97)00205-6.

(33) Izawa, M. R. M.; Nesbitt, H. W.; MacRae, N. D.; Hoffman, E. L. Composition and Evolution of the Early Oceans: Evidence from the Tagish Lake Meteorite. Earth Planet. Sci. Lett. 2010, 298 (3–4), 443–449. 10.1016/j.epsl.2010.08.026.

(34) Friedowitz, S.; Lou, J.; Barker, K. P.; Will, K.; Xia, Y.; Qin, J. Looping-in Complexation and Ion Partitioning in Nonstoichiometric Polyelectrolyte Mixtures. Sci. Adv. 2021, 7 (31). 10.1126/sciadv.abg8654.

(35) Iyer, D.; Syed, V. M. S.; Srivastava, S. Influence of Divalent Ions on Composition and Viscoelasticity of Polyelectrolyte Complexes. J. Polym. Sci. 2021, 59 (22), 2895–2904. 10.1002/pol.20210668.

(36) Schlenoff, J. B.; Yang, M.; Digby, Z. A.; Wang, Q. Ion Content of Polyelectrolyte Complex Coacervates and the Donnan Equilibrium. Macromolecules 2019, 52 (23), 9149–9159. 10.1021/acs.macromol.9b01755.

(37) Li, L.; Srivastava, S.; Andreev, M.; Marciel, A. B.; De Pablo, J. J.; Tirrell, M. V. Phase Behavior and Salt Partitioning in Polyelectrolyte Complex Coacervates. Macromolecules 2018, 51 (8), 2988–2995. 10.1021/acs.macromol.8b00238.

(38) Perry, S. L.; Li, Y.; Priftis, D.; Leon, L.; Tirrell, M. The Effect of Salt on the Complex Coacervation of Vinyl Polyelectrolytes. Polymers (Basel). 2014, 6 (6), 1756–1772. 10.3390/polym6061756.

(39) Shen, Y.; Li, S.; Jiang, J.; Sun, F.; Zhao, Y.; Qiao, F.; Qin, B. Ions Effect on Tunable Coacervate and Its Relevance to the Hofmeister Series. Colloids Surfaces A Physicochem. Eng. Asp. 2024, 702, 134597. 10.1016/j.colsurfa.2024.134597.

(40) Allegri, G.; Huskens, J.; Martinho, R. P.; Lindhoud, S. Distribution of Polyelectrolytes and Counterions upon Polyelectrolyte Complexation. J. Colloid Interface Sci. 2024, 672, 654–663. 10.1016/J.JCIS.2024.06.062.

(41) Duan, C.; Wang, R. A Unified Description of Salt Effects on the Liquid-Liquid Phase Separation of Proteins. ACS Cent. Sci. 2024, 10 (2), 460–468. 10.1021/acscentsci.3c01372.

(42) Frankel, E. A.; Bevilacqua, P. C.; Keating, C. D. Polyamine/Nucleotide Coacervates Provide Strong Compartmentalization of Mg2+, Nucleotides, and RNA. Langmuir 2016, 32 (8), 2041–2049. 10.1021/acs.langmuir.5b04462.

(43) Zhu, L.; Pan, Y.; Hua, Z.; Liu, Y.; Zhang, X. Ionic Effect on the Microenvironment of Biomolecular Condensates. J. Am. Chem. Soc. 2024, 146 (20), 14307–14317. 10.1021/jacs.4c04036.

(44) Ausserwoger, H.; Qian, D.; Krainer, G.; Welsh, T. J.; Sneideris, T.; Franzmann, T. M.; Qamar, S.; Nixon-Abell, J.; Kar, M.; Hyslop, P. S. G.-; Hyman, A. A.; Alberti, S.; Pappu, R. V.; Knowles, T. P. J. Quantifying Collective Interactions in Biomolecular Phase Separation. bioRxiv 2023, 2023.05.31.543137. 10.1101/2023.05.31.543137.

(45) Chen, C.; Yi, R.; Igisu, M.; Sakaguchi, C.; Afrin, R.; Potiszil, C.; Kunihiro, T.; Kobayashi, K.; Nakamura, E.; Ueno, Y.; Antunes, A.; Wang, A.; Chandru, K.; Hao, J.; Jia, T. Z. Spectroscopic and Biophysical Methods to Determine Differential Salt-Uptake by Primitive Membraneless Polyester Microdroplets. Small Methods 2023, 7 (12), 2300119. 10.1002/smtd.202300119.

(46) Neitzel, A. E.; Fang, Y. N.; Yu, B.; Rumyantsev, A. M.; De Pablo, J. J.; Tirrell, M. V. Polyelectrolyte Complex Coacervation across a Broad Range of Charge Densities. Macromolecules 2021, 54 (14), 6878–6890. 10.1021/acs.macromol.1c00703.

(47) Horne, J.; Barker, K.; Xia, Y.; Qin, J. Effects of Divalent Salts on Polyelectrolyte Coacervation. 2024. 10.26434/CHEMRXIV-2024-RS8KX.

(48) Akkaoui, K.; Digby, Z. A.; Do, C.; Schlenoff, J. B. Comprehensive Dynamics in a Polyelectrolyte Complex Coacervate. Macromolecules 2024, 57 (3), 1169–1181. 10.1021/acs.macromol.3c01540.

(49) Kilgore, H. R.; Mikhael, P. G.; Overholt, K. J.; Boija, A.; Hannett, N. M.; Van Dongen, C.; Lee, T. I.; Chang, Y. T.; Barzilay, R.; Young, R. A. Distinct Chemical Environments in Biomolecular Condensates. Nat. Chem. Biol. 2024, 20 (3), 291–301. 10.1038/s41589-023-01432-0.

(50) Kilgore, H. R.; Young, R. A. Learning the Chemical Grammar of Biomolecular Condensates. Nat. Chem. Biol. 2022, 18 (12), 1298–1306. 10.1038/s41589-022-01046-y.

(51) Ambadi Thody, S.; Clements, H. D.; Baniasadi, H.; Lyon, A. S.; Sigman, M. S.; Rosen, M. K. Small-Molecule Properties Define Partitioning into Biomolecular Condensates. Nat. Chem. 2024, 16, 1794–1802. 10.1038/s41557-024-01630-w.

(52) Sing, C. E.; Qin, J. Bridging Field Theory and Ion Pairing in the Modeling of Polyelectrolytes and Complex Coacervation. Macromolecules 2023, 56 (15), 5941–5963. 10.1021/acs.macromol.3c01020.

(53) Lytle, T. K.; Sing, C. E. Transfer Matrix Theory of Polymer Complex Coacervation. Soft Matter 2017, 13 (39), 7001–7012. 10.1039/c7sm01080j.

(54) Kim, H.; Jeon, B. jin; Kim, S.; Jho, Y. S.; Hwang, D. S. Upper Critical Solution Temperature (UCST) Behavior of Coacervate of Cationic Protamine and Multivalent Anions. Polymers (Basel). 2019, 11 (4), 691. 10.3390/polym11040691.

(55) Mountain, G. A.; Keating, C. D. Formation of Multiphase Complex Coacervates and Partitioning of Biomolecules within Them. Biomacromolecules 2020, 21 (2), 630–640. 10.1021/acs.biomac.9b01354.

(56) Lenton, S.; Hervø-Hansen, S.; Popov, A. M.; Tully, M. D.; Lund, M.; Skepö, M. Impact of Arginine-Phosphate Interactions on the Reentrant Condensation of Disordered Proteins. Biomacromolecules 2021, 22 (4), 1532–1544. 10.1021/acs.biomac.0c01765.

(57) Pir Cakmak, F.; Marianelli, A. M.; Keating, C. D. Phospholipid Membrane Formation Templated by Coacervate Droplets. Langmuir 2021, 37 (34), 10366–10375. 10.1021/acs.langmuir.1c01562.

(58) Kota, D.; Prasad, R.; Zhou, H. X. Adenosine Triphosphate Mediates Phase Separation of Disordered Basic Proteins by Bridging Intermolecular Interaction Networks. J. Am. Chem. Soc. 2024, 146 (2), 1326–1336. 10.1021/jacs.3c09134.

(59) Hofmeister, F. Zur Lehre von Der Wirkung Der Salze - Dritte Mittheilung. Arch. für Exp. Pathol. und Pharmakologie 1888, 25 (1), 1–30. 10.1007/BF01838161.

(60) Rembert, K. B.; Paterová, J.; Heyda, J.; Hilty, C.; Jungwirth, P.; Cremer, P. S. Molecular Mechanisms of Ion-Specific Effects on Proteins. J. Am. Chem. Soc. 2012, 134 (24), 10039–10046. 10.1021/ja301297g.

(61) Trujillo, C.; Previtali, V.; Rozas, I. A Theoretical Model of the Interaction between Phosphates in the ATP Molecule and Guanidinium Systems. Theor. Chem. Acc. 2016, 135 (12), 1–12. 10.1007/s00214-016-2012-8.

(62) Collins, K. D. Charge Density-Dependent Strength of Hydration and Biological Structure. Biophys. J. 1997, 72 (1), 65–76. 10.1016/S0006-3495(97)78647-8.

(63) Collins, K. D. The Behavior of Ions in Water Is Controlled by Their Water Affinity. Q. Rev. Biophys. 2019, 52, e11. 10.1017/S0033583519000106.

(64) Jones, G.; Dole, M. The Viscosity of Aqueous Solutions of Strong Electrolytes with Special Reference to Barium Chloride. J. Am. Chem. Soc. 1929, 51 (10), 2950–2964. 10.1021/ja01385a012.

(65) Robinson, J. B.; Strottmann, J. M.; Stellwagen, E. Prediction of Neutral Salt Elution Profiles for Affinity Chromatography. Proc. Natl. Acad. Sci. U. S. A. 1981, 78 (4), 2287–2291. 10.1073/pnas.78.4.2287.

(66) Smokers, I. B. A.; Spruijt, E. Quantification of Biomolecular Condensate Volume Reveals Network Swelling and Dissolution Regimes during Phase Transition. Biomacromolecules 2024. 10.1021/acs.biomac.4c01201.

(67) Hamad, F. G.; Chen, Q.; Colby, R. H. Linear Viscoelasticity and Swelling of Polyelectrolyte Complex Coacervates. Macromolecules 2018, 51 (15), 5547–5555. 10.1021/acs.macromol.8b00401.

(68) Van Haren, M. H. I.; Visser, B. S.; Spruijt, E. Probing the Surface Charge of Condensates Using Microelectrophoresis. Nat. Commun. 2024, 15 (1), 1–10. 10.1038/s41467-024-47885-2.

(69) Smokers, I. B. A.; Visser, B. S.; Lipiñski, W. P.; Nakashima, K. K.; Spruijt, E. Phase-separated Droplets Can Direct Chemical Reaction Kinetics of Polymerization, Self-replication and Oscillating Networks. ChemSystemsChem 2024, e202400056. 10.1002/syst.202400056.

(70) Abbas, M.; Lipiñski, W. P.; Nakashima, K. K.; Huck, W. T. S.; Spruijt, E. A Short Peptide Synthon for Liquid–Liquid Phase Separation. Nat. Chem. 2021, 13 (11), 1046–1054. 10.1038/s41557-021-00788-x.

(71) Cakmak, F. P.; Choi, S.; Meyer, M. C. O.; Bevilacqua, P. C.; Keating, C. D. Prebiotically-Relevant Low Polyion Multivalency Can Improve Functionality of Membraneless Compartments. Nat. Commun. 2020, 11 (1). 10.1038/s41467-020-19775-w.

(72) Nott, T. J.; Craggs, T. D.; Baldwin, A. J. Membraneless Organelles Can Melt Nucleic Acid Duplexes and Act as Biomolecular Filters. Nat. Chem. 2016, 8 (6), 569–575. 10.1038/nchem.2519.

(73) Fu, J.; Schlenoff, J. B. Driving Forces for Oppositely Charged Polyion Association in Aqueous Solutions: Enthalpic, Entropic, but Not Electrostatic. J. Am. Chem. Soc. 2016, 138 (3), 980–990. 10.1021/jacs.5b11878.

(74) Chowdhury, A.; Borgia, A.; Ghosh, S.; Sottini, A.; Mitra, S.; Eapen, R. S.; Borgia, M. B.; Yang, T.; Galvanetto, N.; Ivanović, M. T.; Łukijañczuk, P.; Zhu, R.; Nettels, D.; Kundagrami, A.; Schuler, B. Driving Forces of the Complex Formation between Highly Charged Disordered Proteins. Proc. Natl. Acad. Sci. U. S. A. 2023, 120 (41). 10.1073/pnas.2304036120.

