## Supplementary Information for "Selective ion binding and uptake shape the microenvironment of biomolecular condensates"

3.6. Supplementary data ‘Ion binding changes viscosity and interface potential of condensates’ . 63

### 1. General methods

##### 1.1. Materials

All chemicals and reagents were used as received from commercial suppliers. We used MilliQ (MQ) water (i.e., ultrapure deionized water) from Millipore Corporation.

Protamine chloride grade V, adenosine 5'-triphosphate disodium salt hydrate, poly(A) potassium salt, potassium thiocyanate, potassium thiocyanate-$^{13}$C (99% $^{13}$C), sodium iodide, sodium bromide, sodium chloride, sodium fluoride, sodium formate, sodium acetate, sodium phosphate dibasic anhydrous, calcium chloride dihydrate, magnesium chloride hexahydrate, spermidine trihydrochloride, lithium chloride anhydrous, potassium chloride, cesium chloride, tetramethylammonium chloride, deuterium oxide, hexamethylphosphoramide, phosphonoacetic acid, ethylenediaminetetraacetic acid disodium salt dihydrate and $^{15}$N-ammonium chloride were purchased from Sigma-Aldrich / Merck.

Tris-(hydroxymethyl)-aminomethane (Tris), volumetric 1.0 M hydrochloric acid, volumetric 1.0 M sodium hydroxide, sodium perchlorate, sodium phosphate monobasic anhydrous, sodium sulfate, L(+)-Tartaric acid disodium salt dihydrate and sodium trifluoromethanesulfonate were purchased from Fisher Scientific.

L-Arginine·HCl (guanido-$^{15}N_2$, 98%) and sodium acetate-1,2-$^{13}C_2$ were purchased from Cambridge Isotope Laboratories, Inc. Sodium nitrate and guanidine hydrochloride were purchased from VWR International B.V. Putrescine dihydrochloride was purchased from Bio Connect B.V. 3-(Trimethylsilyl)propionic-2,2,3,3-$d_4$ acid sodium salt was purchased from Deutero GmbH.

Poly-L-lysine hydrobromide (x = 30, MW=6,300 Da) and poly-L-aspartic acid sodium salt (x=30, MW=4,100 Da) were purchased from Alamanda polymers. pLL-*g*-PEG was purchased from SuSoS AG. LLssLL was synthesized according to the procedure in Abbas & Lipiński et al.[1]

Cy3- and Cy5-labelled DNA and RNA oligos for FRET measurements were purchased from Integrated DNA Technologies (IDT).

##### 1.2. Nuclear Magnetic Resonance Spectroscopy – Binding assay

$^{1}$H, $^{15}$N and $^{31}$P-Nuclear Magnetic Resonance (NMR) spectra were measured on a Bruker-AVANCE III 500 MHz spectrometer equipped with a Prodigy BB cryoprobe at 278.15 K for $^{1}$H and $^{31}$P-NMR spectra and at 298.15 K for $^{15}$N-NMR spectra, with a $^{1}$H frequency of 500.13 MHz, a $^{15}$N frequency of 50.68 MHz and a $^{31}$P frequency of 202.46 MHz.

NMR samples were prepared in coaxial NMR tubes with the sample in the outer tube and an internal standard in the inner tube: 3-(trimethylsilyl)propionic-2,2,3,3-$d_4$ acid in $D_2O$ for $^{1}$H-NMR for binding to the α-proton, phosphonoacetic acid in 9 : 1 $H_2O$ : $D_2O$ for $^{1}$H and $^{31}$P-NMR for binding to ATP, and $^{15}$N-ammonium chloride in 9 : 1 $H_2O$ : $D_2O$ for $^{15}$N-NMR for binding to $^{15}$N-labelled arginine. Chemical shifts were determined in reference to the internal standard. All data was processed in MestReNova 14 and Python.

For $^{1}$H-NMR, a solvent suppression sequence was used for the measurement, as a 9 : 1 $H_2O$ : $D_2O$ solvent was used. A pulse sequence was used with 8 or 64 transients (ns = 8 or 64), a P1 pulse length of 12.0 µs which corresponds to approximately a 30° pulse angle and a d1 relaxation delay of 4.0 s. Chemical shifts were determined in reference to the abovementioned internal standards.

For $^{15}N$-NMR, a pulse sequence was set up with 128 transients (ns = 128), a P1 pulse length of 15.0 µs which corresponds to approximately a 90° pulse angle and a d1 relaxation delay of 6 s. Chemical shifts were determined in reference to a $^{15}N$-ammonium chloride internal standard.

For $^{31}P$-NMR, a pulse sequence was set up with 64 transients (ns = 64), a P1 pulse length of 13.0 µs which corresponds to approximately a 90° pulse angle and a d1 relaxation delay of 5 s. Chemical shifts were determined in reference to a hexamethylphosphoramide (HMPA) internal standard.

#### 1.3. Nuclear Magnetic Resonance Spectroscopy – Partitioning

$^{1}H$, $^{7}Li$, $^{13}C$, $^{19}F$, $^{23}Na$, $^{31}P$, $^{35}Cl$ and $^{81}Br$-Nuclear Magnetic Resonance (NMR) spectra were measured on a Bruker-AVANCE III 500 MHz spectrometer equipped with a Prodigy BB cryoprobe at 298.15 K, with a $^{1}H$ frequency of 500.13 MHz, a $^{7}Li$ frequency of 194.37 MHz, a $^{13}C$ frequency of 125.77 MHz, a $^{19}F$ frequency of 470.53 MHz, a $^{23}Na$ frequency of 132.29 MHz, a $^{31}P$ frequency of 202.46 MHz, a $^{35}Cl$ frequency of 49.0 MHz and a $^{81}Br$ frequency of 135.07 MHz. $^{25}Mg$ and $^{39}K$-NMR spectra were measured on a Bruker-AVANCE III 400 MHz HD nanobay spectrometer equipped with a BBFO probe at 298.15 K, with a $^{25}Mg$ frequency of 24.50 MHz and a $^{39}K$ frequency of 18.68 MHz. $^{133}Cs$-NMR spectra were measured on a JEOL JNM-ECZ500R/S3 RoyalHFX 500 MHz spectrometer equipped with an ROHFX probe at 298.15 K, with a $^{133}Cs$ frequency of 65.51 MHz.

Quantitative NMR measurements were performed for all nuclei. Either in reference to an internal standard or to a calibration curve. Relaxation delay d1 values were determined such that all ions and internal standards are fully quantitative (d1 $\geq$ 5x T1). All data was processed in MestReNova 14, and OriginPro 2020b.

For $^{1}H$-NMR, concentrations were determined in reference to a hexamethylphosphoramide (HMPA) internal standard. A solvent suppression sequence was used for the measurement of $^{1}H$-NMR spectra, as a 9 : 1 $H_2O$ : $D_2O$ solvent was used. A pulse sequence was used with 64 transients (ns = 64), a pulse length of 12.0 µs which corresponds to approximately a 30° pulse angle and a d1 relaxation delay of 4.0 s.

For $^{7}Li$-NMR, a pulse sequence was set up with a P1 pulse length of 12.150 µs which corresponds to approximately a 90° pulse angle and a d1 relaxation delay of 200 s, in order to ensure full relaxation of nuclei between each transient. For the dense phase samples 16 transients (ns = 16) were taken with a set receiver gain of rg = 2050. For the dilute phase samples 8 transients (ns = 8) were taken with a set receiver gain of rg = 1030. Concentrations were determined in reference to a calibration curve (Supplementary Figure 14-15), for which the receiver gain and pulse length were kept constant.

For $^{13}C$-NMR, a pulse sequence was set up with a P1 pulse length of 10.5 µs which corresponds to approximately a 90° pulse angle and a d1 relaxation delay of 212 s, in order to ensure full relaxation of nuclei between each transient. For the dense phase samples 64 transients (ns = 64) were taken. For the dilute phase samples 32 transients (ns = 32) were taken. Concentrations were determined in reference to a sodium acetate-1,2-$^{13}C_2$ internal standard.

For $^{19}F$-NMR, a pulse sequence was set up with 128 transients (ns = 128), a P1 pulse length of 15 µs which corresponds to approximately a 30° pulse angle and a d1 relaxation delay of 10 s, in order to ensure full relaxation of nuclei between each transient. O1P was set to -125 ppm. Concentrations were determined in reference to a sodium trifluoromethanesulfonate internal standard.

For $^{23}Na$-NMR, a pulse sequence was set up with 1024 transients (ns = 1024), a P1 pulse length of 20.5 µs which corresponds to approximately a 90° pulse angle and a d1 relaxation delay of 2 s, in order to ensure full relaxation of nuclei between each transient. The receiver gain was set at rg = 1820.

Concentrations were determined in reference to a calibration curve (Supplementary Figure 16-17), for which the receiver gain (rg = 1820) and pulse length were kept constant.

For $^{25}$Mg-NMR, a pulse sequence was set up with a P1 pulse length of 20.0 µs which corresponds to approximately a 90° pulse angle and a d1 relaxation delay of 0.100 s, in order to ensure full relaxation of nuclei between each transient. For the dense phase samples 20480 transients (ns = 20480) were taken. For the dilute phase samples 4096 transients (ns = 4096) were taken. For both the receiver gain was set at rg = 203. Concentrations were determined in reference to a calibration curve (Suppl. Figure 18-19), for which the receiver gain (rg = 203) and pulse length were kept constant.

For $^{31}$P-NMR, a pulse sequence was set up with 64 transients (ns = 64), a P1 pulse length of 13.0 µs which corresponds to approximately a 90° pulse angle and a d1 relaxation delay of 40 s, in order to ensure full relaxation of nuclei between each transient. Concentrations were determined in reference to a hexamethylphosphoramide (HMPA) internal standard.

For $^{35}$Cl-NMR, a pulse sequence was set up with a P1 pulse length of 15.5 µs which corresponds to approximately a 90° pulse angle and a d1 relaxation delay of 2 s, in order to ensure full relaxation of nuclei between each transient.

For $Cl^-$ the following settings were used: For the dense phase samples either 512 transients (ns = 512) were taken with the receiver gain set to rg = 50.8 or 2048 transients (ns = 2048) were taken with the receiver gain set at rg = 64. For the dilute phase samples 512 transients (ns = 512) were taken with the receiver gain set to rg = 50.8. Concentrations were determined in reference to a calibration curve (Supplementary Figure 20-22) for which the receiver gain (rg = 50.8 or 64) and pulse length were kept constant.

For $ClO_4^-$ (and the $Cl^-$ in the $NaClO_4$ samples) the following settings were used: The sweep width was set to 1195 ppm and O1P to 500 ppm. For both the dense and dilute phase samples 1024 transients (ns = 1024) were taken with the receiver gain set to rg = 50.8. Concentrations were determined in reference to a calibration curve (Supplementary Figure 23-24 for $Cl^-$; 25-26 for $ClO_4^-$), for which the receiver gain (rg = 50.8) and pulse length were kept constant.

For $^{39}$K-NMR, a pulse sequence was set up with a P1 pulse length of 25.0 µs which corresponds to approximately a 90° pulse angle and a d1 relaxation delay of 0.100 s, in order to ensure full relaxation of nuclei between each transient. For the dense phase samples 20480 transients (ns = 20480) were taken with the receiver gain set to rg = 203. For the dilute phase samples 4096 transients (ns = 4096) were taken with the receiver gain set to rg = 101. Concentrations were determined in reference to a calibration curve (Supplementary Figure 27-28), for which the receiver gain (rg = 101 or 203) and pulse length were kept constant.

For $^{81}$Br-NMR, a pulse sequence was set up with a P1 pulse length of 12.0 µs which corresponds to approximately a 90° pulse angle and a d1 relaxation delay of 1 s, in order to ensure full relaxation of nuclei between each transient. For the dense phase samples 6144 transients (ns = 6144) were taken with the receiver gain set to rg = 2050. For the dilute phase samples 2048 transients (ns = 2048) were taken with the receiver gain set to rg = 2050. Concentrations were determined in reference to a calibration curve (Supplementary Figure 29-30) for which the receiver gain (rg = 2050) and pulse length were kept constant.

For $^{133}$Cs-NMR, a pulse sequence was set up with 32 transients (ns = 32), a P1 pulse length of 7.5 µs which corresponds to approximately a 90° pulse angle and a d1 relaxation delay of 23 s, in order to ensure full relaxation of nuclei between each transient. The receiver gain was set at rg = 50. Concentrations were determined in reference to a calibration curve (Supplementary Figure 31-32), for which the receiver gain (rg = 50) and pulse length were kept constant.

##### 1.4. Brightfield microscopy

Samples were transferred to 96- or 384-well plates and imaged on an Olympus IX83 fluorescence microscope equipped with a motorized stage (TANGO Märzhäuser) and LED light source (pE-4000 CoolLED). Images were recorded on a 40× universal plan fluorite objective (WD 0.51 mm, NA 0.75, Olympus) and a temperature controlled CMOS camera (Orca-Flash4.0 Hamamatsu).

##### 1.5. pH measurements

The pH of solutions was determined using a Mettler Toledo Five Easy FE20 pH meter equipped with a Hamilton SpinTrode pH probe, and calibrated with Thermo Scientific Orion standard buffers at pH 4.01. pH 7.00 and pH 10.00.

#### 2. Experimental methods

##### 2.1. Design of model system

The condensate system protamine/ATP was chosen because both components are commercially available and cheap, so we could easily make large quantities of condensate sample for the partitioning experiments. We chose the chloride salt of protamine instead of the more commonly used sulfate salt in order to be able to measure the counterions of both protamine and ATP by NMR spectroscopy. The pH of all solutions was maintained at pH 8.5, in order to have both the arginines in protamine and the phosphates of ATP, as well as all tested ions for the partitioning experiments in a single speciation state. The pH of the buffer and stock solutions was adjusted with minimal amounts of hydrochloric acid or sodium hydroxide, since these ions are already present as counterions of the protamine and ATP, and therefore we would not introduce additional types of ions to the system. To investigate the effect of different ions, we used a series of different anions and cations as sodium- and chloride salts, respectively. We selected anions and cations ranging from strongly kosmotropic (strongly hydrated) to strongly chaotropic (weakly hydrated). In the presence of 100 mM (charge-based) of every tested type of ion, condensates were observed by Brightfield microscopy as liquid droplets that fused rapidly (Supplementary Figure 2). Throughout the manuscript, salt ion concentrations are reported as charge-based concentrations, unless mentioned otherwise.

##### 2.2. Condensate formation

To select the most stable protamine/ATP condensate composition, the critical sodium chloride concentration (CSC) of different ratios of protamine : ATP was measured. A composition of 1 mM protamine with 25 mM ATP gave the highest CSC: 585 mM. We assumed that at the maximum CSC, the condensates are charge-neutral.

Condensate emulsions of 1 mM protamine (molecule-based = 21 mM charge-based) and 25 mM ATP (molecule-based = 100 mM charge-based) in 50 mM Tris pH 8.5 were prepared using stock solutions of 4 mM protamine chloride (grade V, Histone free) in 50 mM Tris pH 8.5 and 100 mM adenosine 5'-triphosphate disodium salt hydrate (ATP) in 50 mM Tris pH 8.5. 50 mM Tris pH 8.5 was prepared by dissolving the appropriate amount of Tris-(hydroxymethyl)-aminomethane (Tris) in MQ water and adjusting the pH to 8.5 using 1 M HCl. 4 mM protamine was prepared by dissolving the solid protamine chloride grade V in 50 mM Tris pH 8.5 and adjusting the pH back to 8.5 using 1 M NaOH. The concentration of the protamine stock is molecule-based and based on the typical sequence of protamine H-PRRRRSSSRPIRRRRPRRASRRRRRGGRRRR-OH (MW = 4235.73 g/mol). 100 mM ATP was prepared by dissolving the appropriate amount of adenosine 5'-triphosphate disodium salt hydrate in 50 mM Tris pH 8.5 and adjusting the pH back to 8.5 using 1 M NaOH. Salt stock solutions were similarly prepared in 50 mM Tris pH 8.5 and the pH was corrected back to 8.5 using 1 M NaOH or 1 M HCl.

For a 1 mL condensate sample, 250 µL 4 mM protamine chloride was added to 500 uL of 50 mM Tris pH 8.5 and the appropriate amount of salt, and the solution was pipetted up and down several times. Subsequently, 250 µL 100 mM ATP was added, upon which the solution became turbid. The emulsion was mixed either by vortexing for a few seconds and inverting the tube 3x or by pipetting up and down several times (for volume determination by cell counting tubes).

##### 2.3. NMR assay for ion binding to condensate components

###### 2.3.1. Binding to the α-proton

NMR samples with 1 mM protamine chloride and 0 – 800 mM salt (molecule-based) in $D_2O$ were prepared using stock solutions of 4 mM protamine chloride in $D_2O$, 2 M NaClO4 in $D_2O$, 2 M NaI in $D_2O$, 2 M NaCl in $D_2O$, 800 mM NaF in $D_2O$ and 800 mM $Na_2HPO_4$ in $D_2O$. The pD of the protamine stock solution was corrected to 8.5 using 1 M NaOD. The pD of the salt stock solutions was not adjusted, so no additional salt ions would be introduced. Samples were prepared in $D_2O$ without buffer to minimize the amount of ions in the sample before salt addition. Samples were prepared by mixing 125 µL 4 mM protamine chloride with the appropriate amount of 2 M or 800 mM salt and $D_2O$ was added up to a total volume of 500 µL. The samples were transferred to the outer tube of coaxial NMR tubes. 5 mM (3-(trimethylsilyl)propionic-2,2,3,3-$d_4$ acid in $D_2O$ was added to the inner coaxial tube as internal standard. The pD was determined for samples with different salt concentrations and never varied by more than 0.77 between 0 and 500 mM salt ($Na_2HPO_4$: 0.40, NaF: 0.77, NaCl: 0.37, NaI: 0.50, $NaClO_4$: 0.11). $^1$H-NMR spectra were recorded as described in Supplementary Information Section 1.2, and the changes in NMR chemical shift (Δδ) were determined in reference to the internal standard. Apparent dissociation constants $K_{D,app}$ were calculated by fitting the change in chemical shift as a function of salt concentration to Equation 1:[2]

$$\Delta\delta = -Ac_{\text{ion}} + \frac{\Delta\delta_{\max} c_{\text{ion}}}{K_{\text{D,app}} + c_{\text{ion}}} \qquad \text{(Equation 1)}$$

Because the samples already contained $Cl^-$ from the counterions of protamine (23.1 mM), for NaCl binding, this amount was added to the $K_{D,app}$.

###### 2.3.2. Binding to $^{15}N_2$-guanido-labeled arginine

To determine the strength of charge-charge interaction between different anions and the arginines in protamine, we measured binding to $^{15}$N-guanidino-labelled arginine monomer. NMR samples with 40 mM $^{15}N_2$-guanido-labeled arginine and 0 – 800 mM salt (molecule-based) in 50 mM Tris pH 8.5 in 9 : 1 $H_2O$ : $D_2O$ were prepared using stock solutions of 80 or 160 mM $^{15}$N-arginine in 50 mM Tris pH 8.5 and 2 M NaClO4, NaI and NaCl in 50 mM Tris pH 8.5 and 800 mM NaF and $Na_2HPO_4$ in 50 mM Tris pH 8.5. The pH of the $^{15}$N-arginine stock solution was corrected back to 8.5 using 1 M NaOH. The pH of the salt stock solutions was not adjusted, so no additional salt ions would be introduced. Samples were prepared in Tris buffer, to avoid large changes in pH upon addition of high salt concentrations. Samples were prepared by mixing 125 µL or 250 µL 160 or 80 mM $^{15}N$-arginine with 50 µL $D_2O$ and the appropriate amount of 2 M or 800 mM salt, and 50 mM Tris pH 8.5 was added up to a total volume of 500 µL. The samples were transferred to the outer tube of coaxial NMR tubes. 100 mM $^{15}N$-ammonium chloride in 9 : 1 $H_2O$ : $D_2O$ was added to the inner coaxial tube as internal standard. The pH was determined for samples with different salt concentrations and never varied by more than 0.24 between 0 and 500 mM salt ($Na_2HPO_4$: 0.14, NaF: 0.23, NaCl: 0.18, NaI: 0.03, $NaClO_4$: 0.24). $^{15}N$-NMR spectra were recorded as described in Supplementary Information Section 1.2, at 25 ˚C because no peaks were observed for the guanidinium at 5 ˚C due to slow exchange. Changes in NMR chemical shift (Δδ) were determined in reference to the internal standard. Apparent dissociation constants $K_{D,app}$ were calculated by fitting the change in chemical shift as a function of salt concentration to Equation 1.

Because the samples already contained $Cl^-$ from the counterions of the $^{15}N$-guanidino-labelled arginine and the Tris buffer (61.8 mM), for NaCl binding, this amount was added to the $K_{D,app}$. $F^-$ was deemed non-binding because the error of the fit was larger than the $K_{D,app}$.

2.3.3. Binding to ATP

NMR samples with 25 mM ATP and 0 – 800 mM salt (molecule-based) in 50 mM Tris pH 8.5 in 9 : 1 $H_2O$ : $D_2O$ were prepared using stock solutions of 100 mM ATP in 50 mM Tris pH 8.5 and 2 M $MgCl_2$, LiCl, NaCl, KCl, $(CH_3)_4NCl$ and guandinine HCl in 50 mM Tris pH 8.5. The pH of the ATP stock solution was corrected back to 8.5 using 1 M NaOH. The pH of the salt stock solutions was not adjusted, so no additional salt ions would be introduced. Samples were prepared in Tris buffer, because in $D_2O$ the change in pH upon addition of high salt concentrations was larger than 0.8 pH units. Samples were prepared by mixing 125 µL 100 mM ATP with 50 µL $D_2O$ and the appropriate amount of 2 M salt, and 50 mM Tris pH 8.5 was added up to a total volume of 500 µL. The samples were transferred to the outer tube of coaxial NMR tubes. 5 or 20 mM phosphonoacetic acid in 9 : 1 $H_2O$ : $D_2O$ was added to the inner coaxial tube as internal standard. The pH was determined for samples with different salt concentrations and never varied by more than 0.26 between 0 and 500 mM salt ($MgCl_2$: 0.26, LiCl: 0.15, NaCl: 0.16, KCl: 0.01, $(CH_3)_4NCl$: 0.04, guanidinium HCl: 0.23). $^{31}P$- and $^{1}H$-NMR spectra were recorded as described in Supplementary Information Section 1.2, and the changes in NMR chemical shift (Δδ) were determined in reference to the internal standard. Apparent dissociation constants $K_{D,app}$ were calculated by fitting the change in chemical shift as a function of salt concentration to Equation 1. The resulting $K_{D,app}$'s were grouped for the phosphates, the ribose and the nucleobase. For the ribose, the *g* and *h* H's (Supplementary Figure 8) were not taken into account for the average, because of their close proximity to the phosphates.

Because the samples already contained $Na^+$ from the counterions of ATP and from pH adjustment with NaOH (92.9 mM), for NaCl binding, this amount was added to the $K_{D,app}$.

**2.4. Molecular dynamics simulations**

Molecular Dynamics (MD) simulations were performed using Gromacs 2022[3] paired with the Des-Amber force-field, which was specifically parametrized for protein-protein interactions and included parameters for the ions.[4] tip4p was used as water model. Simulations were run with the leap-frog algorithm, with a time-step of 2 fs. Short-range interactions were treated with a cut-off of 1.0 nm.

Long-range electrostatic interactions were treated with the PME scheme with a cubic interpolation of order 4. Temperature coupling was set at 298 K with the velocity-rescale algorithm[5] and a time constant 0.1 ps. In production runs, the pressure coupling was set at 1 bar with the Parrinello-Rahman algorithm,[6] with a 2 ps time constant and a compressibility of $4.5 \cdot 10^{-5}$ $bar^{-1}$. Each production run was preceded by a system minimization: two equilibrations of 100 ps in the nvt and npt ensemble, respectively.

The simulation boxes contained 1 protamine and water and different concentrations of ions $F^-$, $Cl^-$ or $ClO_4^-$. The total number of ions included the 21 counterions necessary to neutralize the charge of the system, and the ions to produce the following added salt concentrations: 0 mM, 100 mM, 300 mM, 500 mM, 800 mM, 1100 mM. The box shape was a rhombic dodecahedron, which forms a hexagonal lattice with the periodic boundary conditions.

We initially performed simulations with protamine in an unbiased configuration (for Figure 2a-d) and simulated different salt concentrations. For the $Cl^-$ and $ClO_4^-$ we ran a set of three replicas of 150 ns for each salt concentration. For the $F^-$ ions, we ran a set of three replicas at 300 mM added salt.

Interactions between protamine and ions were measured as hydrogen-ion contacts using the Gromacs mindist tool, counting how many ions were localized within a threshold distance of the protamine hydrogens. The threshold distance was chosen from the radial distribution functions (RDFs), including only atoms within the width of the first visible peak of the interaction. For the $Cl^-$ atoms, these distances were 0.3 nm, 0.3 nm and 0.4 nm for the backbone NH, the α-proton and the guanidine terminal NH groups, respectively. For $ClO_4^-$ ions, the distances were 0.28 nm, 0.36 nm and 0.25 nm, and we counted only the closest oxygen to avoid counting the same ion more than once.

To reproduce the NMR binding curves for $Cl^-$ and $ClO_4^-$ from the binding analysis with MD, binding data up to 500 mM added salt was used and fit to a Langmuir isotherm as in Equation 1, without the linear term:

$$P = \frac{\Delta \cdot c_{\text{ion}}}{K_{\text{D,app}} + c_{\text{ion}}} \qquad \text{(Equation 2)}$$

where $P$ is the average number of protamine hydrogen-ion pairs and Δ is the asymptotic occupancy level.

In the unbiased simulations, we observed two distinct conformation populations (Supplementary Figure 12), one in which the protein was folded once onto itself (called closed) and one in which the protein was stretched (called open). To observe exemplary instances of ion binding to specific residues or residue subsets, we extracted one specific configuration from each conformation population and simulated protamine in a fixed open or closed configuration (for Figure 2e-g). To fix the conformation, we used harmonical position restraints on the heavy atoms with a $10^3$ kJ/(mol $nm^2$) force constant. We performed simulations in sets of two replicas of 150 ns of systems with 100 mM and 500 mM added NaCl. Volumetric density maps of ion distributions were measured using the Volmap tool from VMD.

#### 2.5. Small-Angle X-ray Scattering (SAXS) measurements

SAXS measurements were done for samples containing 0 or 1 mM (4.2 mg/mL) protamine and 0, 5, 10, 25, 100, 200 or 400 mM $NaClO_4$ or NaF in 50 mM Tris pH 8.5. Samples were prepared by mixing buffer with the appropriate amount of 1 M $NaClO_4$ or 800 mM NaF in 50 mM Tris pH 8.5. The appropriate amount of 4 mM protamine in 50 mM Tris pH 8.5 was added just before the measurement, after which the samples were vortexed briefly.

SAXS measurements were performed at the BM29 beamline of the European Synchrotron Radiation Facility (ESRF, Grenoble, France), equipped with a Pilatus3 X 2M 2D detector. The sample-to-detector distance was set to 2.87 meters and the beam wavelength to 1.0 Å. To avoid dust or solid contaminants, samples were centrifuged at 10000 rcf for 10 minutes at 4 °C prior to measurements and the supernatant was used. Samples were measured in 10 exposures of 1 second each at room temperature (22 °C), in constant flow via a flow-through quartz capillary set-up to minimize radiation damage. Backgrounds with the same salt concentration as the sample were measured before each sample. The intensity of the integrated background-subtracted data was plotted as a function of the magnitude of the scattering vector $q = 4\pi\lambda^{-1}\sin(\theta/2)$, where λ is the wavelength of the beam and θ the scattering angle. Acquired raw data is available through ESRF (DOI: 10.15151/ESRF-ES-1893861834).

Data was fitted in SASFit software[7] by a generalized gaussian coil form factor,[8] including a hard sphere structure factor to account for repulsion between peptides. This model has been used previously in literature to fit SAXS data of peptides.[9,10] The generalized gaussian coil form factor *P(q)* is described in Equation 3-4.

$$P(q) = I_0\left(\frac{1}{\nu U^{\frac{1}{2\nu}}}\gamma\left(\frac{1}{2\nu},U\right) - \frac{1}{\nu U^{\frac{1}{\nu}}}\gamma\left(\frac{1}{\nu},U\right)\right) \quad \text{(Equation 3)}$$

$$U = (2\nu+1)(2\nu+2)\frac{q^2R_G^2}{6} \quad \text{(Equation 4)}$$

Where $I_0$ is the forward scattering, γ is the lower incomplete Gamma function, $R_g$ is the radius of gyration, and *ν* is the Flory excluded volume parameter. Theoretical values for ν are: ν = ⅓ for collapsed globules (chains in poor solvents), ν = ½ for ideal (Gaussian) chains (in θ-solvents), and ν = ⅗ for swollen chains (in good solvents).

##### 2.6. NMR calibration curves

NMR calibration curves were measured for samples of 0.5, 1, 2, 3, 5, 10, 17.5, 25, 37.5, 50, 100, 150, 200, 250, 300 and 350 mM salt, either in 9 : 1 $H_2O$ : $D_2O$ or in 50 mM Tris pH 8.5 containing 10 % $D_2O$. Receiver gain settings were selected such that there was no saturation of the signal and a linear trend was obtained. The number of scans was selected such that there was signal for all concentrations in the range. Separate calibration curves were prepared for 0 – 350 mM and 0.5 – 5 mM, which were used for the dilute phase and condensate phase samples, respectively. 0.5 – 5 mM samples were typically prepared in duplo or triplo and measurement triplicates were taken.

##### 2.7. Ion partitioning

To determine the protamine, ATP and ion concentration in the dense and dilute phase for different salts, 10 mL samples of 1 mM protamine / 25 mM ATP condensates were prepared with 100 mM of different salts (only for $Na_2HPO_4$ 200 mM charge-based was used), following the procedure in Supplementary Information Section 2.2 using 0.5 M salt stock solutions. The samples were left to equilibrate at room temperature for 20 minutes, after which they were centrifuged for 30 min at 3095 RCF and 20°C. After centrifugation, the supernatant was clear and the condensate was collected as a slightly opaque liquid at the bottom. Most of the dilute phase was removed by micropipette and saved for NMR measurement, making sure that the pipette tip did not touch the condensate phase. The samples were centrifuged again at 3095 RCF and 20°C for 5 minutes to spin down the remaining dilute phase, which was subsequently removed carefully with a micropipette, resulting in an isolated condensate phase. The condensate phase was dissolved by adding 2 - 5 mL 1 M solution of a salt we did not want to measure: 1 M potassium bromide for the partitioning of $Na_2HPO_4$, NaF, NaCl, NaClO4, CsCl, LiCl and $MgCl_2$, 1 M of potassium chloride for NaBr partitioning, 1 M sodium bromide for partitioning of $KS^{13}CN$ and KCl. The condensate phase with added salt was heated to 50°C for 5 minutes, after it was shaken vigorously, and was allowed to cool to room temperature to check if the solution remained clear at room temperature. NMR samples of the dilute phase were prepared by mixing 400 µL dilute phase with 50 µL 1 M potassium bromide (or other salt used for dissolving the condensate phase), 50 µL $D_2O$ and 5 µL 1 M hexamethylphosphoramide (HMPA) internal standard. NMR samples of the condensate phase were prepared by mixing 450 µL of the dissolved condensate phase with 50 µL $D_2O$ and 2 µL 1 M HMPA internal standard. Concentrations of protamine and Tris were determined using $^{1}$H-NMR spectroscopy, the concentration of ATP with $^{31}$P-NMR, and the concentrations of salt ions with $^{7}$Li, $^{13}$C, $^{19}$F, $^{23}$Na, $^{25}$Mg, $^{31}$P, $^{35}$Cl, $^{39}$K, $^{81}$Br and $^{133}$Cs-NMR spectroscopy. All NMR experiments were carried out in triplo. To calculate the dilution factor of the condensate phase upon dissolution and determine the original concentrations in the condensate phase, we used cell counting tubes to determine the total volume of condensate phase in the sample, following the procedure in Smokers & Spruijt (Supplementary Information Section 2.8).[11]

For $MgCl_2$, one condensate sample was prepared for which the condensate phase was dissolved in 2 mL 1 M potassium bromide and heated to 50°C for 5 minutes, which was used to measure the concentration of protamine, ATP, $Na^+$ and $Cl^-$ in the sample. However, because $Mg^{2+}$ is strongly quadrupolar, at neutral pH no signal is observed due to strong binding to ATP. We therefore prepared a sample for which the condensate phase was dissolved in 2 mL 5 M hydrochloric acid and heated to 50°C for 5 minutes to hydrolyze the ATP and neutralize the charges on the phosphates. This sample was used to measure the concentration of $Mg^{2+}$ in the condensate phase. To measure the concentration of $Mg^{2+}$ in the dilute phase, NMR samples were prepared by mixing 400 µL dilute phase with 50 µL 5 M hydrochloric acid and 50 µL $D_2O$.

The partition coefficient of $Mg^{2+}$ was verified by ICP-MS to be $K_P$ = 17.9 (± 1.3), in close agreement with the value found by $^{25}$Mg-NMR. ICP-MS samples were prepared in triplo for 100x dilution and 1000x dilution. Samples of the dilute phase and dense phase dissolved in HCl were diluted 100x and 1000x in MQ with 1% nitric acid and analyzed on a Thermo Scientific iCAP RQ ICP-MS with a limit-of-detection of 0.7 ng/L for $^{24}$Mg and concentrations were determined in reference to a 103Rh internal standard and calibration curve.

To relate the partitioning behavior of the monovalent ions to their water affinity, we made use of the Jones-Dole viscosity *B* coefficient:[12]

$$\frac{\eta}{\eta_0} = 1 + A\sqrt{c} + Bc \qquad \text{(Equation 5)}$$

where $\eta$ is the viscosity of an aqueous salt solution, $\eta_0$ the viscosity of water, $c$ the concentration of salt, $A$ an electrostatic term, and $B$ the Jones-Dole coefficient. $B$ is a measure for the strength of ion-water interactions, and is normalized to the strength of water-water interactions.[13,14] This parameter is an easily interpretable measure for the water affinity of ions: when $B$ = 0, the ion-water interaction strength is equal to that of water-water interactions; when $B$ > 0, the ion-water interactions are stronger than water-water, meaning that the ion is strongly hydrated; when $B$ < 0, the ion-water interactions are weaker than water-water, meaning that the ion is weakly hydrated. Therefore, $B$ = 0 is the transition point between weak hydration (chaotropes) and strong hydration (kosmotropes).

#### 2.8. Condensate volume fraction determination by cell counting tubes

PCV cell counting tubes (capillary graduations only, no cap, Sigma-Adrich) were used directly without surface modification. 1 mL condensate samples were prepared directly in cell counting tubes following the preparation method above, after which the emulsion was mixed by pipetting up and down several times. The samples were centrifuged for 30 min at 3100 RCF and 20°C directly after preparation. Condensate volume was read out from the graduations (Supplementary Figure 1). All experiments were carried out in triplo.

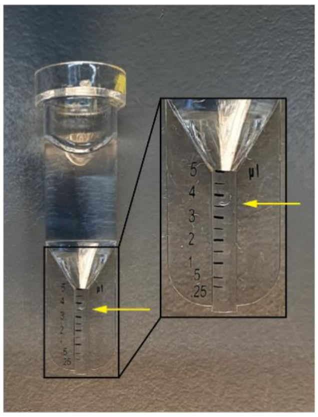

**Supplementary Figure 1:** Read-out of condensate volume fraction from a cell counting tube. Figure reused from ref[11].

#### 2.9. Raman microscopy of $SCN^-$ and $Formate^-$ concentration in individual condensate droplets

Condensate samples for Raman microscopy were prepared as described in Supplementary Information Section 2.2 and added to Ibidi µ-Slide $I^{0.2}$ Luer channel slides, which were modified in advance by cleaning them in a plasma cleaner, modifying for 24 hours using 0.01 mg/mL PLL-*g*-PEG (SuSoS AG) in MQ and subsequently cleaning with MQ and drying with pressurized air. The samples were left to equilibrate and settle on the bottom of the slide for half an hour, after which the slide was inverted and the suspended condensates were visualized using a WITec alpha 300R microscope equipped with a 457 nm laser and Zeiss EC Epiplan-Neofluar 100x/0.9 DIC M27 objective. A 15 mW laser power, 600 g/mm grating (BLZ 500.0 nm) and 10 s integration time were used to obtain per-pixel Raman spectra of a 20x20 µm area (pixel size 1x1 µm). The measurement was performed at the z-height where the protamine and ATP C-H signal ($\tilde{\nu}$ = 2950 $cm^{-1}$) was maximal, i.e. the slice with maximum signal form the condensate. At the start of each measurement day, the Raman signal was calibrated using a $SiO_4$ sample ($\tilde{\nu}$ = 520.7 $cm^{-1}$).

Data was prepared for analysis using the WiTec Project 6.1 software. Cosmic rays were removed using a median filter size (spectral) of 8 and background subtraction was performed using a shape size of 1000. The resulting data was further analyzed using a Python script that fit the signal at $\tilde{\nu}$ = 2064 $cm^{-1}$ (for $SCN^-$) with a Gaussian curve and plotted the integral of the signal per pixel.

#### 2.10. Phase diagrams for NaF, NaCl, $NaClO_4$ and LiCl

To determine the composition of condensates at different total salt concentrations and make phase diagrams, condensate samples were prepared following the procedure in Supplementary Information Section 2.2 with 0 – 300 mM LiCl, 0 – 250 mM NaF, 0 – 400 mM NaCl or 0 – 1 M $NaClO_4$. Phase diagrams were measured using $^{1}H$, $^{7}Li$, $^{19}F$, $^{23}Na$, $^{31}P$ and $^{35}Cl$-NMR, for which samples were prepared following the procedure in Supplementary Information Section 2.7. Instead of 10 mL total volume, 2 mL was used. The condensate phase was isolated by removing most of the supernatant by micropipette and subsequently taking up the remaining dilute phase by lightly touching it with filter paper. The isolated condensate phase was dissolved in either 1 mL (for volume fractions > 0.1%) or 0.5 mL (for volume fractions < 0.1%) 1 M KBr.

##### 2.11. Protocol CSC homotypic

The critical salt concentration (CSC) for homotypic protamine / salt condensates was determined for different salts using a plate reader turbidity assay. Samples of 1 mM protamine and different concentrations of salts in 50 mM Tris pH 8.5 with a total volume of 100 µL were prepared by mixing 50 mM Tris pH 8.5 and different amounts of salt stock solution and adding 25 µL 4 mM protamine. Stock solutions of 4 mM protamine, 4 M NaCl, 5 M NaBr, 4 M KSCN and 5 M $NaClO_4$ were prepared in 50 mM Tris pH 8.5 and their pH was corrected back to 8.5 using 1 M NaOH or 1 M HCl. Samples were vortexed and transferred to a transparent 96-well plate. The absorbance at 600 nm was measured in a Tecan Spark M10 plate reader and the turbidity was calculated as (100 – T%), with T% being the fraction of transmitted light at 600 nm, calculated as:

$$T_{\%} = \frac{I}{I_0} \cdot 100\,\% = 10^{-Abs} \cdot 100\,\% \qquad \text{(Equation 6)}$$

Where Abs is the measured absorbance at 600 nm. The CSC for homotypic protamine / salt condensate formation was determined by plotting the recorded turbidity (100 – T%) as a function of the salt concentration in the sample, and taking the intersect of the tangent lines to the highest and lowest first order derivative of the turbidity, calculated by a MATLAB script. For NaCl the threshold concentration could not be measured exactly due to limited solubility of the salt.

##### 2.12. Determination of protamine/ATP UCST and LLssLL LCST

To determine the upper critical solution temperature (UCST) of protamine/ATP condensates in presence of 100 mM of different salts, a UV-Vis turbidity assay as a function of temperature was used. Condensate samples of 120 µL were prepared as detailed in Supplementary Information Section 2.2 directly in a Hellma Analytics High Precision Cell quartz cuvette with an optical path length of 10x2 mm and center height of 15 mm, suitable for sample volumes of 100 µL. A JASCO V-630 UV-Vis spectrophotometer equipped with Peltier controlled cell holder was used measure the loss in absorbance at 600 nm as a function of temperature from 20 to 70 ˚C at a ramp rate of 1.0 ˚C per minute. The turbidity was calculated as (100 – T%), with T% being the fraction of transmitted light at 600 nm, calculated using Equation 6. The UCST for protamine/ATP condensates was determined by plotting the recorded turbidity (100 – T%) as a function of the temperature in the sample, and taking the intersect of the tangent lines to the highest and lowest first order derivative of the turbidity, calculated by a MATLAB script.

A similar procedure was used to determine the LCST of LLssLL condensates. Samples of 3 mg/mL LLssLL (TFA salt) with 100 mM salt in 50 mM Tris pH 8.5 were prepared by mixing 60 µL 100 mM Tris pH 8.5, 24 µL 0.5 M salt and 36 µL 10 mg/mL LLssLL. The 10 mg/mL LLssLL stock was prepared in MQ water. At room temperature, these samples were clear, but they become turbid upon heating. The appearance of turbidity as a function of temperature was measured using the same method described above, and the LCST was determined using the MATLAB script described above.

##### 2.13. Determination of critical polyion concentration of condensates

The critical polyion concentration of condensates was assessed using a plate reader turbidity assay. 100 µL solution of a polyion with 100 mM salt (charge-based; molecule-based: 50 mM for divalent ions, 40 mM for spermidine) in 50 mM Tris buffer pH 8.5 were added to 96-well plates (Greiner Bio-one, clear flat bottom) and absorbance at 600 nm was measured on a microplate reader (Tecan Spark M10) equipped with an automated injector. After every readout, 2 µL solution of the oppositely charged polyion was added and the samples were mixed by shaking for 5 s. A total of 40 readouts was made, and all measurements were made in triplo. The following pairs of polyions were tested:

**Supplementary Table 1:** Stock solutions used to determine the critical polyion concentration for condensate formation ($[\text{polyion}]_{crit}$) by plate reader assay.

| Polyion in sample | Polyion stock added in titration |
|---|---|
| 0.5 mM protamine | 5 mM ATP for monovalent ions, sodium sulphate, sodium tartrate, sodium phosphate; 10 mM ATP for putrescine HCl; 25 mM for calcium chloride, spermidine; 100 mM for magnesium chloride |
| 0.5 mM polyA | 75 µM protamine |
| 1 mM protamine | 5 mM $D_{30}$ |
| 0.5 mM $D_{30}$ | 0.2 mM protamine |
| 4 mM $K_{30}$ | 5 mM $D_{30}$ |
| 3 mM $D_{30}$ | 10 mM $K_{30}$ |

Turbidity was calculated as (100 – T%), with T% being the fraction of transmitted light at 600 nm, calculated using Equation 6. The critical polyion concentration was determined by plotting the recorded turbidity (100 – T%) as a function of the added polyion concentration in the sample, and taking the intersect of the tangent lines to the highest and lowest first order derivative of the turbidity, calculated by a MATLAB script.

##### 2.14. Viscosity determination using Raster Image Correlation Spectroscopy (RICS)

Condensate samples for RICS were prepared as described in Supplementary Information Section 2.2 with 100 or 200 mM salt (charge-based) and 0.1 mg/mL 4.4 kDa TAMRA-labelled dextran added before addition of protamine and ATP. The samples were added to Ibidi 18-well chambered µ-slides with glass bottom, which were modified in advance by cleaning them in a plasma cleaner, modifying for 24 hours using 0.01 mg/mL PLL-*g*-PEG (SuSoS AG) in MQ and subsequently cleaning with MQ and drying with pressurized air.

The viscosity of condensates was determined using Raster Image Correlation Spectroscopy (RICS) on a Leica SP8 confocal microscope equipped with a single-photon detector. Calibration of the focal volume waist $\omega_0$ was performed using the known diffusion coefficient of Alexa 488 of 435 $\mu m^2\ s^{-1}$ (T = 22.5 ± 0.5 °C) in water, and $\omega_z$ was set to 3 times the value of $\omega_0$.[15] All measurements were captured at a resolution of 256 × 256 pixels with a 20 nm pixel size using a 63x objective. Condensate samples were measured at 10 Hz line speed with 15 frames acquired per data point. Analysis of autocorrelation curves was done using PAM.[16] Measured diffusion coefficients and respective viscosities for each sample are shown in Supplementary Table 24.

##### 2.15. ζ-potential determination by microelectrophoresis

Condensate samples for ζ-potential determination were prepared as described in Supplementary Information Section 2.2 with 100 or 200 mM salt (charge-based). All samples were imaged on Ibidi 6-well µ-channel slides VI 0.4 with bioinert surface modification. Before image acquisition, 100 µL condensate suspensions were transferred to the channels and incubated for 30 minutes to allow droplets to coalesce and settle on the glass surface. Electrodes (2 mm, silver) connected with copper wires to a BT-305A PSU direct current power source (Basetech) were lowered into opposing ends of the microchannel slide and an electric field of 5 to 12 V/cm was applied, with the cathode at the top of the field of view. Moving condensates were imaged in the middle of the channel of the microslide. Samples were imaged on an Olympus IX83 inverted fluorescence microscope equipped with a motorized stage (TANGO, Märzhäuser) and LED light source (pE-4000 CoolLED). Images were recorded with a 40× universal plan fluorite objective (WD 0.51 mm, NA 0.75, Olympus) with a temperature-controlled CMOS camera (Hamamatsu Orca-Flash 4.0).

Raw microscopy videos were processed and analyzed with MATLAB 2021 Image processing Toolbox and droplet trajectories were determined using the method described in Van Haren et al.[17] ζ-potentials for all detected droplets in a sample were determined from their velocities with a modified Smoluchowski equation, using the applied electric field strength, Debye length calculated from salt concentration and the droplet viscosity determined by raster image correlation spectroscopy (RICS). All parameters used to calculate the condensate ζ-potential are available in Supplementary Table 25.

##### 2.16. FRET microscopy analysis of RNA and DNA duplex stability

###### 2.16.1. Preparation of FRET pair oligo stock solutions

For FRET measurements on duplex stability, DNA and RNA decamer complementary strands were used which were labeled with Cy3 on the forward strand and Cy5 on the reverse strand (Supplementary Table 2). DNA and RNA oligo stock solutions were prepared by dissolving the oligos in TE buffer (10 mM Tris pH 8.0, 1 mM EDTA) up to final concentrations of 0.1 mM for the Cy3- and Cy5-labeled oligos and of 1 mM for the unlabeled oligos. Upon addition of TE buffer the samples were vortexed, left for 15 minutes and vortexed again, after which they were aliquoted and stored at -20 ˚C (DNA) or -80 ˚C (RNA) until used. A fresh stock was used for every day of experiments.

**Supplementary Table 2:** DNA and RNA sequences used for FRET analysis of duplex stability.

| **Name** | **Sequence (5’ – 3’)** |
|---|---|
| Cy3-DNA (forward) | ACCTTGTTCC-Cy3 |
| Cy5-DNA (reverse) | Cy5-GGAACAAGGT |
| Cy3-RNA (forward) | ACCUUGUUCC-Cy3 |
| Cy5-RNA (reverse) | Cy5-GGAACAAGGU |
| DNA forward | ACCTTGTTCC |
| DNA reverse | GGAACAAGGT |
| RNA forward | ACCUUGUUCC |
| RNA reverse | GGAACAAGGU |

2.16.2. Sample preparation for FRET measurements

For FRET measurements, condensate samples containing 0.1 µM of both the Cy3- and Cy5-oligo, 50 mM Tris pH 8.5, 15 mM NaCl, 0.5 mM $MgCl_2$, 100 mM (charge-based) of added salt of interest, 1 mM protamine and 25 mM ATP were made. All stock solutions except for the oligo's were prepared in 50 mM Tris pH 8.5. Samples were prepared by mixing Tris buffer, NaCl, $MgCl_2$, the salt of interest and the oligos, after which they were vortexed and centrifuged briefly, heated to 90˚ C for two minutes and left to cool down to room temperature for at least half an hour. Protamine and ATP were added and the samples were vortexed briefly before being transferred to the microscopy slide. For each salt of interest, FRET samples containing both Cy3- and Cy5-labelled DNA or RNA were measured in triplo, and two control samples were measured to determine the FRET correction parameters α and β: one with the Cy3-labelled forward strand and non-labelled reverse strand (donor only), and one with the Cy5-labelled reverse strand and non-labelled forward strand (acceptor only).

Control samples without condensates were prepared in a similar way, but the protamine and ATP were omitted.

2.16.3. FRET measurement by confocal microscopy

Condensate samples were imaged in Ibidi 18-well chambered µ-slides with glass bottom, which were modified in advance by cleaning them in a plasma cleaner, modifying for 24 hours using 0.01 mg/mL PLL-*g*-PEG (SuSoS AG) in MQ and subsequently cleaning with MQ and drying with pressurized air. Before image acquisition, 50 µL condensate suspension was transferred to the well and incubated for at least 5 minutes to allow droplets to settle on the surface. Samples without condensates were imaged in Ibidi 6-well µ-channel slides VI 0.4 (No. 1.5 polymer coverslip) without surface modification. In all cases, triplicate samples and controls were added to separate coverslips to avoid heating during measurement of a neighboring sample.

FRET efficiencies of dsRNA or dsDNA with a Cy3 / Cy5 FRET pair were determined inside condensates using a Leica Sp8x laser scanning confocal microscope equipped with a white-light laser and a 100x magnification oil-immersion objective and the whole setup was incubated at 29 – 30 ˚C. Images were recorded at a line scanning speed of 50 Hz, zoom factor 4.0, a pinhole of 3.0, without averaging or accumulation and saved in 16-bit 256x256 pixels format.

FRET efficiencies in absence of condensates were determined in a similar way, but using a 10x magnification objective, higher laser power, a line scanning speed of 10 Hz, zoom factor 1.00, a pinhole of 3.0, without averaging or accumulation and images were saved in 16-bit 16x16 pixels format. To find the right z-position for the samples, the edge of the channel was put in focus, and this height was used to measure the FRET efficiency inside the channel.

The Cy3 donor was excited at 514 nm and its emission was collected between 555 – 620 nm. FRET emission from the Cy5 acceptor was collected between 650 – 750 nm. The Cy5 acceptor was excited at 633 nm and its emission collected between 650 – 750 nm. Three fluorescence channels were recorded for each image:

$DD_{obs}$= observed Cy3 donor emission after Cy3 donor excitation

$DA_{obs}$= observed Cy5 acceptor emission after Cy3 donor excitation (FRET)

$AA_{obs}$= observed Cy5 acceptor emission after Cy5 acceptor excitation

To correct for the overlap between the emission and excitation wavelengths of the donor and acceptor, the DD, DA and AA measurements described above were also performed for samples containing only donor or only acceptor, with a non-labeled complementary nucleic acid strand and equivalent hardware settings as described above, from which $DD_{donor}$, $DA_{donor}$, $DA_{acceptor}$ and $AA_{acceptor}$ were obtained.

ImageJ was used to quantify the fluorescence intensities from the obtained images. Circular regions of interest (ROIs) were selected inside three different droplets per image, and for each ROI the average DD, DA and AA signal was quantified. The corrected FRET intensity was then calculated according to Equation 7:

$$\mathrm{E}_{CT} = \frac{\mathrm{DA}}{\mathrm{DA+DD}} = \frac{\mathrm{DA_{obs}} - \alpha \cdot \mathrm{DD_{obs}} - \beta \cdot \mathrm{AA_{obs}}}{\mathrm{DD_{obs}} + (\mathrm{DA_{obs}} - \alpha \cdot \mathrm{DD_{obs}} - \beta \cdot \mathrm{AA_{obs}})} \qquad \text{(Equation 7)}$$

With the correction terms calculated according to Equation 8 and 9:

$$\alpha = \frac{\mathrm{DA_{donor}}}{\mathrm{DD_{donor}}} \quad \text{and} \quad \beta = \frac{\mathrm{DA_{accept}}}{\mathrm{AA_{accept}}} \qquad \text{(Equation 8 \& 9)}$$

Correction terms were determined for three ROIs in three images of single measurements. Using the average correction terms, the corrected FRET efficiency was determined per ROI in the images of samples with both donor and acceptor. For each salt of interest, samples were prepared in triplicate and for each replicate three images were recorded, for which three ROIs were measured, resulting in 27 measurements per salt of interest.

### 3. Supplementary Data

##### 3.1. General supplementary data

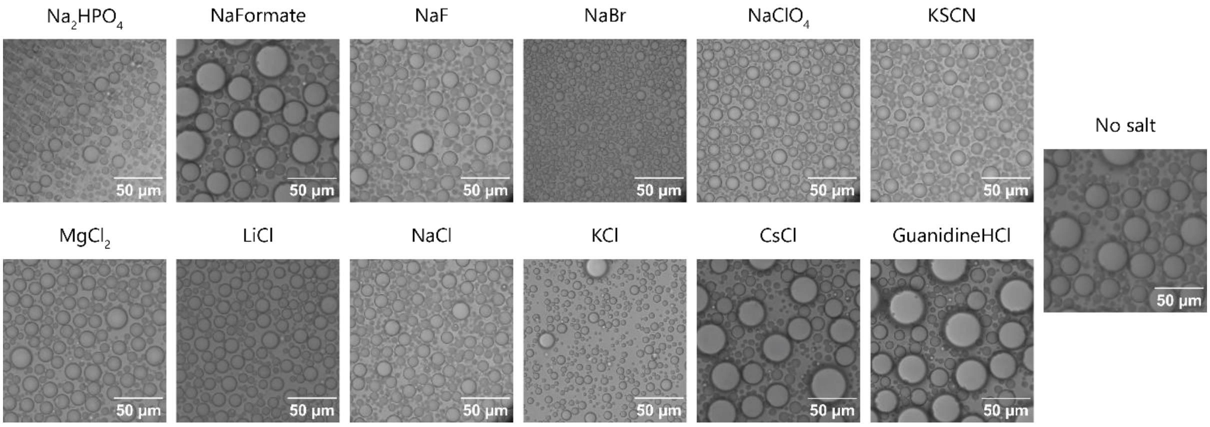

**Supplementary Figure 2:** Brightfield microscopy images of 1 mM protamine / 25 mM ATP condensates in presence of 100 mM (charge-based) of different salts and in absence of salt show that in all cases the condensates are liquid droplets that fuse rapidly.

##### 3.2. Supplementary data 'Selective binding of ions to condensate components'

###### 3.2.1. Binding to the α-proton

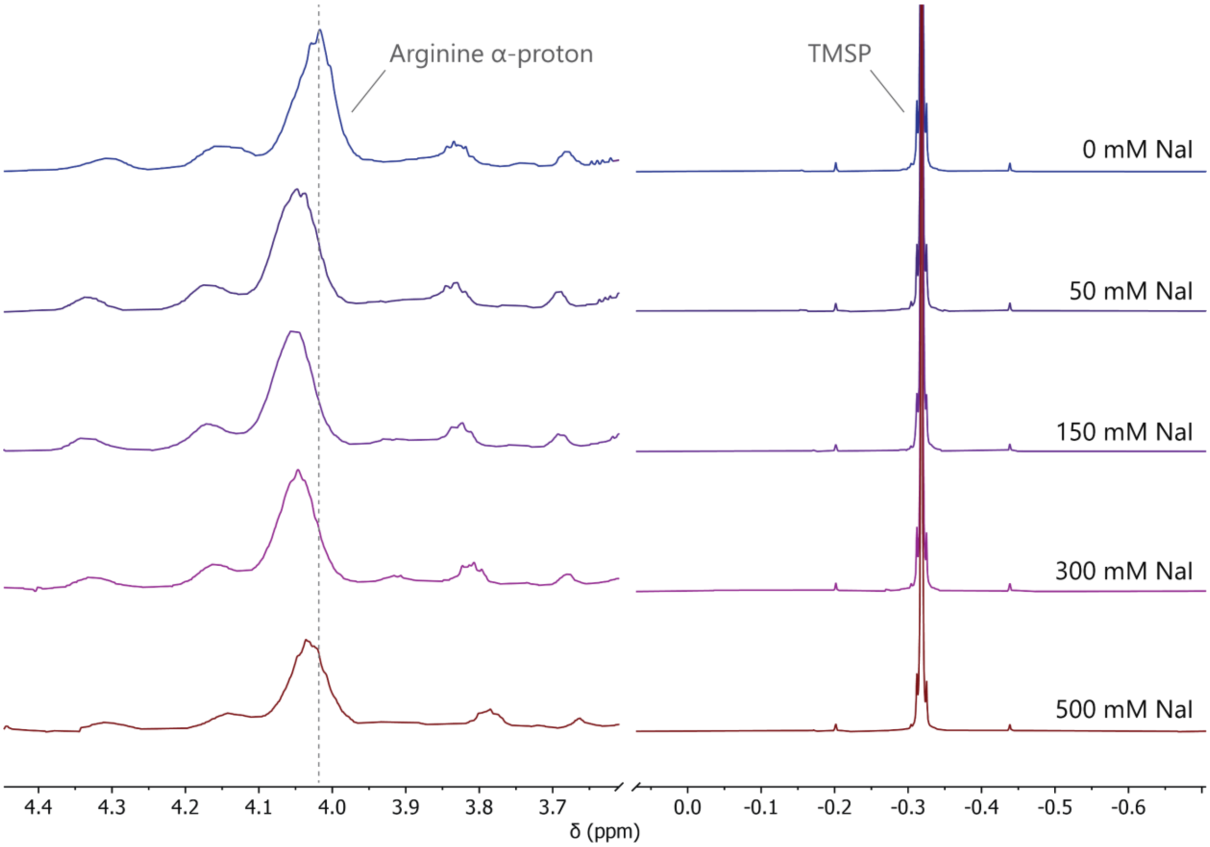

**Supplementary Figure 3:** [1]H-NMR-spectra for binding of NaI to the α-proton of the arginines on 1 mM protamine in $D_2O$ pD 8.5 with 5 mM (3-(trimethylsilyl)propionic-2,2,3,3-$d_4$ acid (TMSP) in $D_2O$ as internal standard in the inner tube of an NMR coaxial tube, measured at 5˚C. Full NMR spectra and spectra for other salts can be found on the Radboud Data Repository (Section 5).

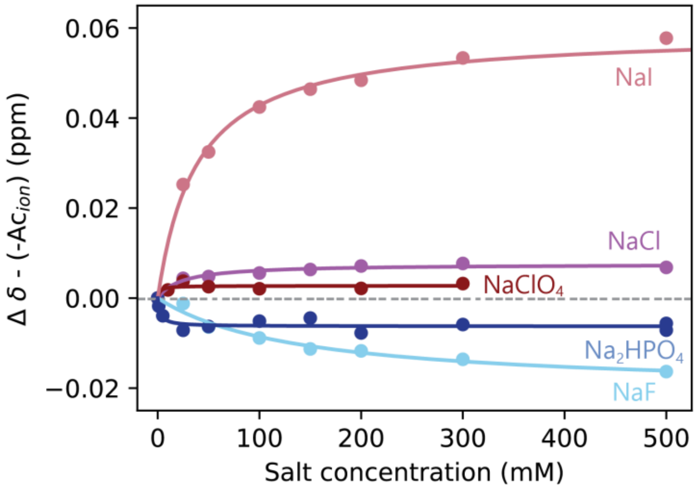

**Supplementary Figure 4:** Fitted binding curves for the non-linear part $\Delta\delta - (-Ac_{\text{ion}}) = \frac{\Delta\delta_{\max}\, c_{\text{ion}}}{K_{\text{D,app}} + c_{\text{ion}}}$ for binding of different anions to the α-proton of the arginines on 1 mM protamine at 5˚C. The difference in sign of $\Delta\delta$-($-Ac_{ion}$) between the chaotropes and kosmotropes likely indicates a different mechanism of binding. The magnitude $\Delta\delta_{max}$ likely correlates to the saturation / number of binding sites, which may be lower for the large $ClO_4^-$ than for the smaller halides.

#### 3.2.2. Binding to $^{15}N_2$-guanido-labeled arginine

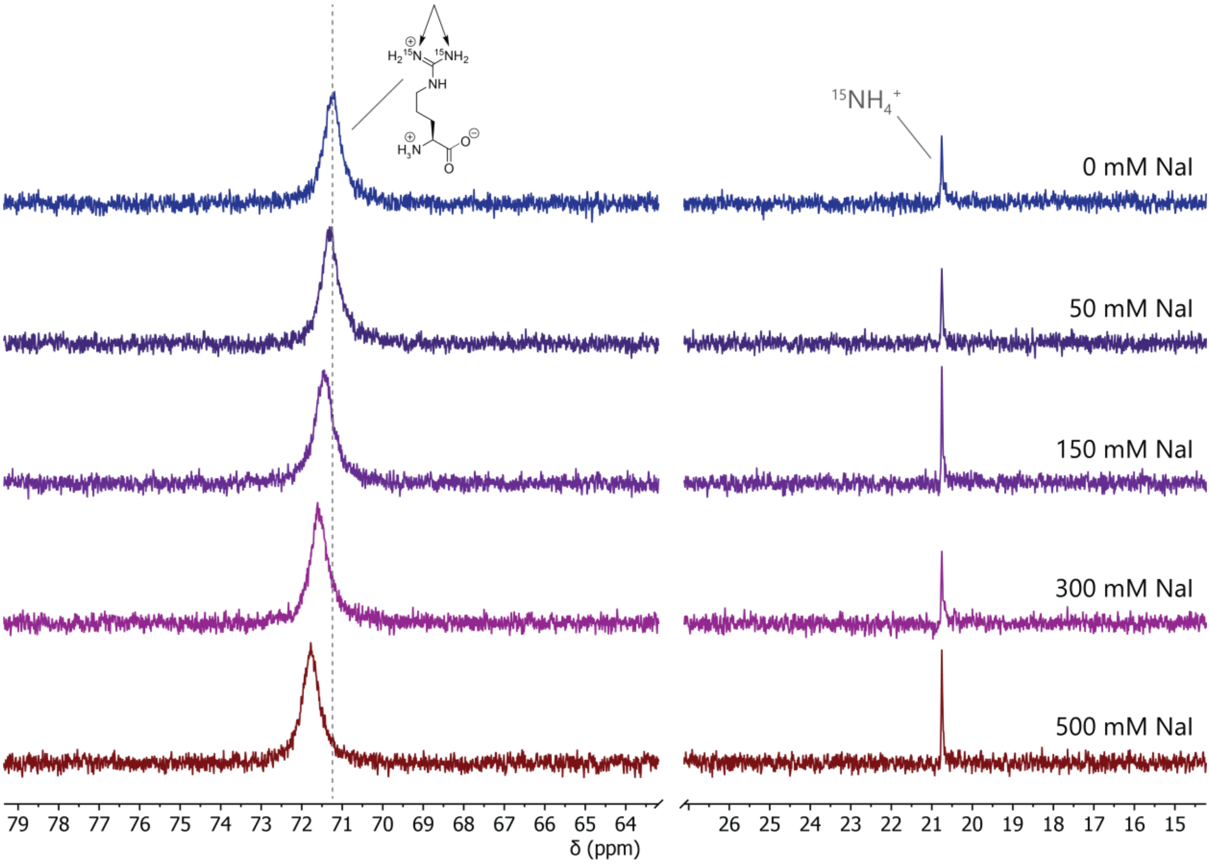

**Supplementary Figure 5:** $^{15}$N-NMR-spectra for binding of NaI to the guanidinium nitrogens of $^{15}N_2$-guanido-labeled arginine in 50 mM Tris pH 8.5 in 9 : 1 $H_2O$ : $D_2O$, with 100 mM $^{15}$N-ammonium chloride in 9 : 1 $H_2O$ : $D_2O$ as internal standard in the inner tube of an NMR coaxial tube, measured at 25˚C. Full NMR spectra and spectra for other salts can be found on the Radboud Data Repository (Section 5).

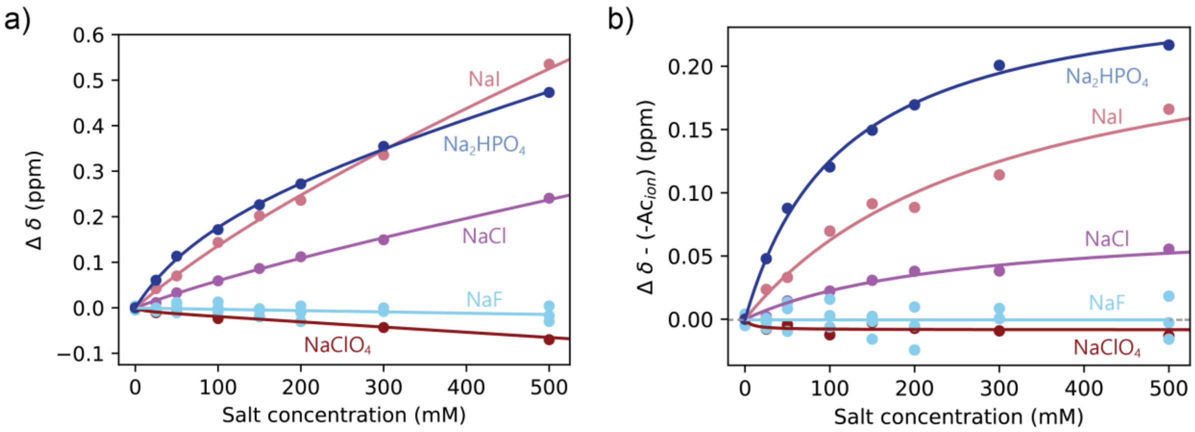

**Supplementary Figure 6: a)** Fitted binding curves for binding of sodium salts of different anions to the guanidinium nitrogens of $^{15}N_2$-guanido-labeled arginine at 25˚C. **b)** Fitted binding curves for the non-linear part $\Delta\delta - (-Ac_{\text{ion}}) = \frac{\Delta\delta_{\max} c_{\text{ion}}}{K_{\text{D,app}} + c_{\text{ion}}}$ for binding of different anions to the guanidinium nitrogens of $^{15}N_2$-guanido-labeled arginine at 25˚C. The difference in sign of Δδ-(-A$c_{ion}$) between different ions likely indicates a different mechanism of binding.

##### 3.2.3. Binding to ATP

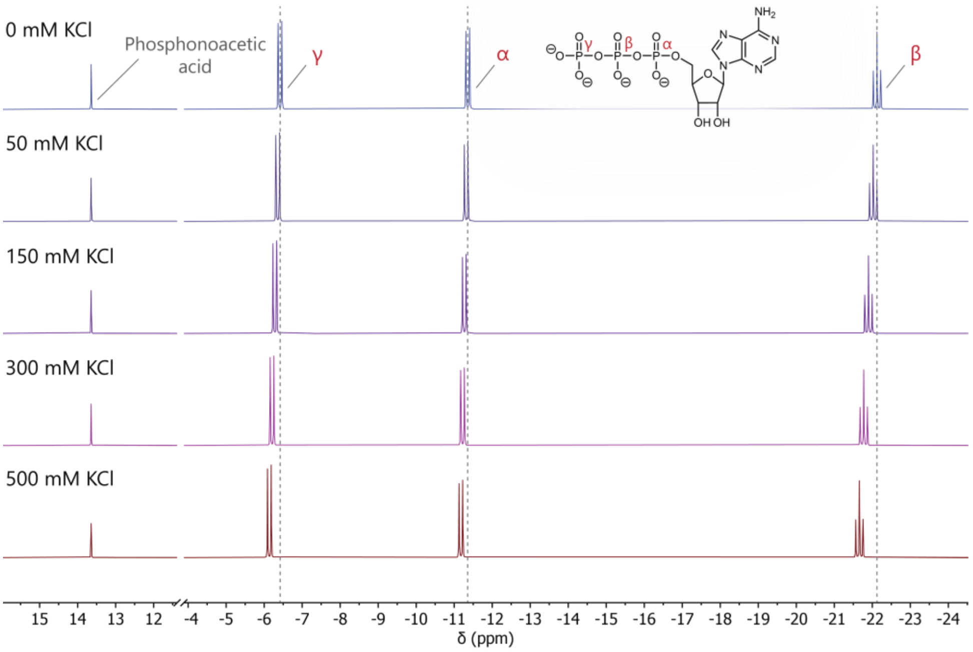

**Supplementary Figure 7:** [31]P-NMR-spectra for binding of KCl to the phosphates on 25 mM ATP in 50 mM Tris pH 8.5 in 9 : 1 $H_2O$ : $D_2O$, with 20 mM phosphonoacetic acid in 9 : 1 $H_2O$ : $D_2O$ as internal standard in the inner tube of an NMR coaxial tube, measured at 5˚C. Full NMR spectra and spectra for other salts can be found on the Radboud Data Repository (Section 5). In the analysis for Figure 1g, the $K_{D,app}$'s of the α-, β- and γ-phosphate were averaged for binding to the phosphates.

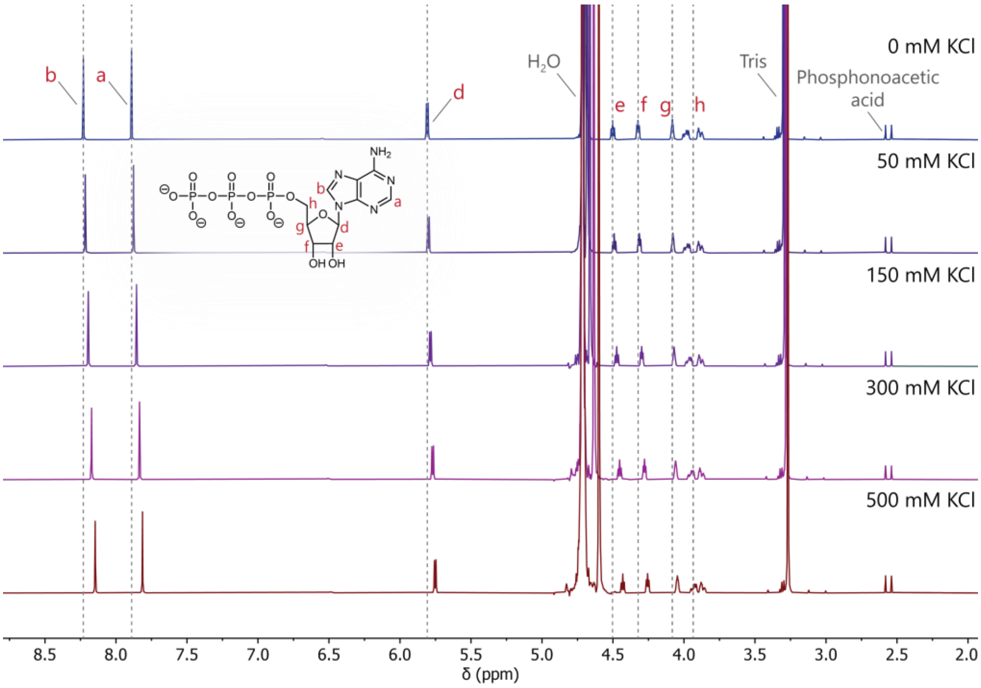

**Supplementary Figure 8:** [1]H-NMR-spectra for binding of KCl to the ribose and nucleobase protons on 25 mM ATP in 50 mM Tris pH 8.5 in 9 : 1 $H_2O$ : $D_2O$, with 20 mM phosphonoacetic acid in 9 : 1 $H_2O$ : $D_2O$ as internal standard in the inner tube of an NMR coaxial tube, measured at 5˚C. Full NMR spectra and spectra for other salts can be found on the Radboud Data Repository (Section 5). Two peaks are observed for $H_2O$, as the water in the inner coaxial tube has a different environment than the water in the outer coaxial tube. In the analysis for Figure 1g, the $K_{D,app}$'s of the *a* and *b* protons were averaged for binding to the nucleobase, and the $K_{D,app}$'s of the *d*, *e* and *f* protons were averaged for binding to the sugar.

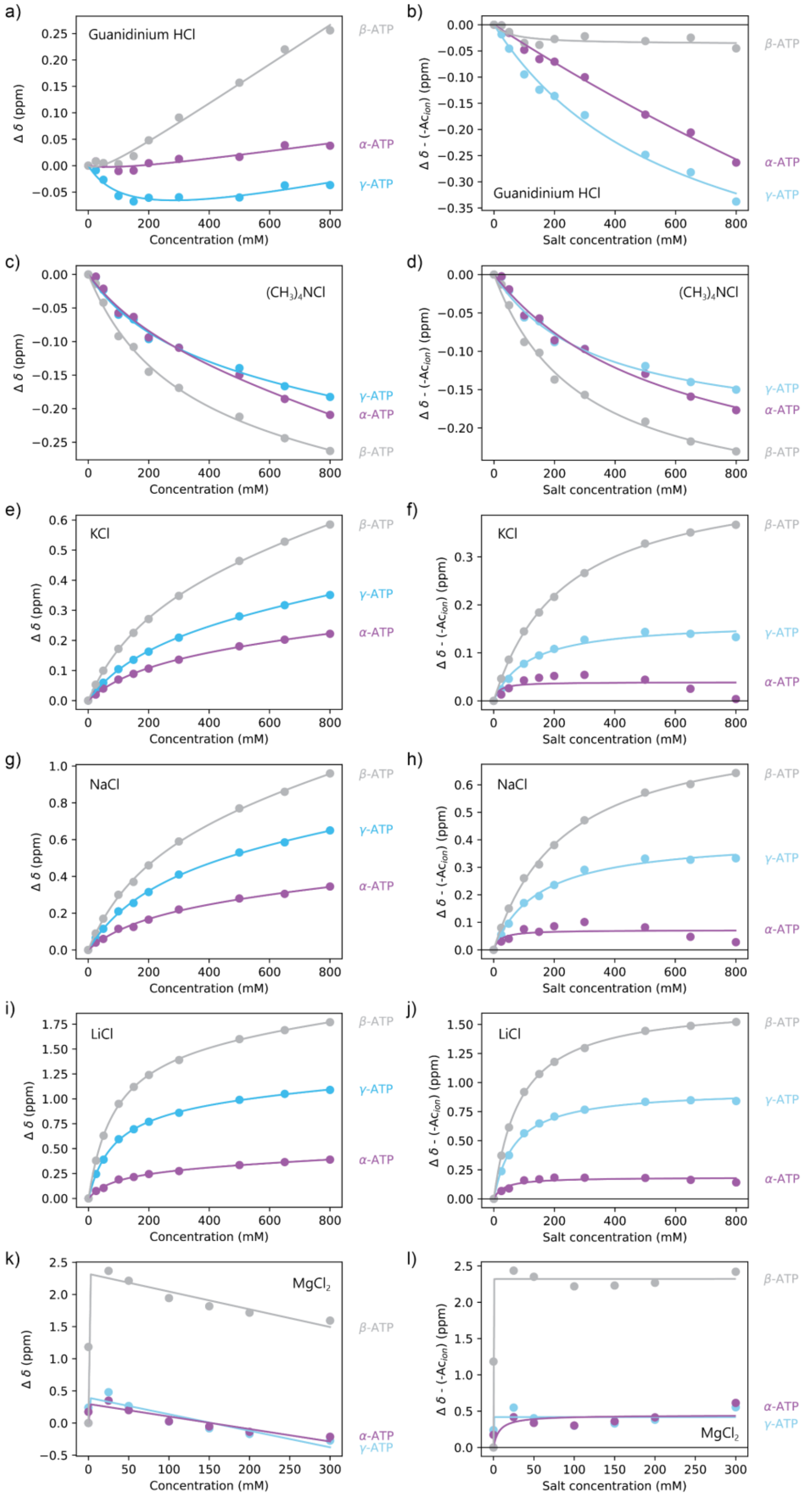

**Supplementary Figure 9: a,c,e,g,i,k)** Fitted binding curves for binding of chloride salts of different cations to the phosphates on 25 mM ATP at 5˚C. **b,d,f,h,j,l)** Fitted binding curves for the non-linear part $\Delta\delta - (-Ac_{\text{ion}}) = \frac{\Delta\delta_{\max} c_{\text{ion}}}{K_{\text{D,app}} + c_{\text{ion}}}$ for binding of chloride salts of different cations to the phosphates on 25 mM ATP at 5˚C. The difference in sign of Δδ-(-$Ac_{\text{ion}}$) between different ions likely indicates a different mechanism of binding.

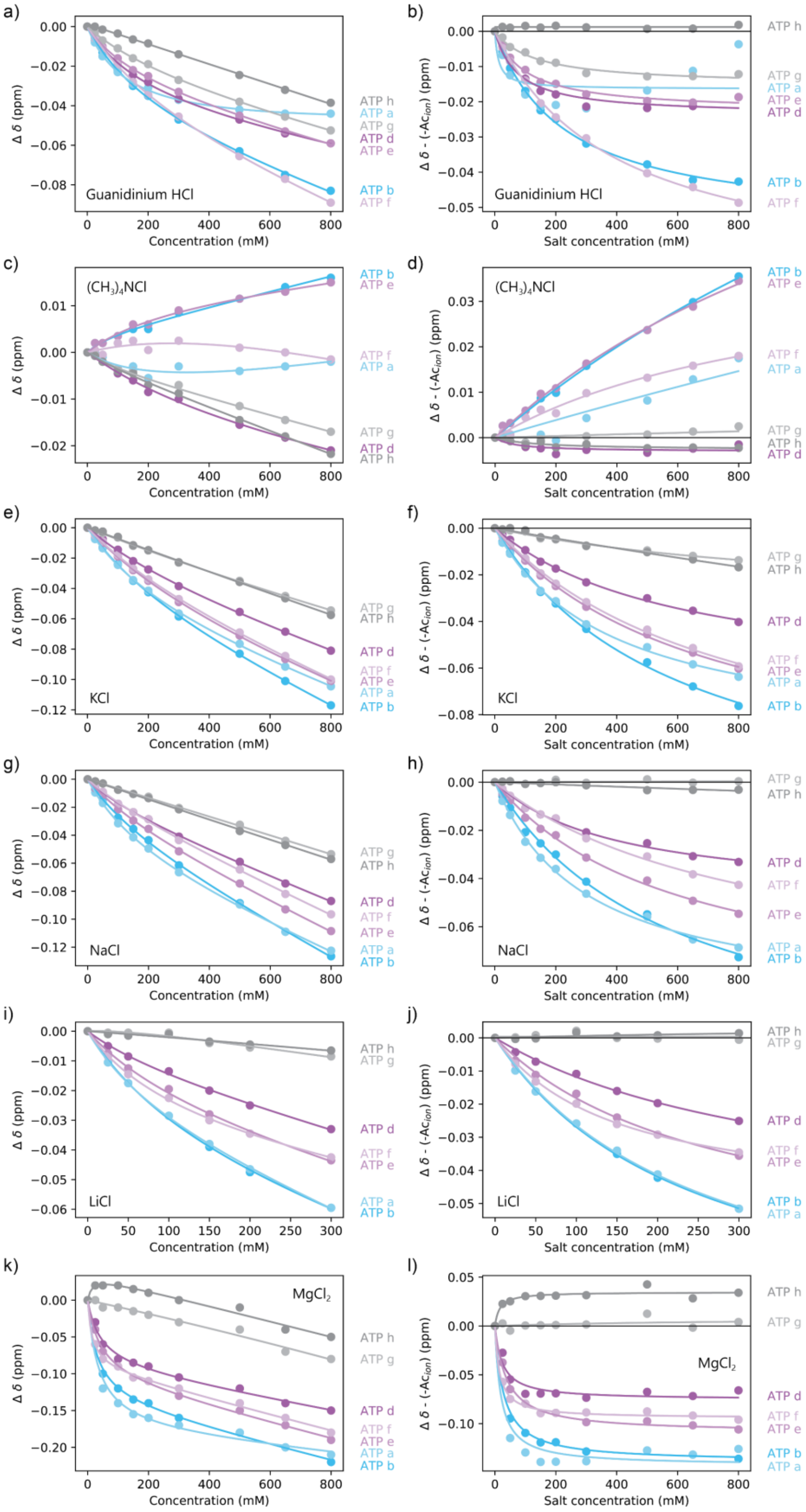

**Supplementary Figure 10: a,c,e,g,i,k)** Fitted binding curves for binding of chloride salts of different cations to the ribose (purple, grey) and nucleobase (blue) on 25 mM ATP at 5˚C. Grey binding curves were not taken into account for the average $K_{D,app}$ of the ribose. **b,d,f,h,j,l)** Fitted binding curves for the non-linear part of the binding curve. The difference in sign of Δδ-(-$Ac_{ion}$) between different ions likely indicates a different mechanism of binding.

##### 3.3. Supplementary data 'Ion binding is sequence specific and compacts protamine'

###### 3.3.1. Supplementary data MD-simulations

###### 3.3.1.1. Simulations with unbiased configuration

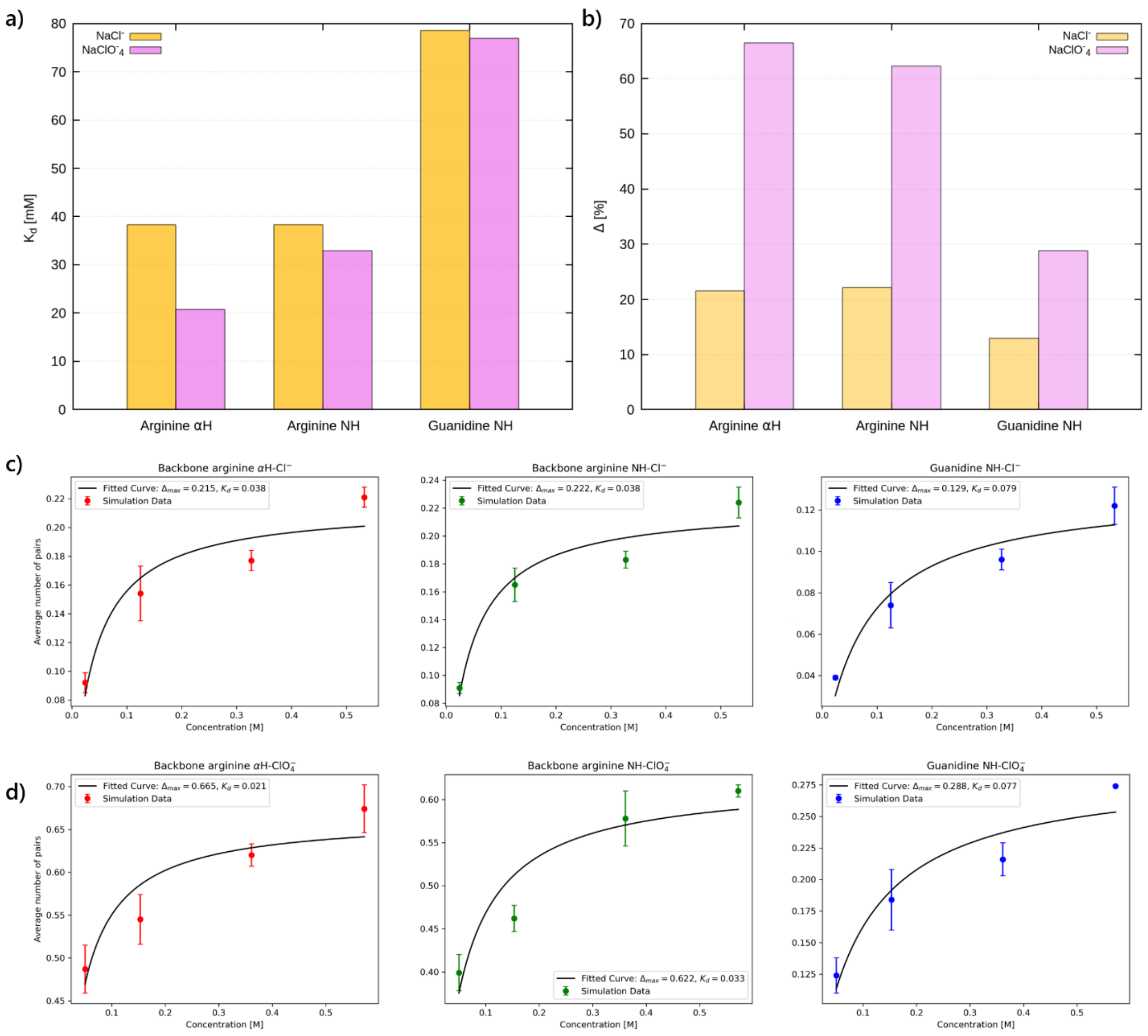

**Supplementary Figure 11:** Determination of dissociation constants from ion-binding MD simulations. **a)** $K_D$ of the protamine hydrogen-ion interactions fitted via the isothermal binding curve from the MD simulations at different concentrations. The obtained $K_D$'s show stronger binding to the backbone than to the arginine guanidinium, in accordance with the NMR binding data. **b)** The second parameter of the fit, Δ, which quantifies the asymptotic occupancy level, i.e. the saturation of binding sites, shows that $ClO_4^-$ ions bind more strongly to the backbone and guanidine compared to the $Cl^-$. **c, d)** Plots of the fitted binding curves and simulation data for **c)** $Cl^-$ and **d)** $ClO_4^-$. The binding does not saturate at high salt concentrations. Due to the fluctuation between a predominant open form and less prevalent closed state (Supporting Figure 10), not all binding sites are simultaneously available.

#### 3.3.1.2. Simulations with fixed configuration

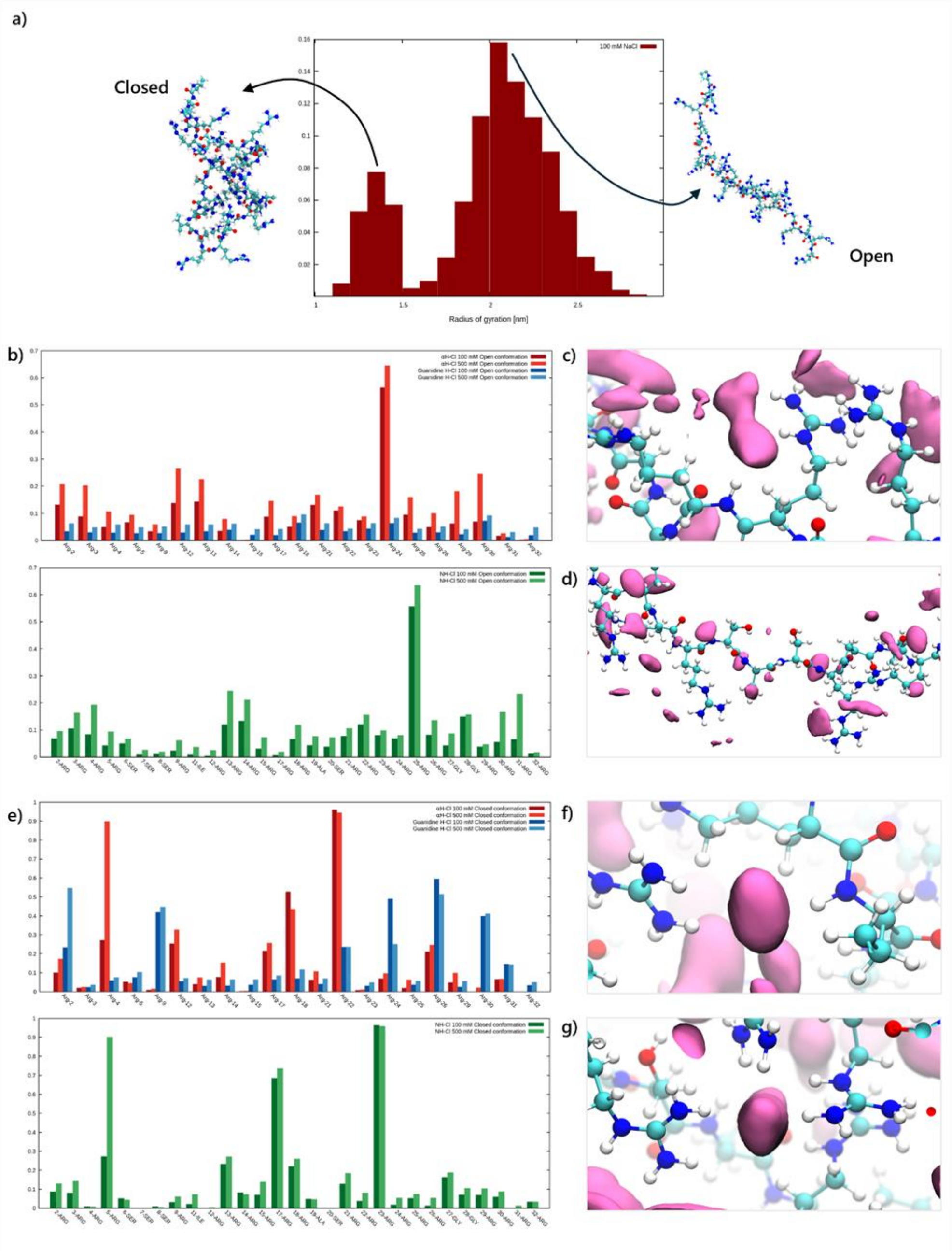

**Supplementary Figure 12:** MD simulations for protamine in a fixed open or closed configuration. **a)** Example distribution of the radius of gyration of protamine, showing a distinct open and closed population. **b)** Per-residue contact analysis for the open configuration: arginine α-protons and guanidine NH (top) and backbone NH (bottom). Contacts are significantly reduced compared to the unbiased simulations for all backbone residues except the α-proton and NH pair of residues 24 and 25. **c)** Snapshot of the open configuration showing the region with the highest ion density, corresponding to arginines 24 and 25, showing that the guanidine of residue 25 is oriented in such a way that the ions interacting with them are also interacting with the backbone hydrogens of residue 24, creating a binding pocket. **d)** Snapshot of the open configuration, showing the Ser-rich region of protamine. The serines are depleted of ions compared to the surrounding arginine-rich regions. **e)** Per-residue contact analysis for the closed configuration: arginine α-protons and guanidine NH (top) and backbone NH (bottom). Both the backbone and guanidines show heterogeneous ion affinities. **f)** Snapshot of the closed configuration showing a region of high ion density at the guanidine of residue 9 and the backbone hydrogen pair formed by the α-proton of residue 22 and the NH group of residue 23. **g)** Snapshot of the closed configuration showing a region of high ion density at the guanidines of residues 24, 26 and 30.

3.3.2. Supplementary data SAXS measurements

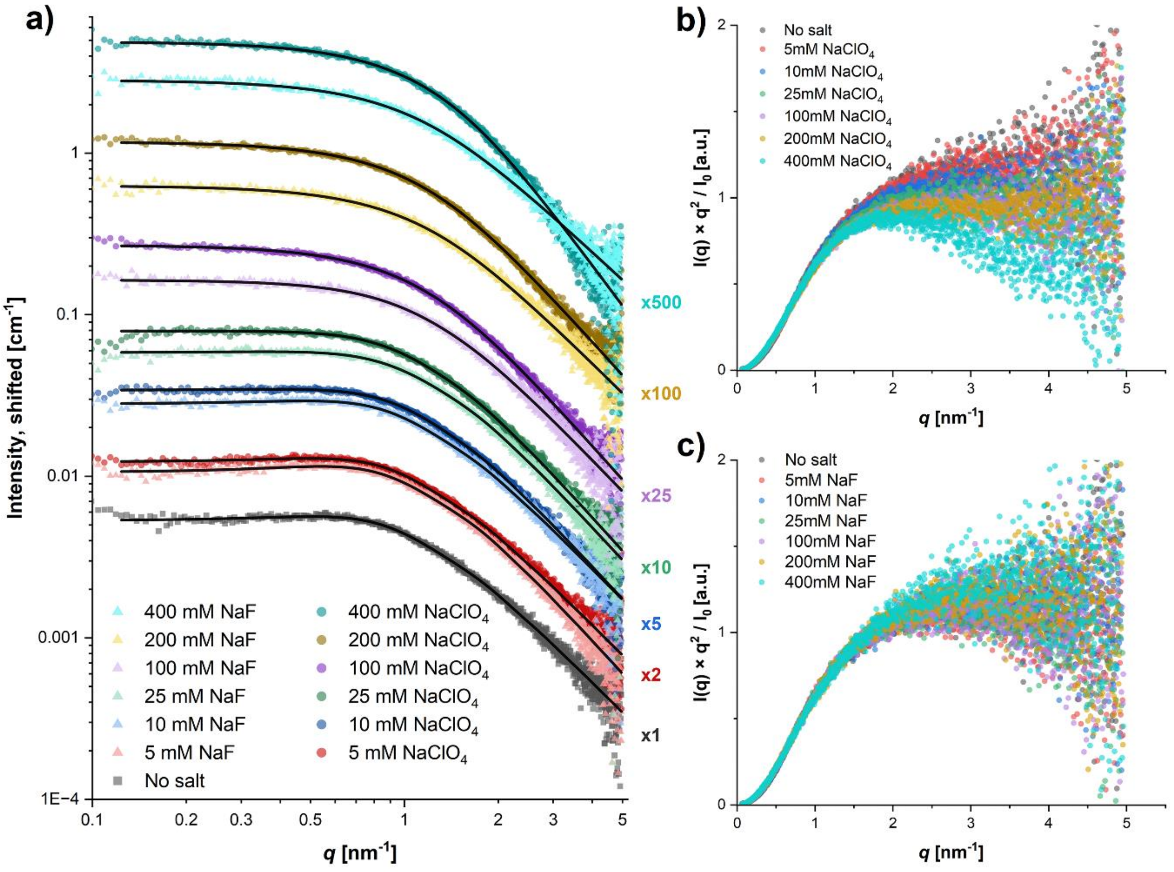

**Supplementary Figure 13:** Obtained SAXS curves for 1 mM protamine with different concentrations of $NaClO_4$ or NaF. **a)** SAXS curves fitted with generalized gaussian coil form factor, including a hard sphere structure factor to account for repulsion between peptides. Curves are displayed shifted by the factors shown on the right to avoid overlap. **b)** Kratky plot for different concentrations of $NaClO_4$ (Figure 2i in main text), showing a clear compaction of the protamine for higher $NaClO_4$ concentrations. **c)** Kratky plot for different concentrations of NaF, showing that the addition of NaF does not have a pronounced effect on the protamine conformation.

**Supplementary Table 3:** Parameters obtained from fitted SAXS curves of 1 mM (4 mg/mL) protamine with different concentrations of $NaClO_4$.

| **[$NaClO_4$] (mM)** | **Radius of gyration $R_g$ (nm)** | **Flory excluded volume parameter ν** | **Forward scattering $I_0$ (cm$^{-1}$)** | **Effective hard sphere radius $R_{HS}$ (nm)** | **Effective volume fraction $\phi_{HS}$** |
|---|---|---|---|---|---|
| 0 | 1.275 ± 0.002 | 0.507 ± 0.010 | 0.0070 ± 0.0013 | 3.30 ± 0.01 | 0.033 ± 0.019 |
| 5 | 1.275 ± 0.002 | 0.504 ± 0.008 | 0.0080 ± 0.0012 | 3.24 ± 0.01 | 0.033 ± 0.016 |
| 10 | 1.263 ± 0.002 | 0.468 ± 0.008 | 0.0084 ± 0.0012 | 3.17 ± 0.01 | 0.025 ± 0.016 |
| 25 | 1.283 ± 0.002 | 0.456 ± 0.007 | 0.0092 ± 0.0011 | 3.35 ± 0.01 | 0.018 ± 0.018 |
| 100 | 1.319 ± 0.002 | 0.454 ± 0.007 | 0.0109 ± 0.0014 | 3.08 ± 0.19 | 0.001 ± 0.061 |
| 200 | 1.323 ± 0.002 | 0.459 ± 0.008 | 0.01179 ± 0.00209 | 2.52 ± 9.02 | 0.001 ± 0.095 |
| 400 | 1.272 ± 0.002 | 0.379 ± 0.011 | 0.00978 ± 0.00207 | - | 0.000 |

##### 3.4. Supplementary data 'Ion partitioning into condensates follows LMWA'

###### 3.4.1. NMR calibration curves

###### 3.4.1.1. NMR Calibration curves $^{7}Li^{+}$

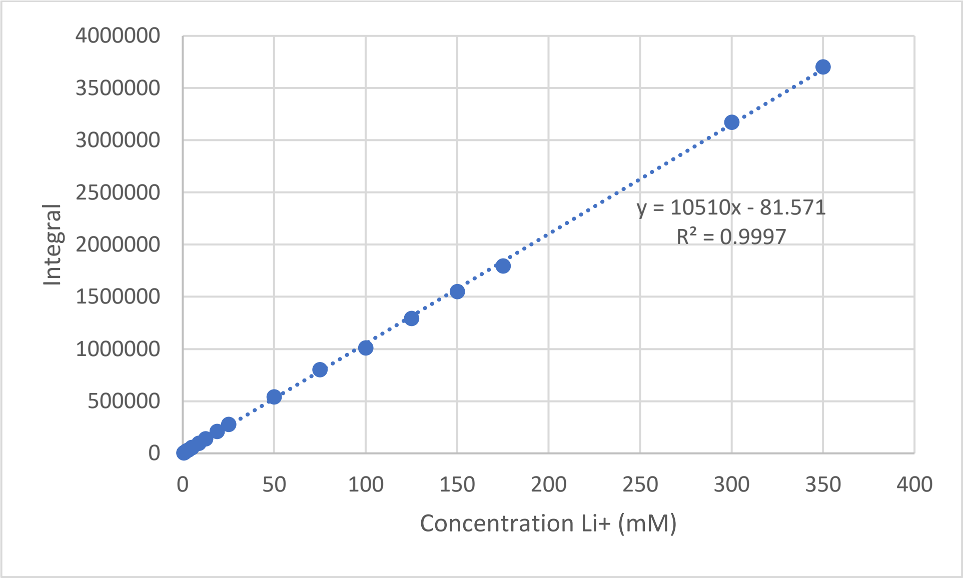

**Supplementary Figure 14:** $^{7}$Li-NMR calibration curve for 0.5 – 350 mM lithium chloride, used for determining the dilute phase $Li^{+}$ concentration. P1 = 12.15 µs, d1 = 200 s, rg = 1030, ns = 8. See Supplementary Information Section 1.3 for full description of settings.

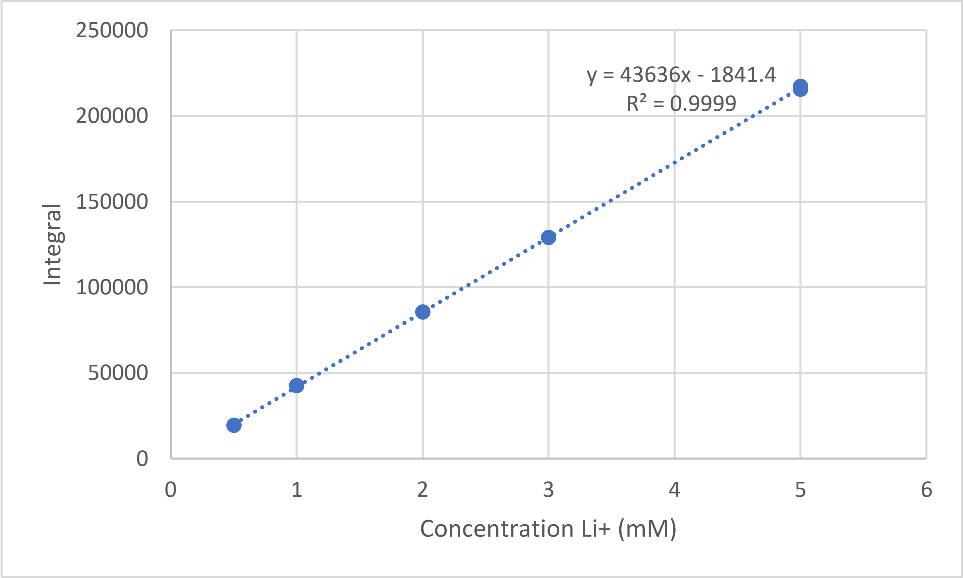

**Supplementary Figure 15:** $^{7}$Li-NMR calibration curve for 0.5 – 5 mM lithium chloride, used for determining the condensate phase $Li^{+}$ concentration. P1 = 12.15 µs, d1 = 200 s, rg = 2050, ns = 16. See Supplementary Information Section 1.3 for full description of settings.

3.4.1.2. NMR Calibration curves $^{23}Na^{+}$

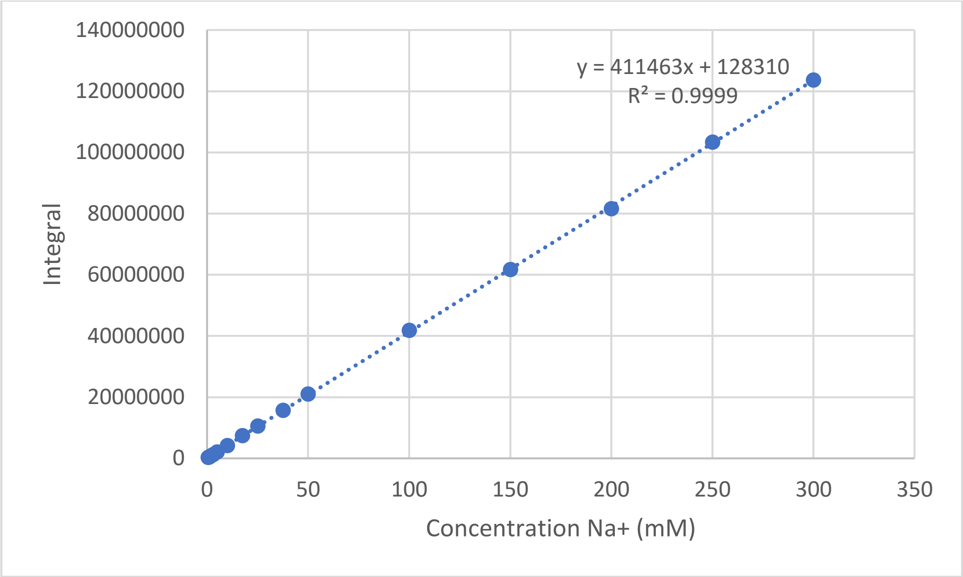

**Supplementary Figure 16:** $^{23}$Na-NMR calibration curve for 0.5 – 300 mM sodium chloride, used for determining the dilute phase $Na^{+}$ concentration. P1 = 20.5 µs, d1 = 2 s, rg = 1820, ns = 1024. See Supplementary Information Section 1.3 for full description of settings.

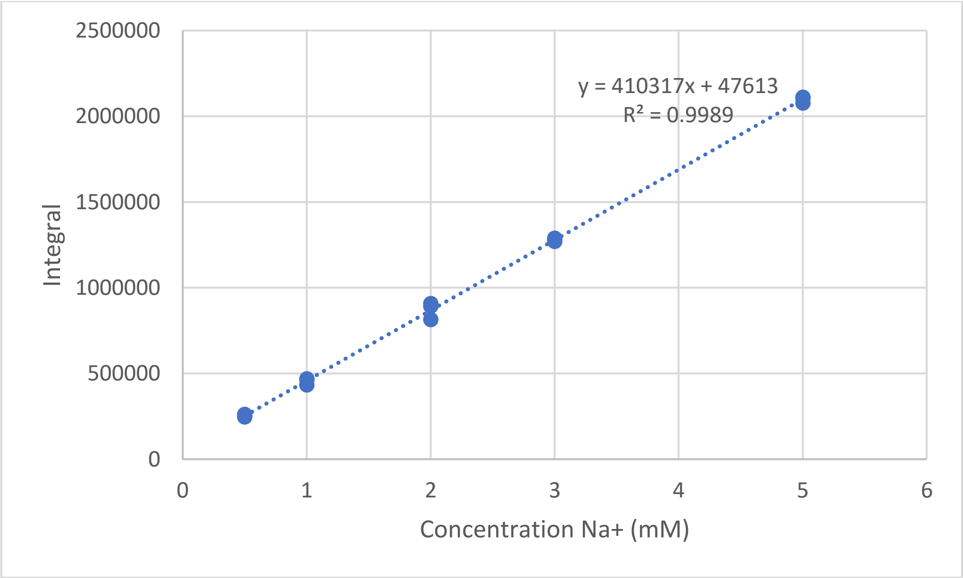

**Supplementary Figure 17:** $^{23}$Na-NMR calibration curve for 0.5 – 5 mM sodium chloride, used for determining the condensate phase $Na^{+}$ concentration. P1 = 20.5 µs, d1 = 2 s, rg = 1820, ns = 1024. See Supplementary Information Section 1.3 for full description of settings.

3.4.1.3. NMR Calibration curves $^{25}Mg^{2+}$

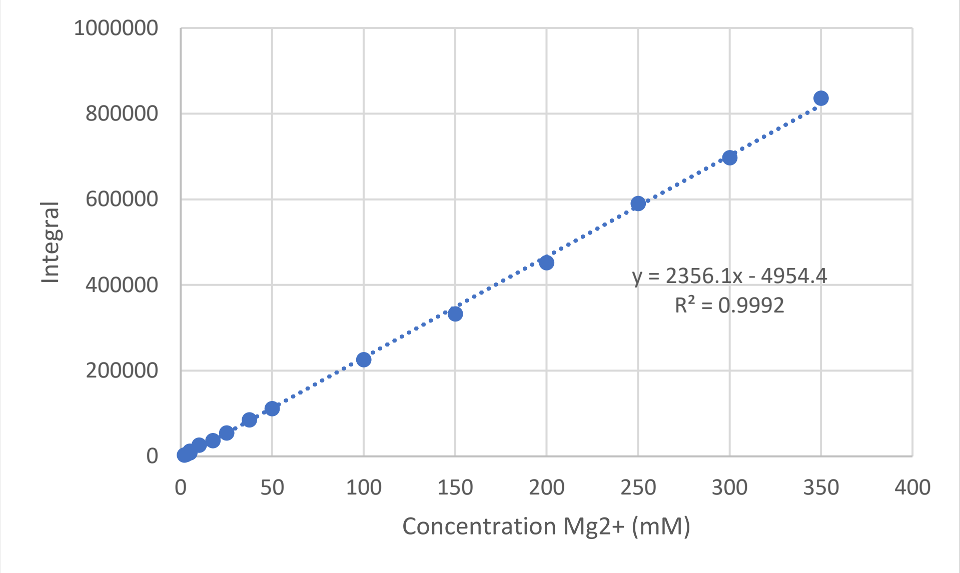

**Supplementary Figure 18:** $^{25}$Mg-NMR calibration curve for 0.5 – 350 mM magnesium chloride (molecule-based), used for determining the dilute phase $Mg^{2+}$ concentration. P1 = 20.0 µs, d1 = 0.1 s, rg = 203, ns = 4096. See Supplementary Information Section 1.3 for full description of settings.

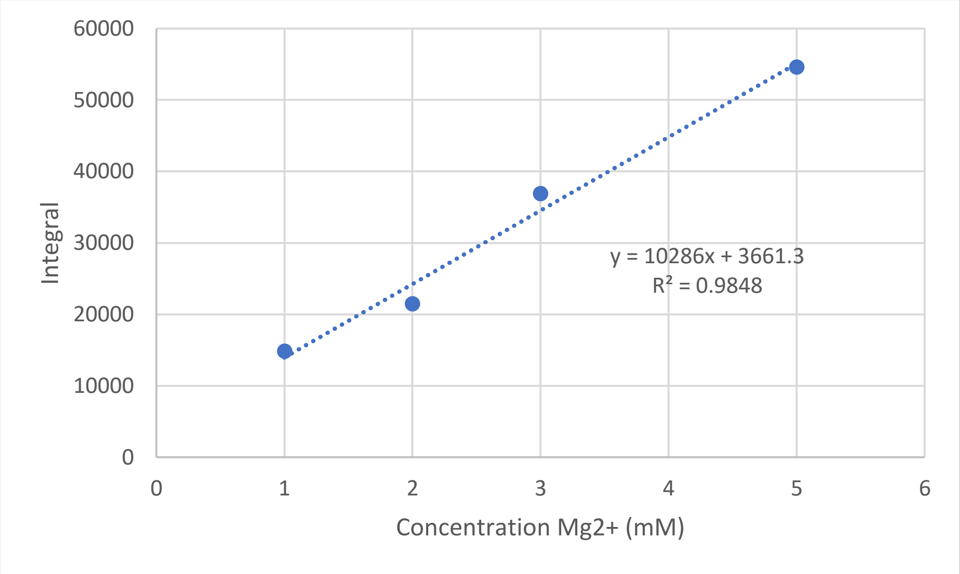

**Supplementary Figure 19:** $^{25}$Mg-NMR calibration curve for 1 – 5 mM magnesium chloride (molecule-based), used for determining the condensate phase $Mg^{2+}$ concentration. P1 = 20.0 µs, d1 = 0.1 s, rg = 203, ns = 20480. See Supplementary Information Section 1.3 for full description of settings.

3.4.1.4. NMR Calibration curves $^{35}Cl^-$

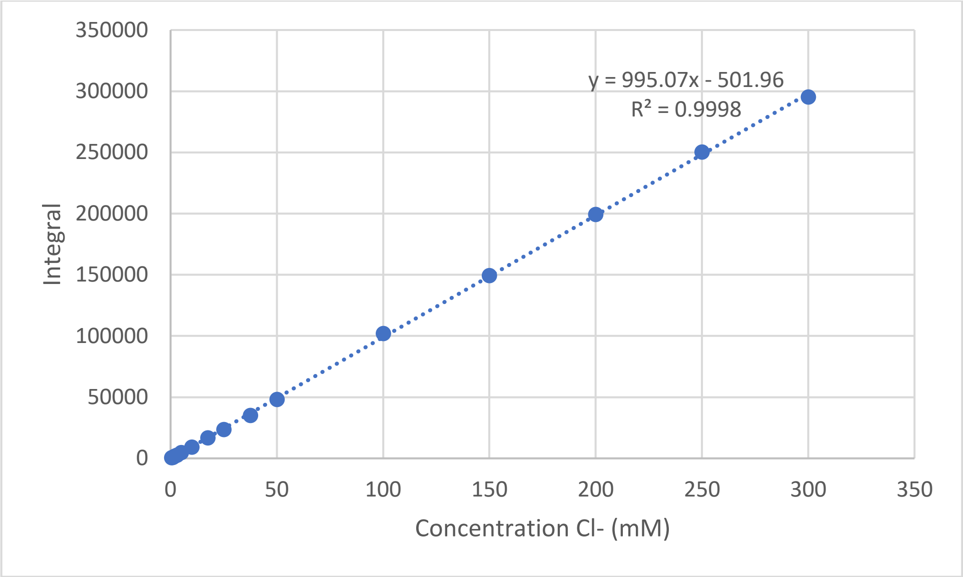

**Supplementary Figure 20:** $^{35}Cl$-NMR calibration curve for 0.5 – 300 mM sodium chloride, used for determining the dilute phase $Cl^-$ concentration. P1 = 15.5 µs, d1 = 2 s, rg = 50.8, ns = 512. See Supplementary Information Section 1.3 for full description of settings.

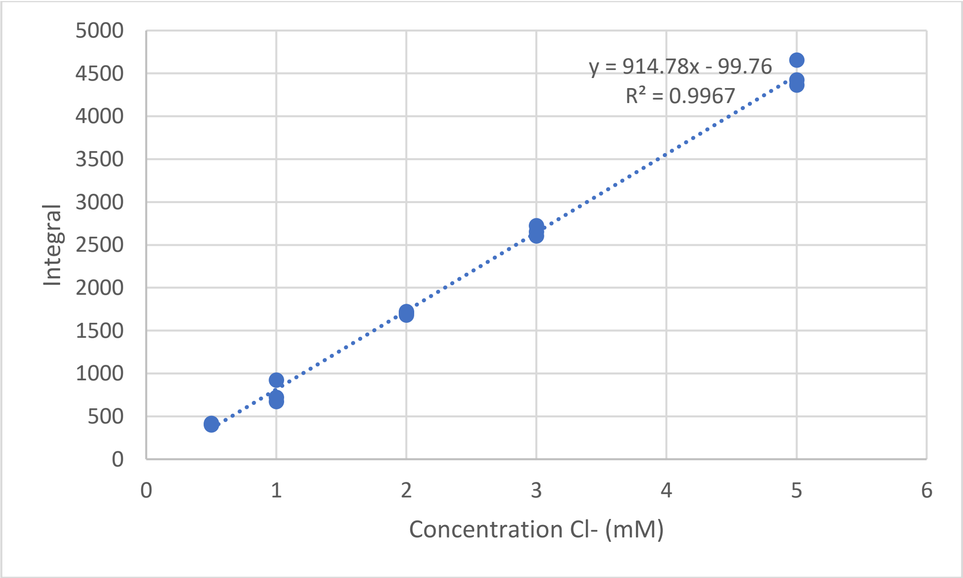

**Supplementary Figure 21:** $^{35}Cl$-NMR calibration curve for 0.5 – 5 mM sodium chloride, used for determining the condensate phase $Cl^-$ concentration. P1 = 15.5 µs, d1 = 2 s, rg = 50.8, ns = 512. See Supplementary Information Section 1.3 for full description of settings.

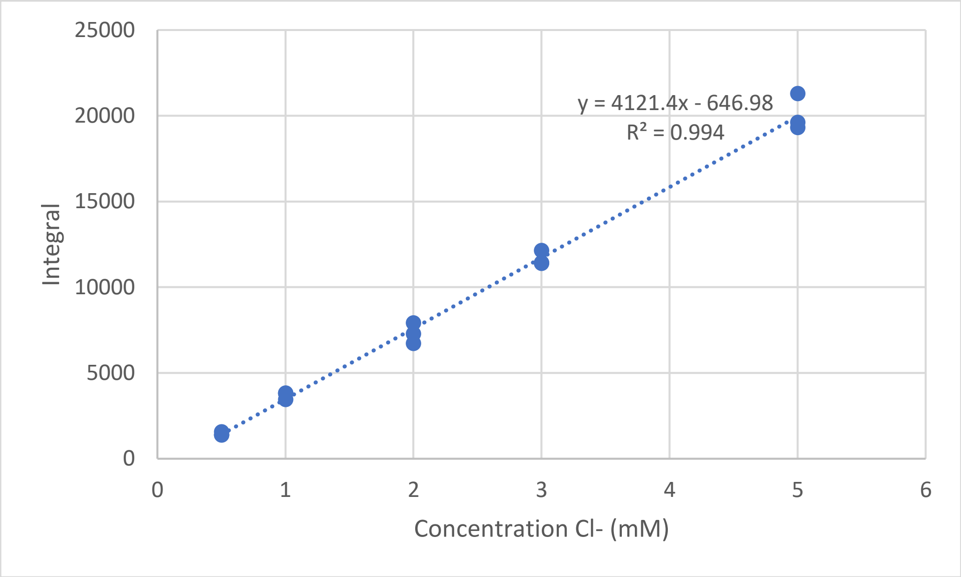

**Supplementary Figure 22:** $^{35}$Cl-NMR calibration curve for 0.5 – 5 mM sodium chloride, used for determining the condensate phase $Cl^-$ concentration. P1 = 15.5 µs, d1 = 2 s, rg = 64, ns = 2048. See Supplementary Information Section 1.3 for full description of settings.

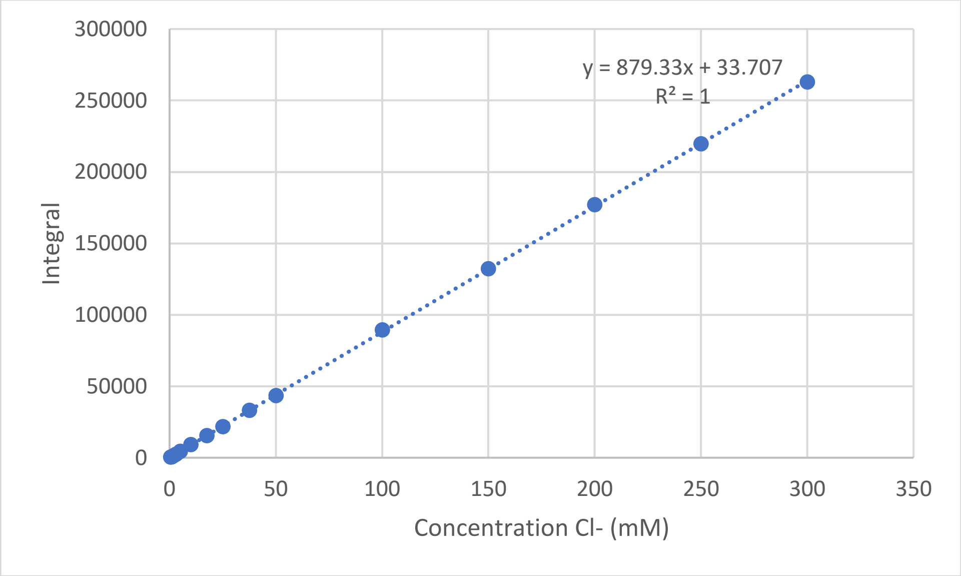

**Supplementary Figure 23:** $^{35}$Cl-NMR calibration curve for 0.5 – 300 mM sodium chloride, used for determining the dilute phase $Cl^-$ concentration in samples with $ClO_4^-$. P1 = 15.5 µs, d1 = 2 s, sw = 1195 ppm, O1P = 500 ppm, rg = 50.8, ns = 1024. See Supplementary Information Section 1.3 for full description of settings.

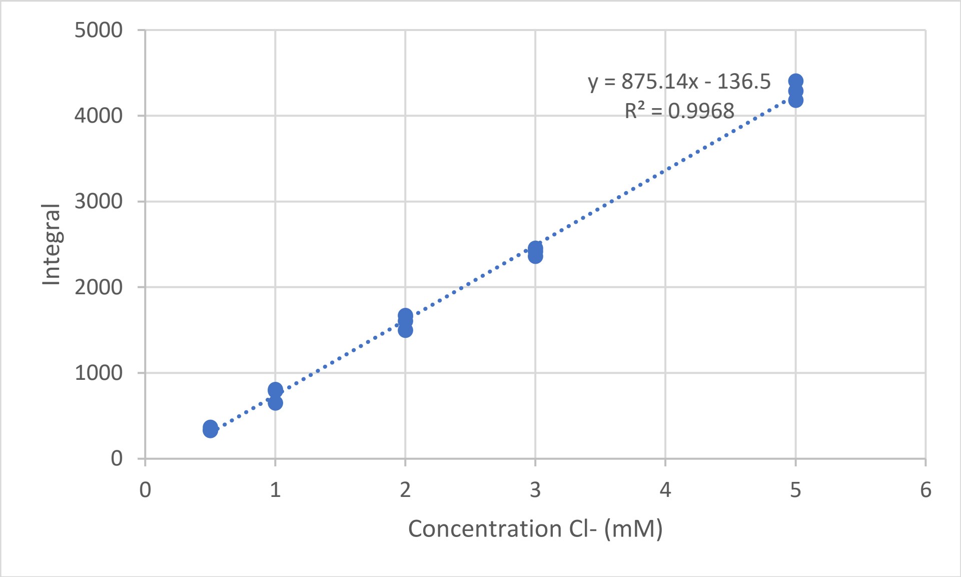

**Supplementary Figure 24:** $^{35}$Cl-NMR calibration curve for 0.5 – 5 mM sodium chloride, used for determining the condensate phase $Cl^-$ concentration in samples with $ClO_4^-$. P1 = 15.5 µs, d1 = 2 s, sw = 1195 ppm, O1P = 500 ppm, rg = 50.8, ns = 1024. See Supplementary Information Section 1.3 for full description of settings.

3.4.1.5. NMR Calibration curves $^{35}ClO_4^-$

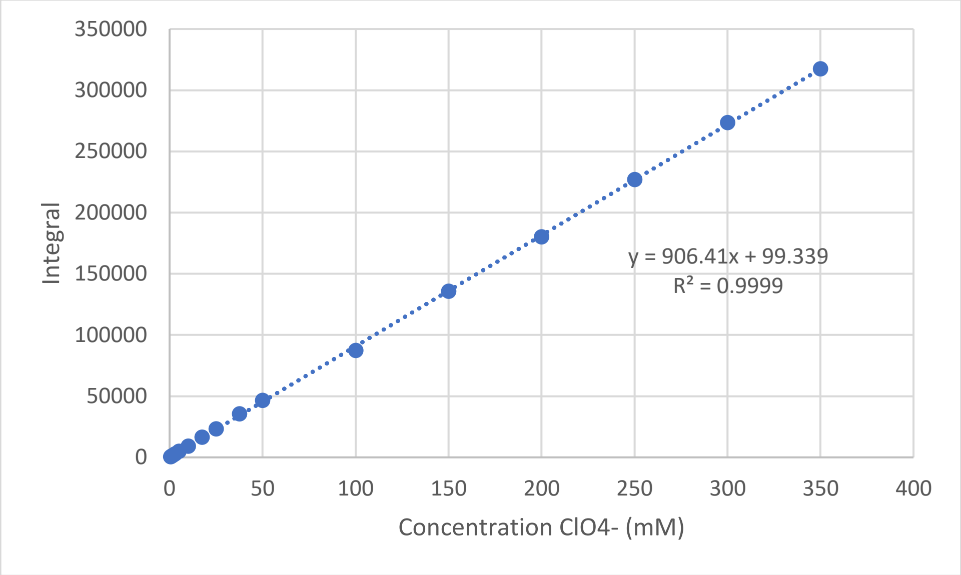

**Supplementary Figure 25:** $^{35}$Cl-NMR calibration curve for 0.5 – 350 mM sodium perchlorate, used for determining the dilute phase $ClO_4^-$ concentration. P1 = 15.5 µs, d1 = 2 s, sw = 1195 ppm, O1P = 500 ppm, rg = 50.8, ns = 1024. See Supplementary Information Section 1.3 for full description of settings.

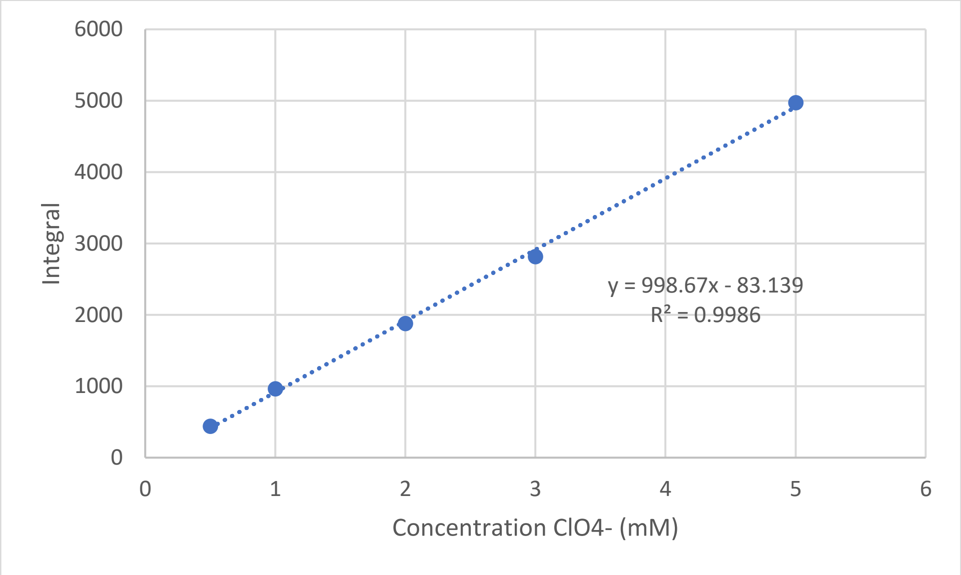

**Supplementary Figure 26:** $^{35}$Cl-NMR calibration curve for 0.5 – 5 mM sodium perchlorate, used for determining the condensate phase $ClO_4^-$ concentration. P1 = 15.5 µs, d1 = 2 s, sw = 1195 ppm, O1P = 500 ppm, rg = 50.8, ns = 1024. See Supplementary Information Section 1.3 for full description of settings.

3.4.1.6. NMR Calibration curves $^{39}K^{+}$

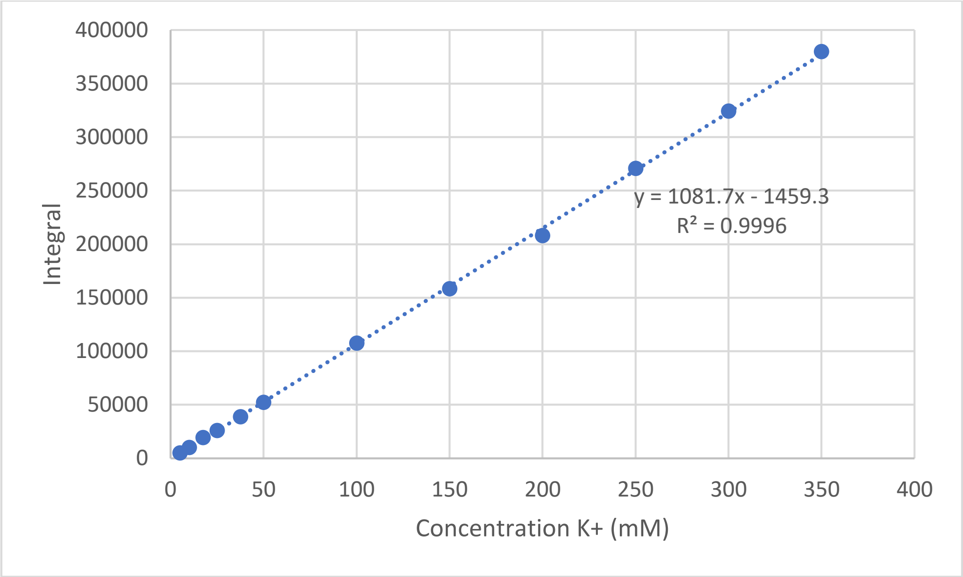

**Supplementary Figure 27:** $^{39}$K-NMR calibration curve for 0.5 – 350 mM potassium chloride, used for determining the dilute phase $K^{+}$ concentration. P1 = 25.0 µs, d1 = 0.1 s, rg = 101, ns = 4096. See Supplementary Information Section 1.3 for full description of settings.

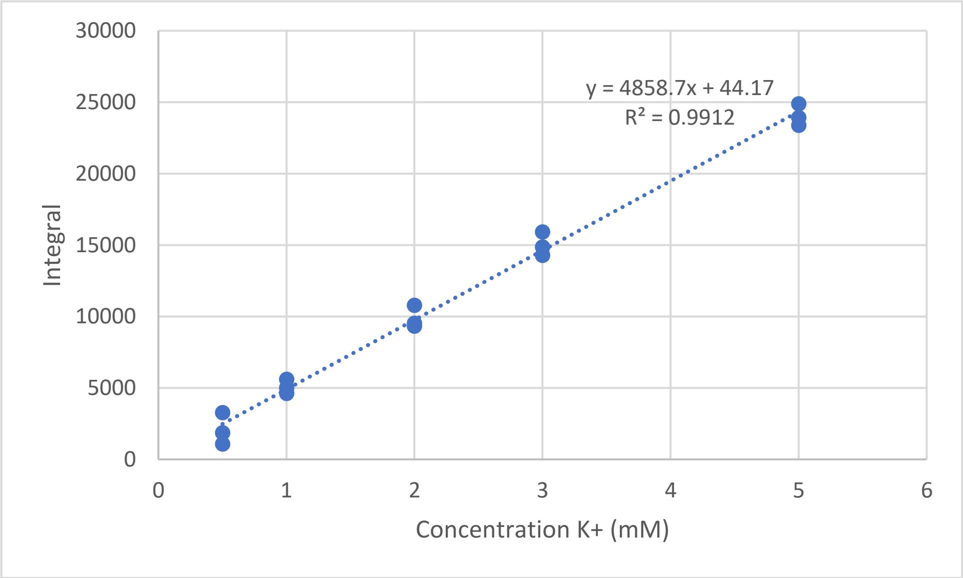

**Supplementary Figure 28:** $^{39}$K-NMR calibration curve for 0.5 – 5 mM potassium chloride, used for determining the condensate phase $K^{+}$ concentration. P1 = 25.0 µs, d1 = 0.1 s, rg = 203, ns = 20480. See Supplementary Information Section 1.3 for full description of settings.

3.4.1.7. NMR Calibration curves $^{81}Br^-$

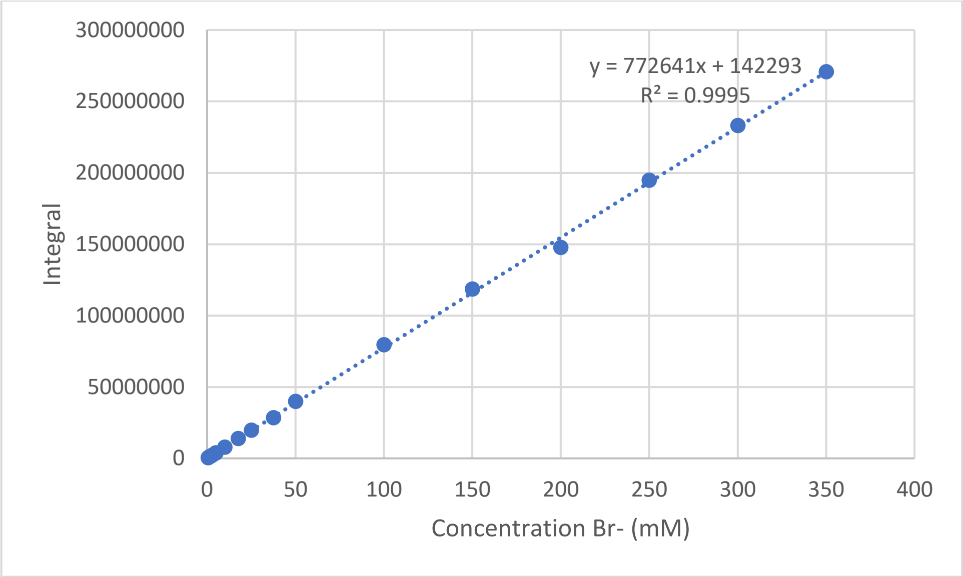

**Supplementary Figure 29:** $^{81}$Br-NMR calibration curve for 0.5 – 350 mM sodium bromide, used for determining the dilute phase $Br^-$ concentration. P1 = 12.0 µs, d1 = 1 s, rg = 2050, ns = 2048. See Supplementary Information Section 1.3 for full description of settings.

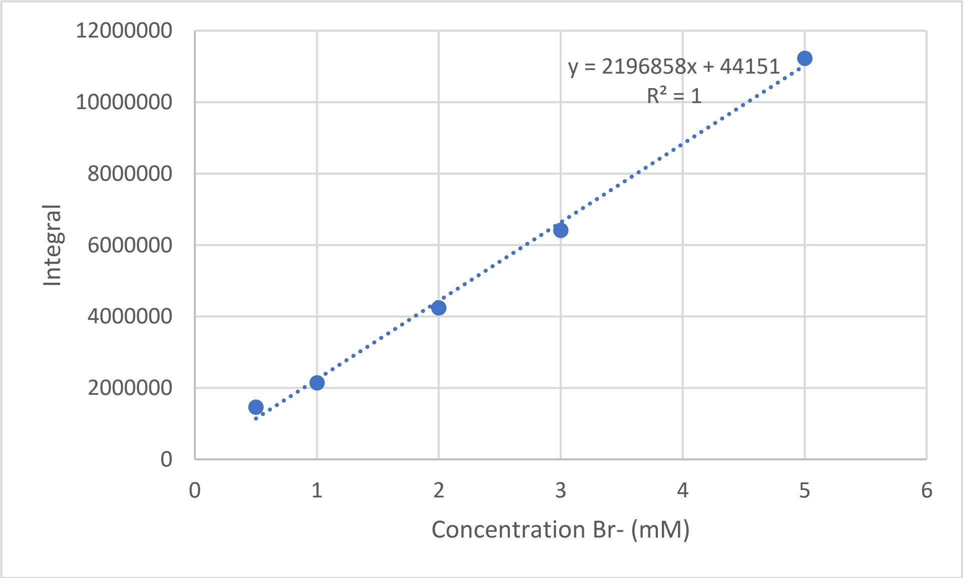

**Supplementary Figure 30:** $^{81}$Br-NMR calibration curve for 0.5 – 5 mM sodium bromide, used for determining the condensate phase $Br^-$ concentration. P1 = 12.0 µs, d1 = 1 s, rg = 2050, ns = 6144. See Supplementary Information Section 1.3 for full description of settings.

3.4.1.8. NMR Calibration curves $^{133}Cs^{+}$

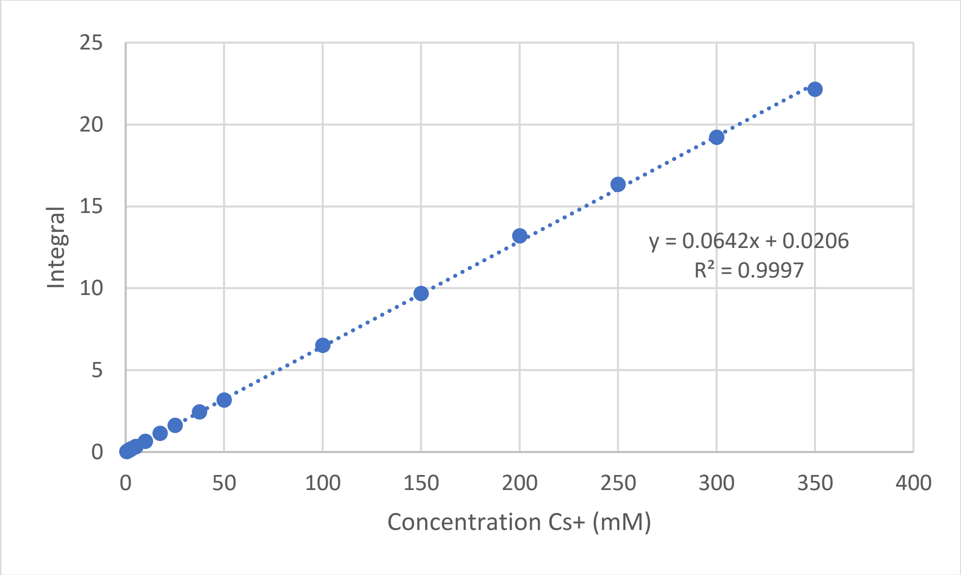

**Supplementary Figure 31:** $^{133}$Cs-NMR calibration curve for 0.5 – 350 mM cesium chloride, used for determining the dilute phase $Cs^{+}$ concentration. P1 = 7.5 µs, d1 = 23 s, rg = 50, ns = 32. See Supplementary Information Section 1.3 for full description of settings.

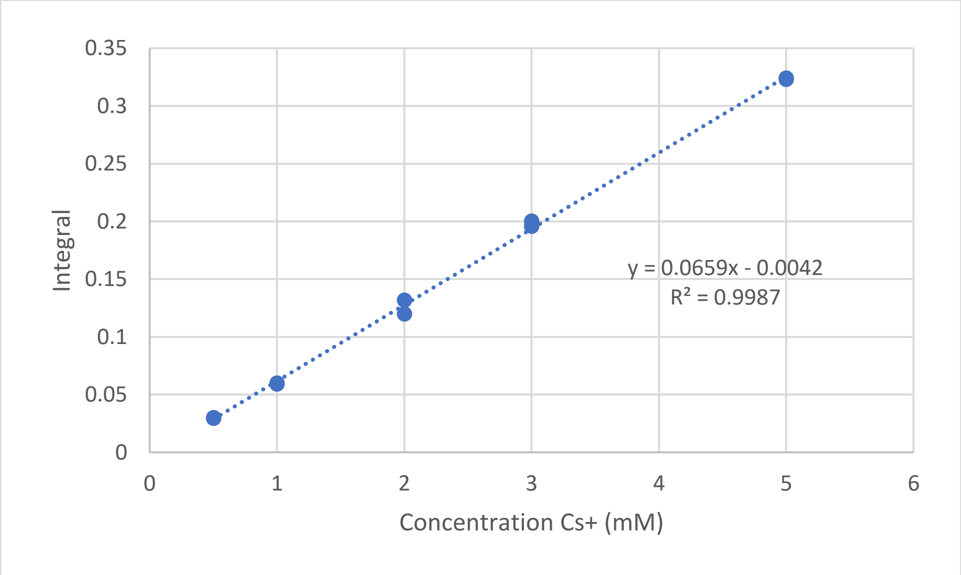

**Supplementary Figure 32:** $^{133}$Cs-NMR calibration curve for 0.5 – 5 mM cesium chloride, used for determining the condensate phase $Cs^{+}$ concentration. P1 = 7.5 µs, d1 = 23 s, rg = 50, ns = 32. See Supplementary Information Section 1.3 for full description of settings.

##### 3.4.2. NMR spectra and measured concentrations ion partitioning

In this section we show an example of the $^{1}H$, $^{7}Li$, $^{13}C$, $^{19}F$, $^{23}Na$, $^{31}P$, $^{35}Cl$, $^{39}K$, $^{81}Br$ and $^{133}Cs$-NMR measurements we performed to determine the concentration protamine, ATP, Tris and different anions and cations in the condensate and dilute phase. The full set of raw NMR spectra can be found on the Radboud Data repository (Section 5).

###### 3.4.2.1. No salt

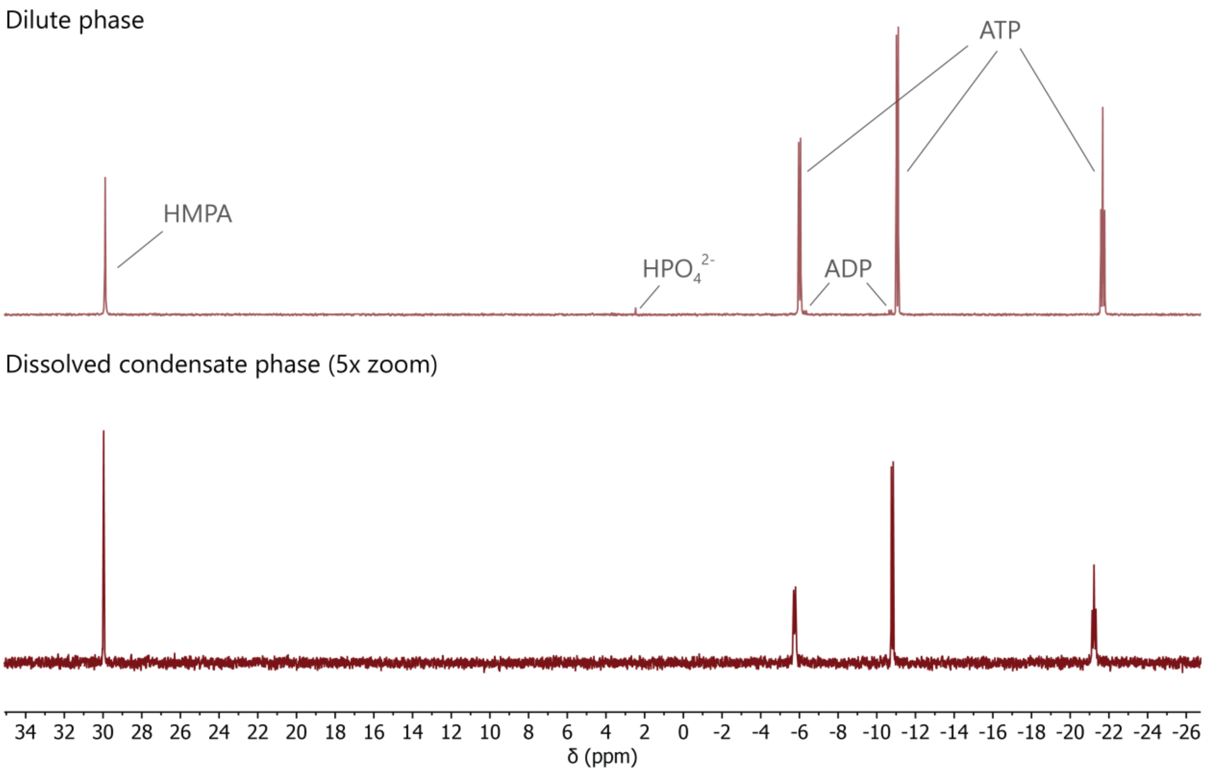

**Supplementary Figure 33:** $^{31}P$-NMR-spectra of the dilute phase and dissolved condensate phase for condensates of 1 mM protamine chloride / 25 mM ATP without added salt in 50 mM Tris pH 8.5, used to determine the concentration of ATP. HMPA is the hexamethylphosphoramide internal standard. Only very minor peaks for ADP and phosphate are observed, showing that no significant hydrolysis of ATP takes place during the sample preparation.

**Supplementary Table 4:** Measured charge-based concentrations in the condensate phase and dilute phase and $K_P$ for condensates without added salt.

| Compound | $c_{condensate}$ (mM) | $c_{dilute}$ (mM) | $K_P$ |
|---|---|---|---|
| Protamine | 2191.3 ± 36.3 | 6.65 ± 0.05 | 329.5 ± 5.9 |
| ATP | 2340.1 ± 57.8 | 82.5 ± 0.9 | 28.4 ± 0.8 |
| Tris | 65.1 ± 0.7 | 52.5 ± 0.4 | 1.24 ± 0.02 |
| $Na^+$ | 183.3 ± 5.8 | 94.2 ± 0.6 | 1.95 ± 0.06 |
| $Cl^-$ | 161.0 ± 15.1 | 41.7 ± 0.8 | 3.86 ± 0.37 |

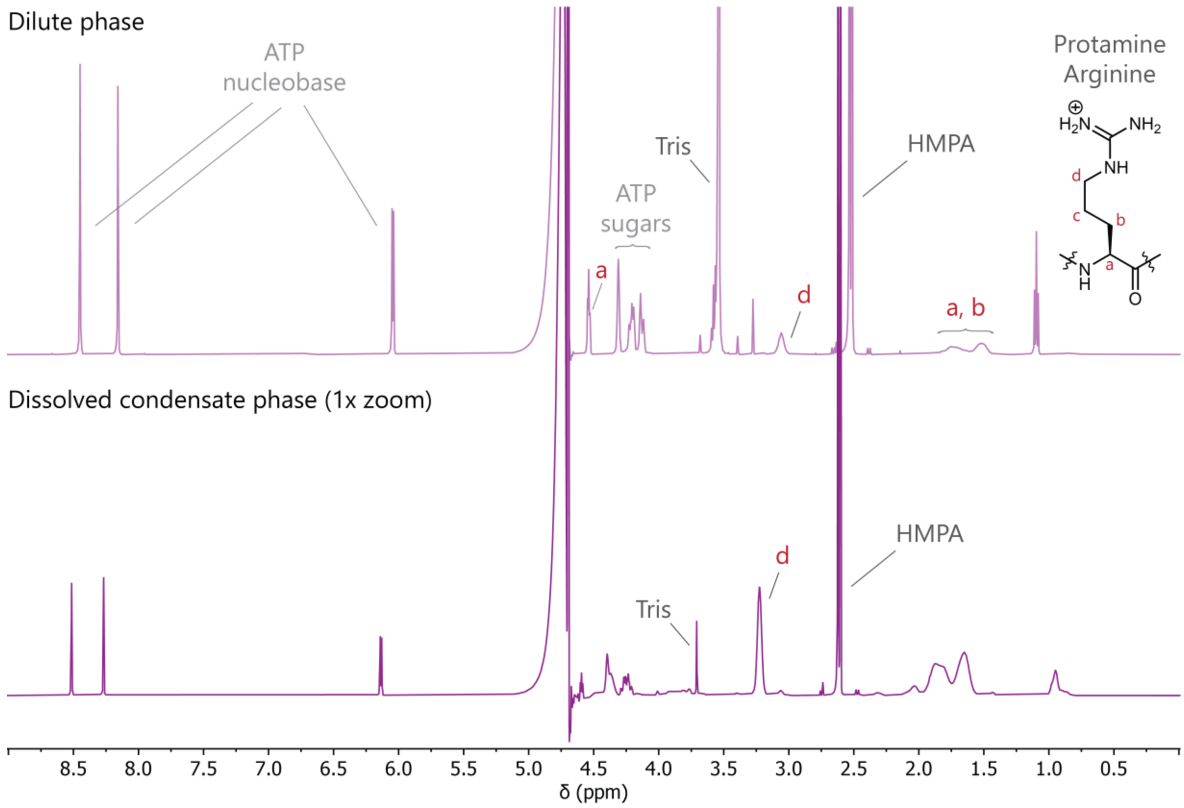

**Supplementary Figure 34:** $^{1}$H-NMR-spectra of the dilute phase and dissolved condensate phase for condensates of 1 mM protamine chloride / 25 mM ATP without added salt in 50 mM Tris pH 8.5, used to determine the concentration of protamine and Tris. For the quantification of protamine, the peak of the *d* protons was used.

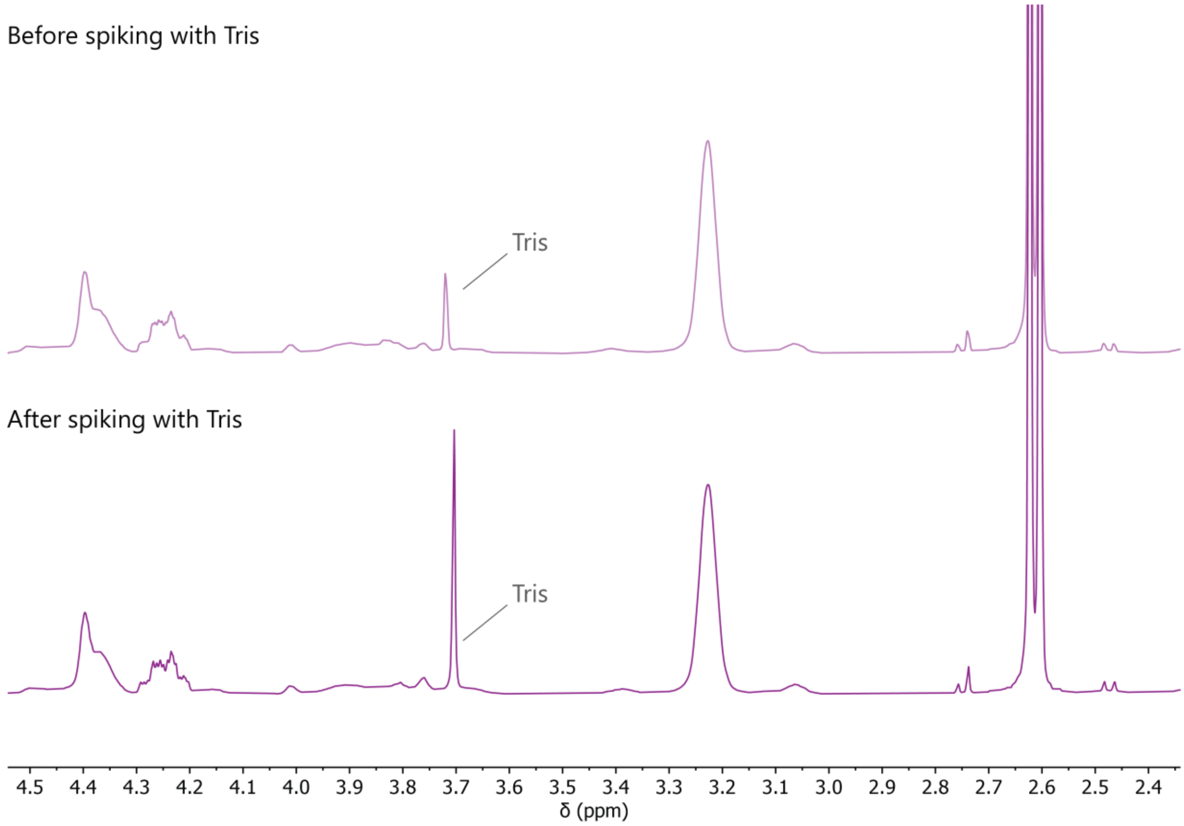

**Supplementary Figure 35:** Spiking with Tris to confirm the chemical shift of its peak in the condensate phase. $^{1}$H-NMR-spectra of the dissolved condensate phase of 1 mM protamine chloride / 25 mM ATP condensates prepared with 50 mM LiCl in 50 mM Tris pH 8.5, before and after spiking with an additional 0.5 mM Tris pH 8.5.

3.4.2.2. $Na_2HPO_4$

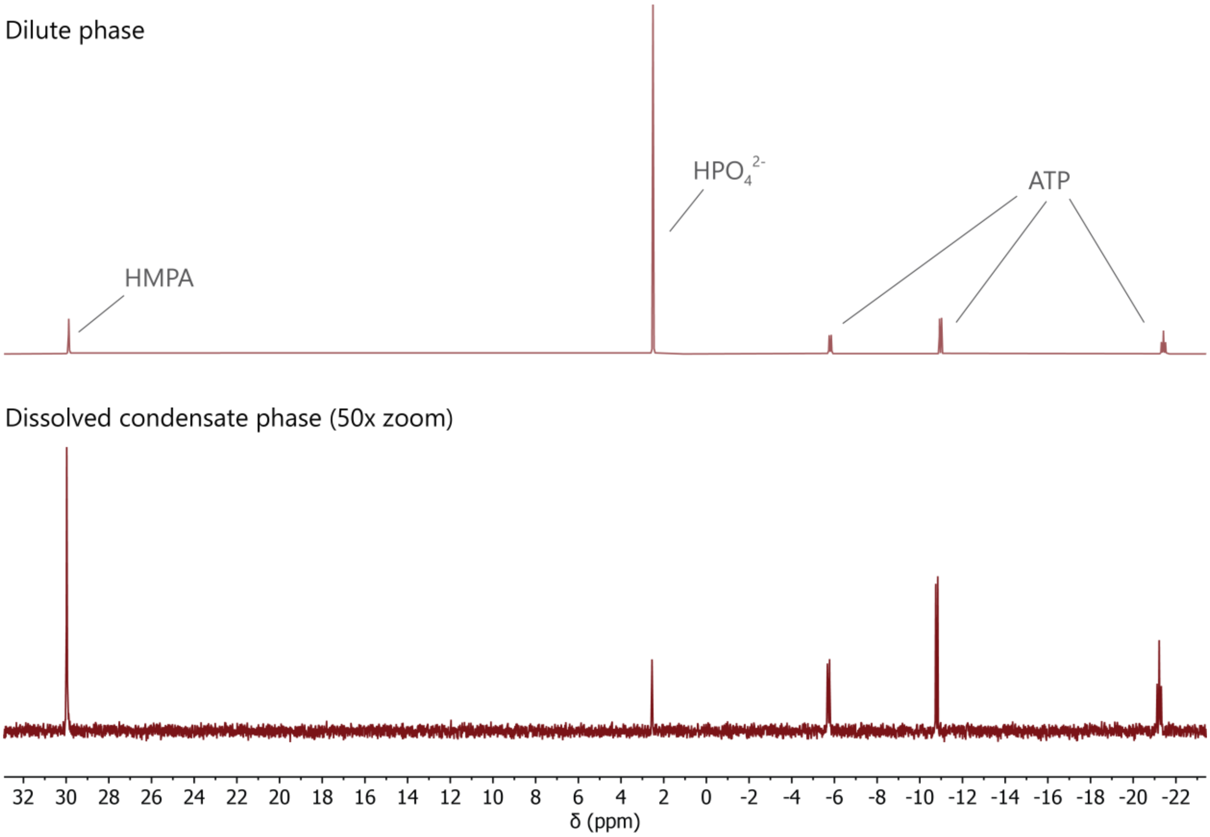

**Supplementary Figure 36:** $^{31}$P-NMR-spectra of the dilute phase and dissolved condensate phase for partitioning of 200 mM $Na_2HPO_4$ (charge-based) in condensates of 1 mM protamine chloride / 25 mM ATP in 50 mM Tris pH 8.5, used to determine the concentration of $HPO_4^{2-}$ and ATP.

**Supplementary Table 5:** Measured charge-based concentrations in the condensate phase and dilute phase and $K_P$ for $Na_2HPO_4$ partitioning.

| Compound | $c_{condensate}$ (mM) | $c_{dilute}$ (mM) | $K_P$ |
|---|---|---|---|
| Protamine | 2200.5 ± 38.6 | 10.8 ± 0.1 | 202.9 ± 3.7 |
| ATP | 2095.1 ± 113.1 | 88.1 ± 0.3 | 23.8 ± 1.3 |
| Tris | 59.1 ± 4.1 | 53.7 ± 0.2 | 1.10 ± 0.08 |
| $Na^+$ | 317.9 ± 21.6 | 279.6 ± 0.9 | 1.14 ± 0.08 |
| $Cl^-$ | 276.9 ± 12.6 | 40.7 ± 0.3 | 6.80 ± 0.31 |
| $HPO_4^{2-}$ | 380.1 ± 15.1 | 211.0 ± 0.6 | 1.80 ± 0.07 |

3.4.2.3. NaF

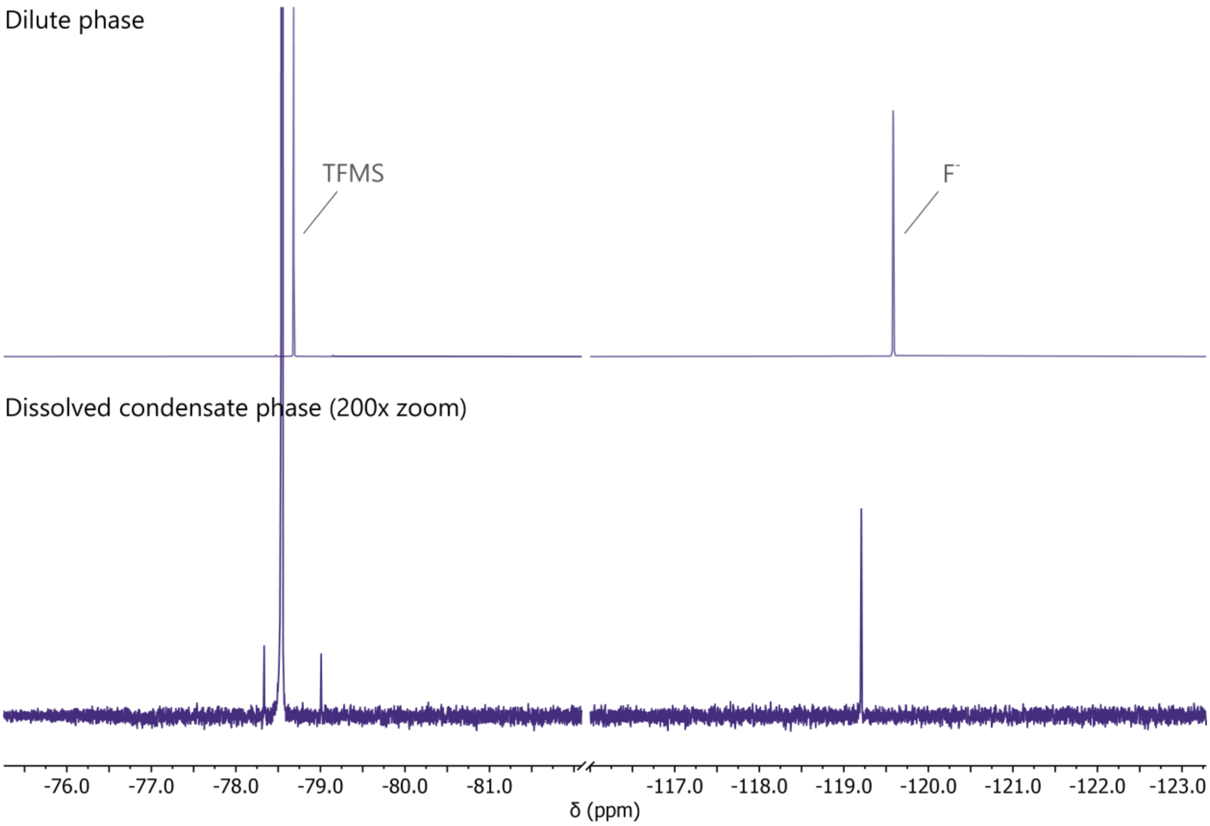

**Supplementary Figure 37:** [19]F-NMR-spectra of the dilute phase and dissolved condensate phase for partitioning of 100 mM NaF in condensates of 1 mM protamine chloride / 25 mM ATP in 50 mM Tris pH 8.5, used to determine the concentration of $F^-$ in reference to a sodium trifluoromethanesulfonate (TFMS) internal standard.

**Supplementary Table 6:** Measured charge-based concentrations in the condensate phase and dilute phase and $K_P$ for NaF partitioning. For the $Na^+$ concentration the added amount of $Na^+$ from the sodium trifluoromethanesulfonate internal standard was subtracted from the data.

| Compound | $c_{condensate}$ (mM) | $c_{dilute}$ (mM) | $K_P$ |
|---|---|---|---|
| Protamine | 1979.9 ± 79.1 | 7.41 ± 0.03 | 267.0 ± 10.7 |
| ATP | 2098.0 ± 112.2 | 82.9 ± 1.1 | 25.3 ± 1.4 |
| Tris | 54.9 ± 3.8 | 52.2 ± 0.3 | 1.05 ± 0.07 |
| $Na^+$ | 136.9 ± 33.2 | 184.9 ± 1.7 | 0.74 ± 0.18 |
| $Cl^-$ | 182.7 ± 27.2 | 41.8 ± 0.4 | 4.37 ± 0.65 |
| $F^-$ | 47.2 ± 1.9 | 104.9 ± 0.2 | 0.45 ± 0.02 |

3.4.2.4. NaBr

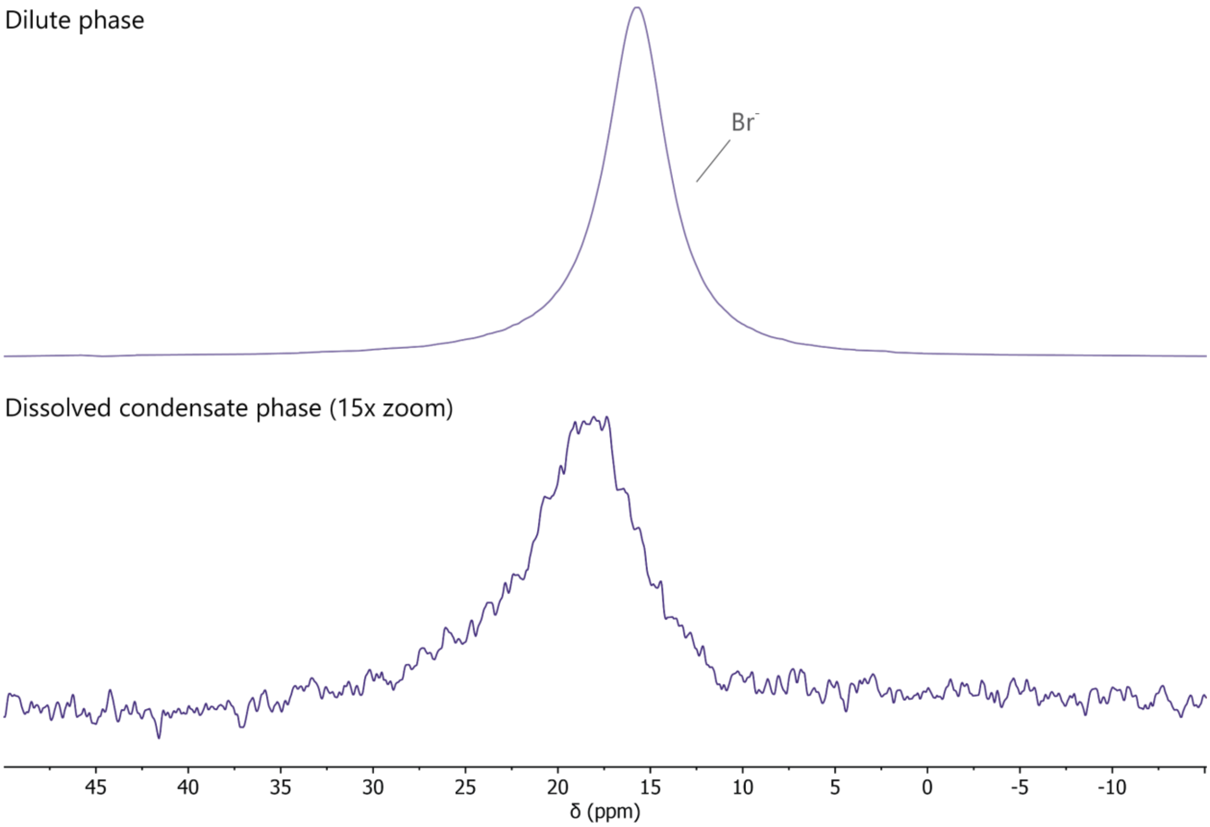

**Supplementary Figure 38:** $^{81}$Br-NMR-spectra of the dilute phase and dissolved condensate phase for partitioning of 100 mM NaBr in condensates of 1 mM protamine chloride / 25 mM ATP in 50 mM Tris pH 8.5, used to determine the concentration of $Br^-$.

**Supplementary Table 7:** Measured charge-based concentrations in the condensate phase and dilute phase and $K_P$ for NaF partitioning. The concentration of $Cl^-$ was not determined because the condensates were dissolved in 1 M KCl.

| Compound | $c_{condensate}$ (mM) | $c_{dilute}$ (mM) | $K_P$ |
|---|---|---|---|
| Protamine | 2135.5 ± 18.4 | 7.87 ± 0.1 | 271.4 ± 4.0 |
| ATP | 2325.0 ± 60.1 | 86.1 ± 1.3 | 27.0 ± 0.8 |
| Tris | 55.4 ± 0.07 | 53.3 ± 0.5 | 1.04 ± 0.01 |
| Na+ | 154.2 ± 3.0 | 186.7 ± 0.7 | 0.83 ± 0.02 |
| $Br^-$ | 125.2 ± 4.6 | 99.8 ± 0.8 | 1.26 ± 0.05 |

3.4.2.5. $NaClO_4$

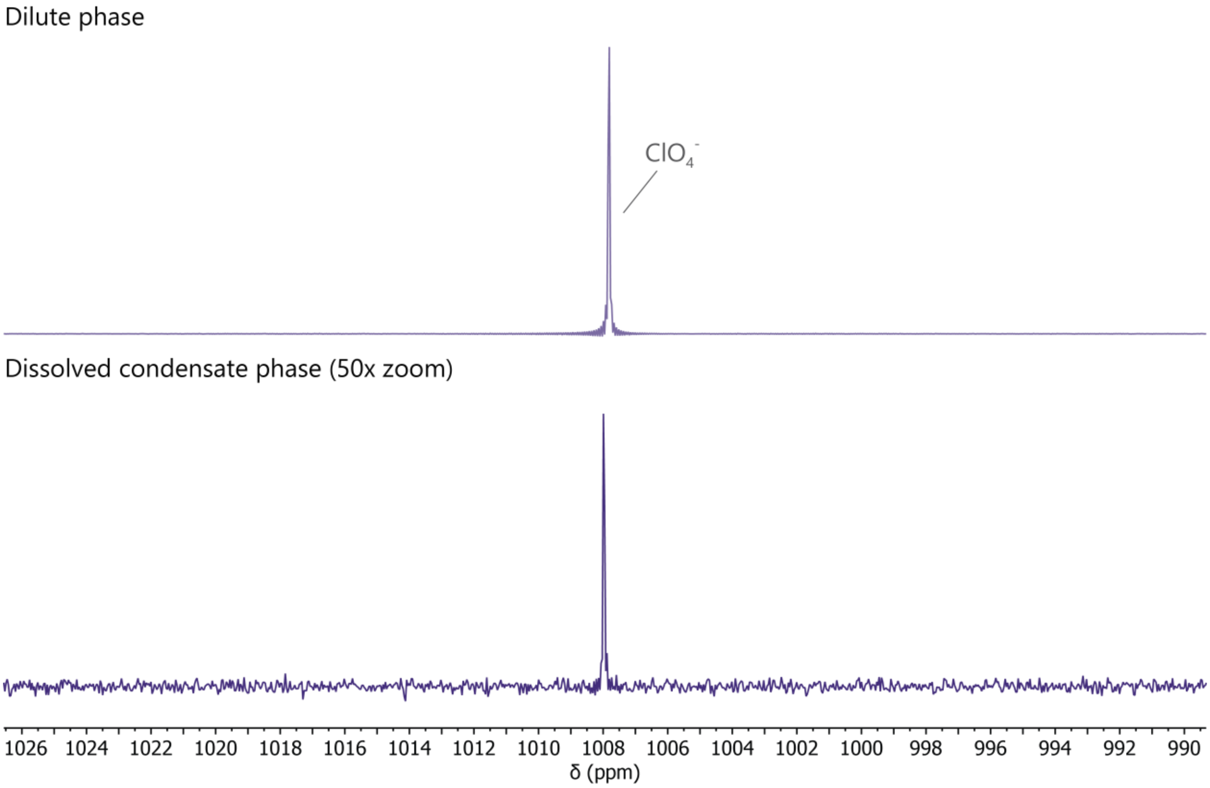

**Supplementary Figure 39:** $^{35}$Cl-NMR-spectra of the dilute phase and dissolved condensate phase for partitioning of 100 mM $NaClO_4$ in condensates of 1 mM protamine chloride / 25 mM ATP in 50 mM Tris pH 8.5, used to determine the concentration of $ClO_4^-$.

**Supplementary Table 8:** Measured charge-based concentrations in the condensate phase and dilute phase and $K_P$ for $NaClO_4$ partitioning.

| Compound | $c_{condensate}$ (mM) | $c_{dilute}$ (mM) | $K_P$ |
|---|---|---|---|
| Protamine | 2134.9 ± 11.2 | 7.63 ± 0.04 | 279.7 ± 2.0 |
| ATP | 2130.2 ± 18.8 | 85.5 ± 1.8 | 24.9 ± 0.6 |
| Tris | 53.3 ± 1.8 | 52.3 ± 0.2 | 1.02 ± 0.03 |
| $Na^+$ | 213.8 ± 1.4 | 184.5 ± 1.1 | 1.16 ± 0.01 |
| $Cl^-$ | 203.5 ± 4.9 | 40.6 ± 0.6 | 5.02 ± 0.14 |
| $ClO_4^-$ | 248.6 ± 7.1 | 96.9 ± 0.2 | 2.57 ± 0.07 |

3.4.2.6. KS$^{13}$CN

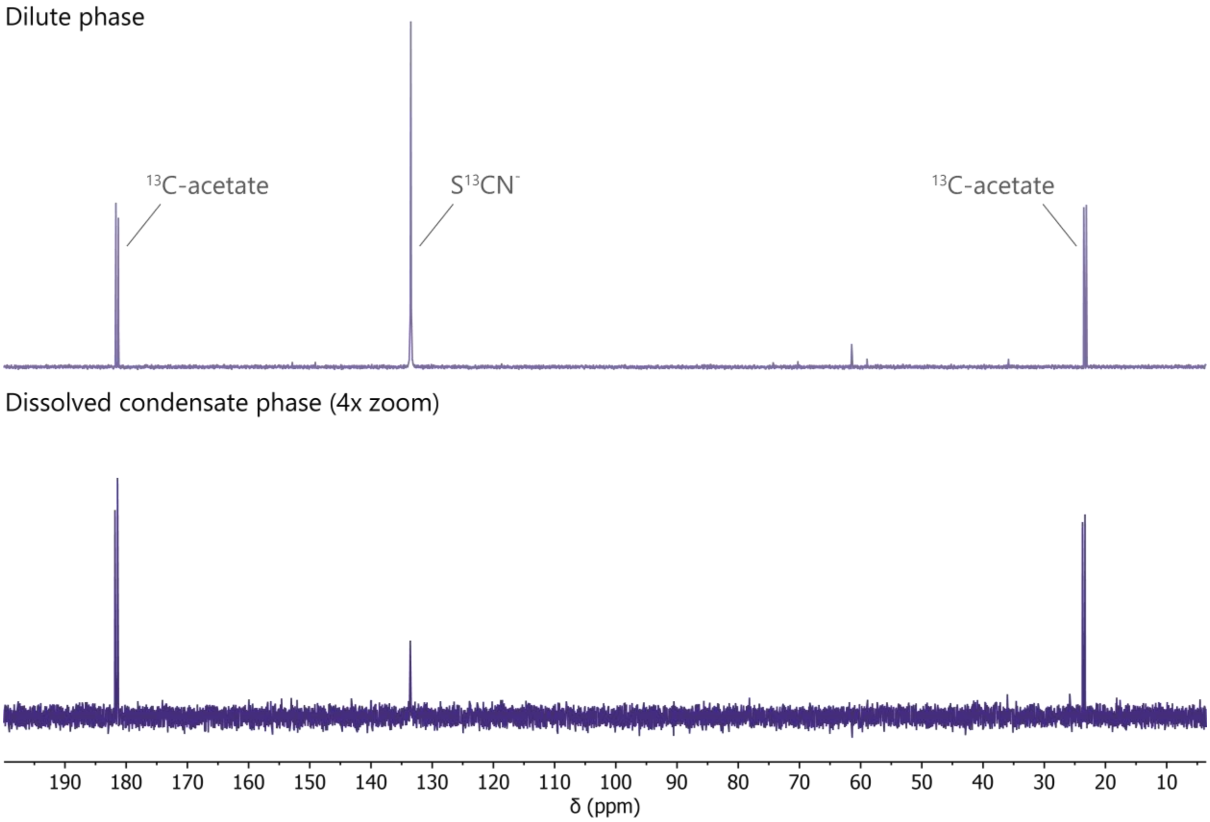

**Supplementary Figure 40:** $^{13}$C-NMR-spectra of the dilute phase and dissolved condensate phase for partitioning of 100 mM KS$^{13}$CN in condensates of 1 mM protamine chloride / 25 mM ATP in 50 mM Tris pH 8.5, used to determine the concentration of S$^{13}$CN$^-$ in reference to a sodium acetate-1,2-$^{13}C_2$ internal standard.

**Supplementary Table 9:** Measured charge-based concentrations in the condensate phase and dilute phase and $K_P$ for KS$^{13}$CN partitioning. The concentration of Na$^+$ was not determined because the condensates were dissolved in 1 M NaBr.

| Compound | $c_{condensate}$ (mM) | $c_{dilute}$ (mM) | $K_P$ |
|---|---|---|---|
| Protamine | 2334.0 ± 24.4 | 7.71 ± 0.05 | 302.6 ± 3.8 |
| ATP | 2292.0 ± 23.9 | 84.8 ± 1.3 | 27.0 ± 0.5 |
| Tris | 58.2 ± 0.9 | 52.3 ± 0.6 | 1.11 ± 0.02 |
| $K^+$ | 79.2 ± 14.3 | 100.3 ± 0.7 | 0.79 ± 0.14 |
| $Cl^-$ | 54.8 ± 8.3 | 39.3 ± 0.9 | 1.39 ± 0.21 |
| S$^{13}$CN$^-$ | 245.0 ± 3.6 | 95.4 ± 2.7 | 2.57 ± 0.08 |

##### 3.4.2.7. $MgCl_2$

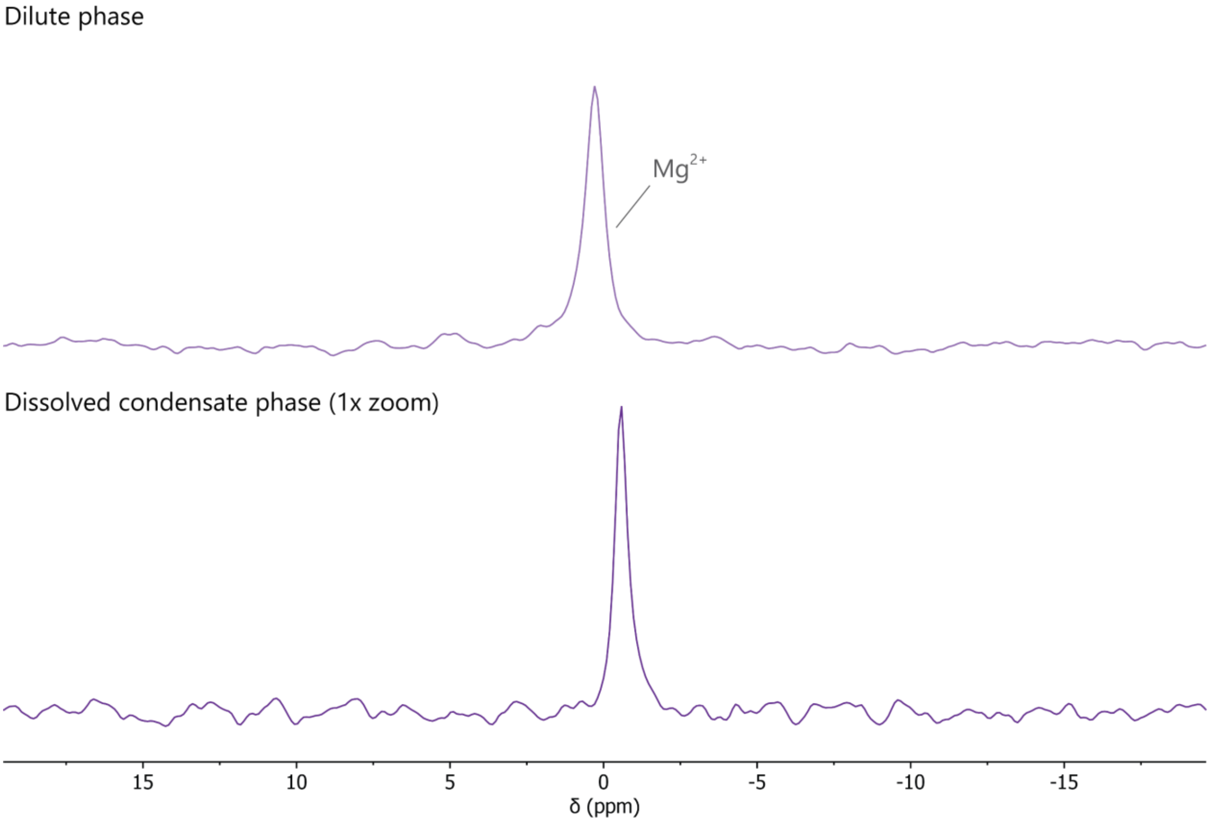

**Supplementary Figure 41:** $^{25}$Mg-NMR-spectra of the dilute phase and dissolved condensate phase for partitioning of 100 mM $MgCl_2$ (charge-based) in condensates of 1 mM protamine chloride / 25 mM ATP in 50 mM Tris pH 8.5, used to determine the concentration of $Mg^{2+}$.

**Supplementary Table 10:** Measured charge-based concentrations in the condensate phase and dilute phase and $K_P$ for $MgCl_2$ partitioning. The concentrations of $Mg^{2+}$ are from a separate sample made with different stock solutions. The protamine, ATP, Tris, $Na^+$ and $Cl^-$ concentrations were determined for a sample where the condensate phase was dissolved in 1 M KBr; the $Mg^{2+}$ concentration for a sample where the condensate phase was dissolved in 5 M HCl.

| Compound | $c_{condensate}$ (mM) | $c_{dilute}$ (mM) | $K_P$ |
|---|---|---|---|
| Protamine | 1553.7 ± 12.8 | 11.44 ± 0.06 | 135.8 ± 1.3 |
| ATP | 2952.1 ± 31.5 | 81.2 ± 0.6 | 36.4 ± 0.5 |
| Tris | 47.6 ± 1.04 | 52.1 ± 0.2 | 0.91 ± 0.02 |
| $Na^+$ | 119.7 ± 1.5 | 94.1 ± 1.1 | 1.27 ± 0.02 |
| $Cl^-$ | 304.7 ± 11.4 | 140.7 ±1.3 | 2.17 ± 0.08 |
| $Mg^{2+}$ | 2023.9 ± 57.2 | 118.0 ± 1.6 | 17.1 ± 0.5 |

3.4.2.8. LiCl

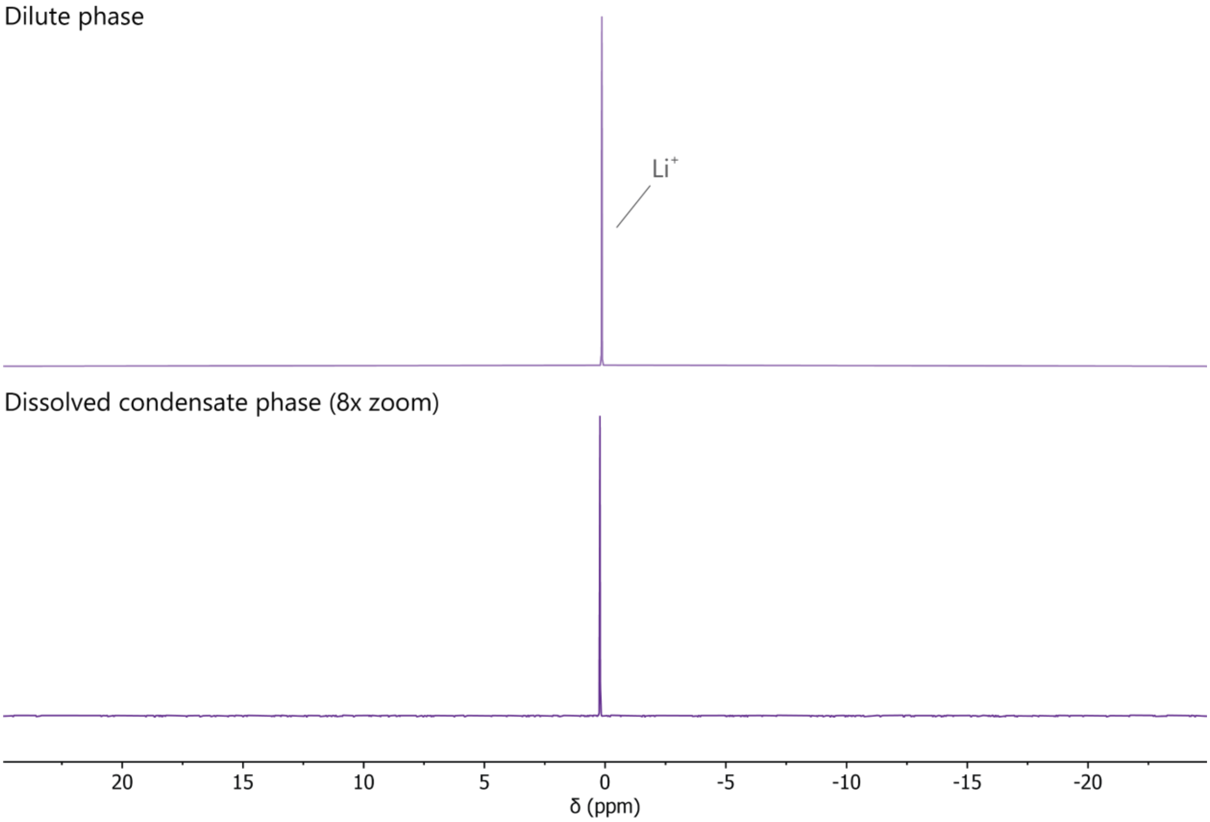

**Supplementary Figure 42:** $^{7}$Li-NMR-spectra of the dilute phase and dissolved condensate phase for partitioning of 100 mM LiCl in condensates of 1 mM protamine chloride / 25 mM ATP in 50 mM Tris pH 8.5, used to determine the concentration of $Li^+$.

**Supplementary Table 11:** Measured charge-based concentrations in the condensate phase and dilute phase and $K_P$ for LiCl partitioning.

| Compound | $c_{condensate}$ (mM) | $c_{dilute}$ (mM) | $K_P$ |
|---|---|---|---|
| Protamine | 2004.4 ± 9.3 | 8.04 ± 0.02 | 249.2 ± 1.3 |
| ATP | 2239.7 ± 98.7 | 83.8 ± 1.3 | 26.7 ± 1.3 |
| Tris | 56.5 ± 0.8 | 52.1 ± 0.2 | 1.09 ± 0.01 |
| $Na^+$ | 149.5 ± 2.4 | 96.0 ± 1.6 | 1.56 ± 0.04 |
| $Cl^-$ | 299.6 ± 5.8 | 143.2 ± 0.6 | 2.09 ± 0.04 |
| $Li^+$ | 189.2 ± 2.4 | 99.8 ± 0.3 | 1.90 ± 0.02 |

3.4.2.9. NaCl

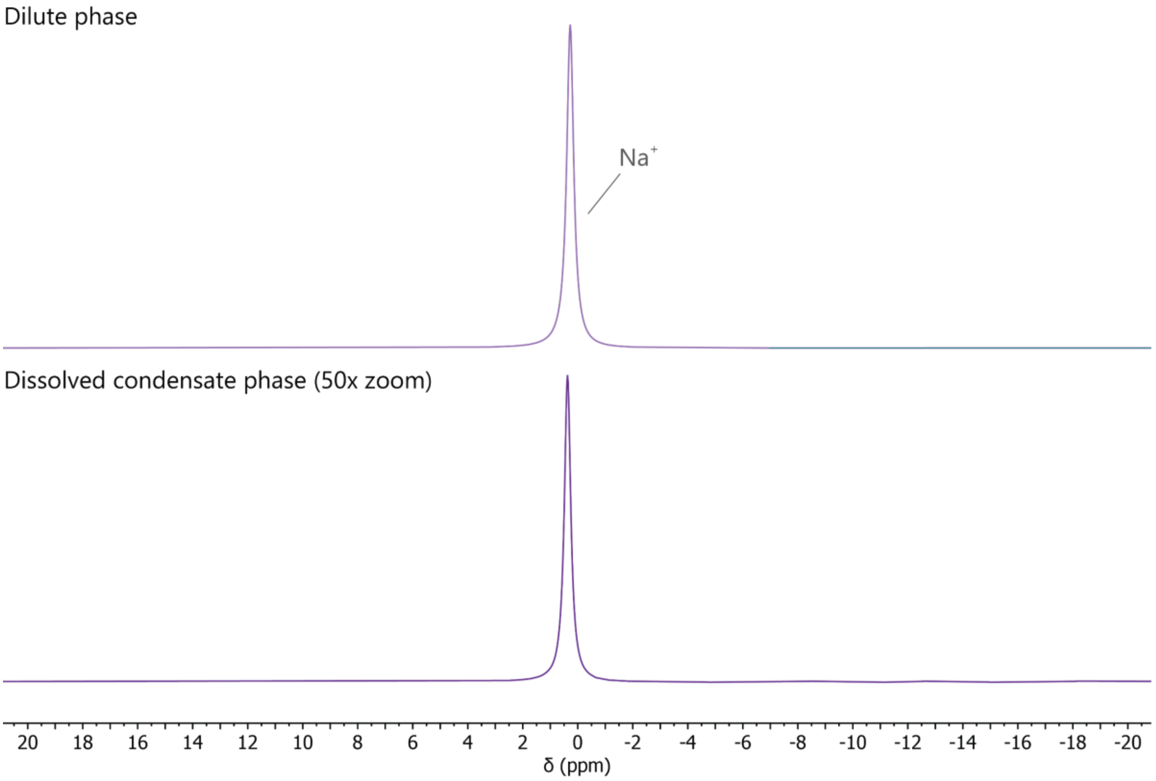

**Supplementary Figure 43:** $^{23}Na$-NMR-spectra of the dilute phase and dissolved condensate phase for partitioning of 100 mM NaCl in condensates of 1 mM protamine chloride / 25 mM ATP in 50 mM Tris pH 8.5, used to determine the concentration of $Na^{+}$.

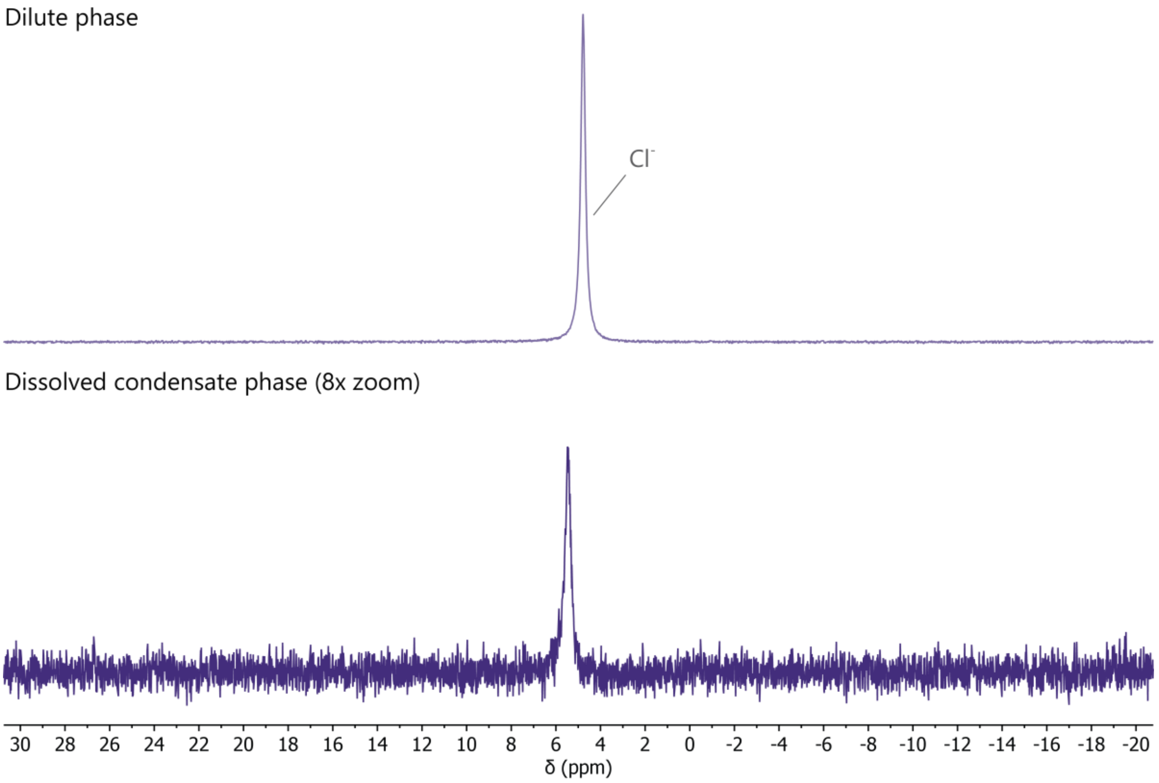

**Supplementary Figure 44:** $^{35}Cl$-NMR-spectra of the dilute phase and dissolved condensate phase for partitioning of 100 mM NaCl in condensates of 1 mM protamine chloride / 25 mM ATP in 50 mM Tris pH 8.5, used to determine the concentration of $Cl^{-}$.

**Supplementary Table 12:** Measured charge-based concentrations in the condensate phase and dilute phase and $K_P$ for NaCl partitioning for 1st repeat.

| Compound | $c_{condensate}$ (mM) | $c_{dilute}$ (mM) | $K_P$ |
|---|---|---|---|
| Protamine | 1793.8 ± 3.1 | 7.69 ± 0.06 | 233.3 ± 2.0 |
| ATP | 1908.7 ± 83.6 | 85.2 ± 1.2 | 22.4 ± 1.0 |
| Tris | 46.7 ± 0.5 | 52.5 ± 0.2 | 0.89 ± 0.01 |
| $Na^+$ | 194.5 ± 16.9 | 181.1 ± 2.2 | 1.07 ± 0.09 |
| $Cl^-$ | 292.4 ± 22.0 | 136.7 ± 4.3 | 2.14 ± 0.17 |

**Supplementary Table 13:** Measured charge-based concentrations in the condensate phase and dilute phase and $K_P$ for NaCl partitioning for 2nd repeat (with new protamine & ATP stocks).

| Compound | $c_{condensate}$ (mM) | $c_{dilute}$ (mM) | $K_P$ |
|---|---|---|---|
| Protamine | 2131.5 ± 18.9 | 7.78 ± 0.33 | 273.9 ± 12.0 |
| ATP | 2335.8 ± 55.4 | 84.9 ± 1.1 | 27.5 ± 0.7 |
| Tris | 57.2 ± 2.2 | 52.6 ± 1.7 | 1.09 ± 0.05 |
| $Na^+$ | 237.0 ± 5.2 | 198.6 ± 4.3 | 1.19 ± 0.03 |
| $Cl^-$ | 270.2 ± 4.7 | 141.2 ± 2.4 | 1.91 ± 0.04 |

3.4.2.10. KCl

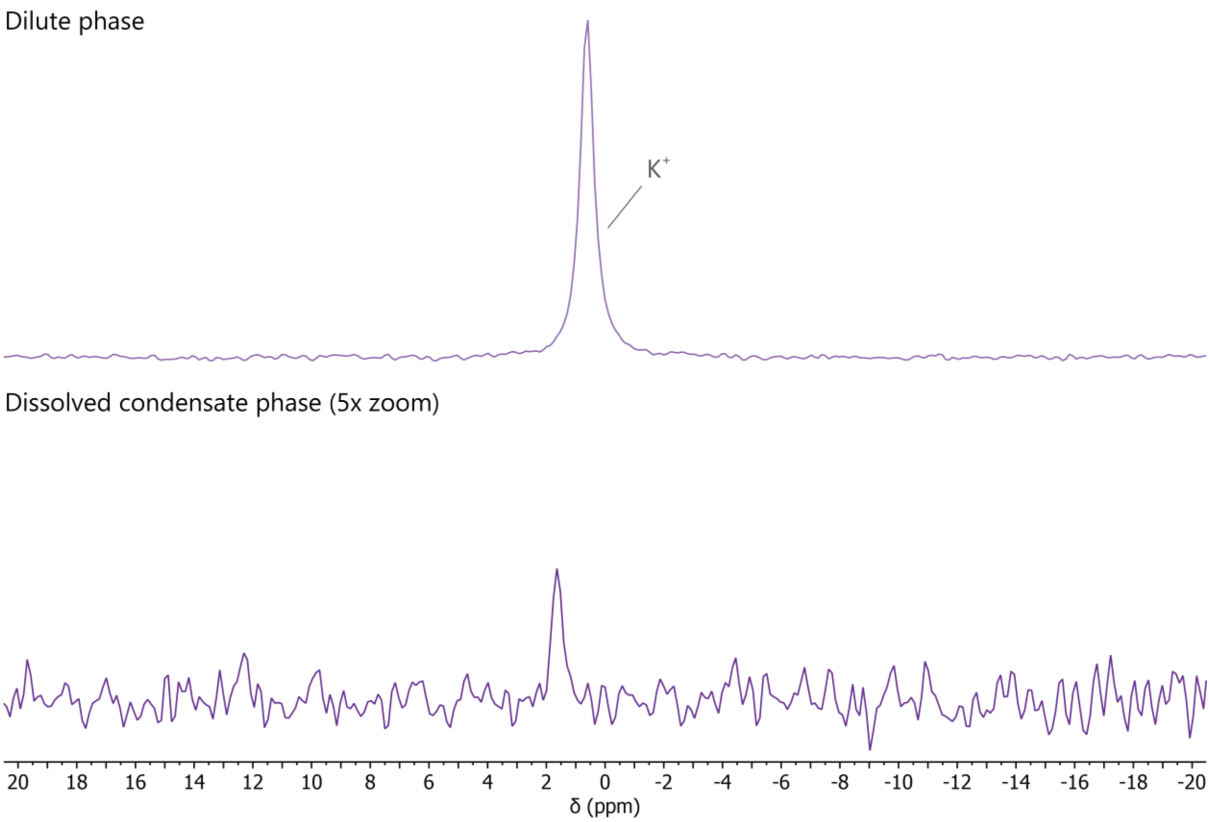

**Supplementary Figure 45:** $^{39}$K-NMR-spectra of the dilute phase and dissolved condensate phase for partitioning of 100 mM KCl in condensates of 1 mM protamine chloride / 25 mM ATP in 50 mM Tris pH 8.5, used to determine the concentration of $K^+$.

**Supplementary Table 14:** Measured charge-based concentrations in the condensate phase and dilute phase and $K_P$ for KCl partitioning. The concentration of $Na^+$ was not determined because the condensates were dissolved in 1 M NaBr.

| Compound | $c_{condensate}$ (mM) | $c_{dilute}$ (mM) | $K_P$ |
|---|---|---|---|
| Protamine | 2184.6 ± 3.3 | 7.68 ± 0.03 | 284.3 ± 1.4 |
| ATP | 2388.6 ± 61.1 | 83.9 ± 1.1 | 28.5 ± 0.8 |
| Tris | 61.1 ± 1.8 | 52.0 ± 0.3 | 1.17 ± 0.03 |
| $K^+$ | 91.7 ± 15.3 | 100.3 ± 2.9 | 0.91 ± 0.16 |
| $Cl^-$ | 166.3 ± 4.6 | 139.8 ± 2.7 | 1.19 ± 0.04 |

3.4.2.11. CsCl

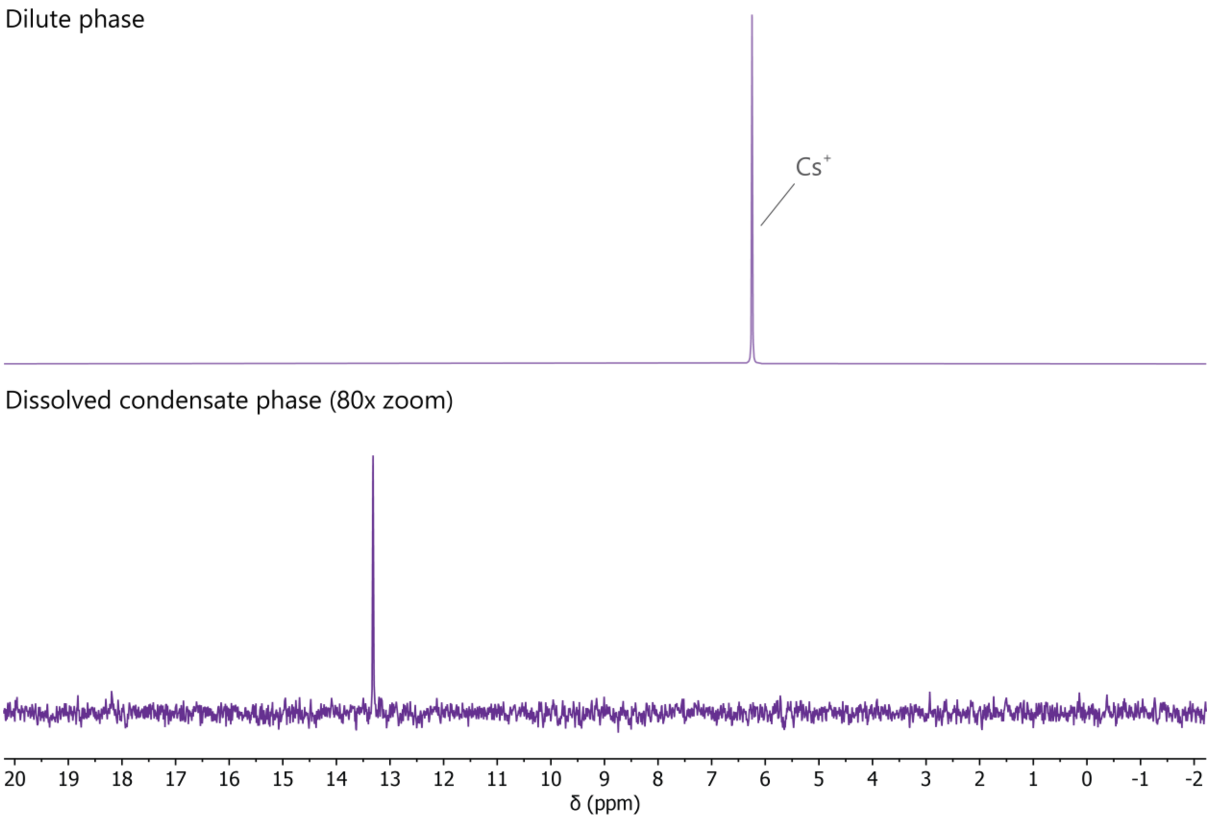

**Supplementary Figure 46:** $^{133}Cs$-NMR-spectra of the dilute phase and dissolved condensate phase for partitioning of 100 mM CsCl in condensates of 1 mM protamine chloride / 25 mM ATP in 50 mM Tris pH 8.5, used to determine the concentration of $Cs^+$.

**Supplementary Table 15:** Measured charge-based concentrations in the condensate phase and dilute phase and $K_P$ for CsCl partitioning.

| Compound | $c_{condensate}$ (mM) | $c_{dilute}$ (mM) | $K_P$ |
|---|---|---|---|
| Protamine | 2093.0 ± 3.2 | 7.34 ± 0.02 | 284.9 ± 1.0 |
| ATP | 2329.8 ± 33.5 | 78.3 ± 0.4 | 29.7 ± 0.5 |
| Tris | 58.4 ± 1.5 | 48.8 ± 0.1 | 1.20 ± 0.03 |
| $Na^+$ | 155.2 ± 2.6 | 92.2 ± 3.6 | 1.68 ± 0.07 |
| $Cl^-$ | 280.0 ± 1.0 | 140.2 ± 3.2 | 2.00 ± 0.05 |
| $Cs^+$ | 90.8 ± 1.4 | 103.1 ± 1.8 | 0.88 ± 0.02 |

##### 3.4.3. Supplementary figures ion partitioning

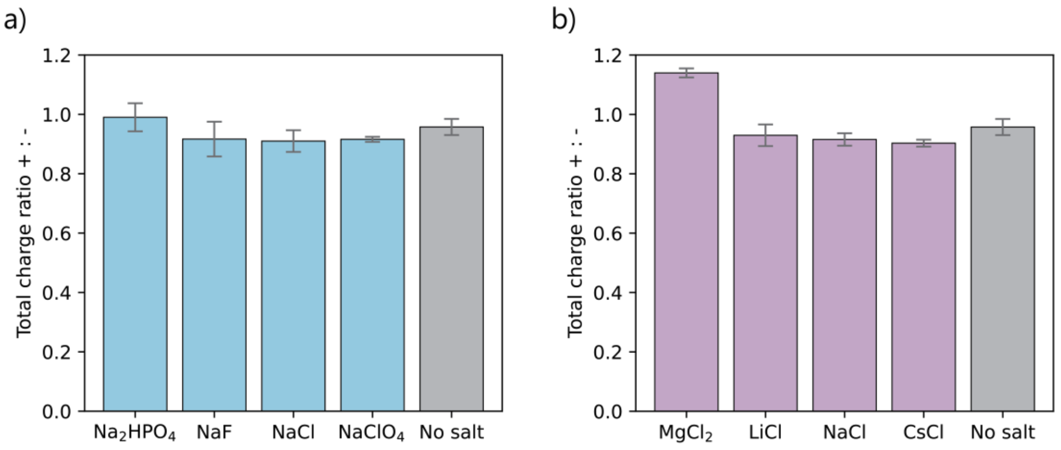

**Supplementary Figure 47:** Total charge ratio of the protamine/ATP condensates $\sum(c_{protamine} + c_{cations} + c_{TrisH^+})/\sum(c_{ATP} + c_{anions})$, charge-based concentrations, showing that only divalent ions give a significant change in the total charge ratio of the condensates. NaBr, $KS^{13}CN$ and KCl were left out of the comparison because these condensates were dissolved in a salt that contained either $Na^+$ or $Cl^-$, because of which the total charge balance could not be determined. For $MgCl_2$, the concentration of $Mg^{2+}$ was determined for another sample than the concentration of the other condensate components, which may slightly skew the data.

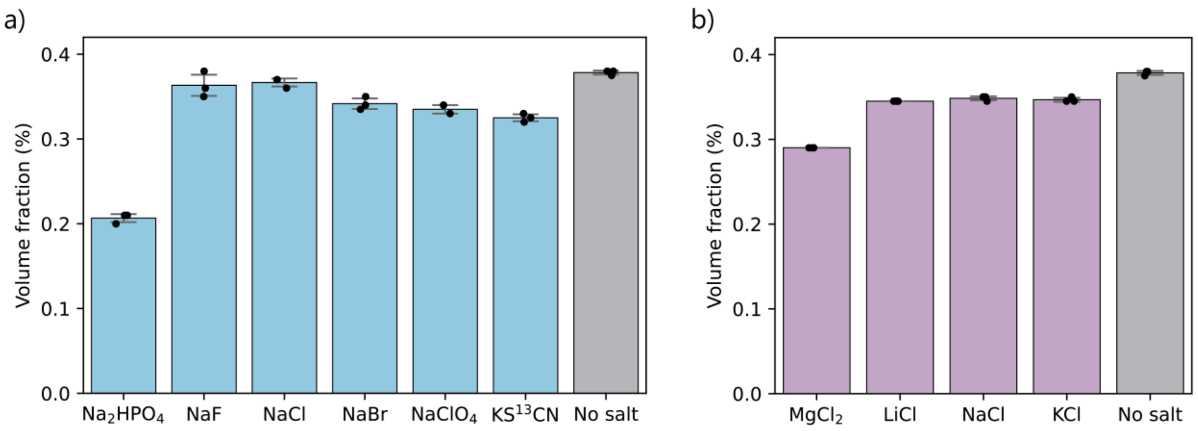

**Supplementary Figure 48:** Volume fraction of protamine/ATP condensates with 100 mM salt (charged-based, except $Na_2HPO_4$ which is at 200 mM charge-based) as determined by cell counting tubes. **a)** Strong-binding chaotropic anions give a reduction in the volume fraction, as does the divalent phosphate. **b)** Strong-binding cations give a reduction in the volume fraction. CsCl was left out of this comparison because the sample was made with a different protamine and ATP stock.

##### 3.4.4. $SCN^-$ partitioning by Raman microscopy

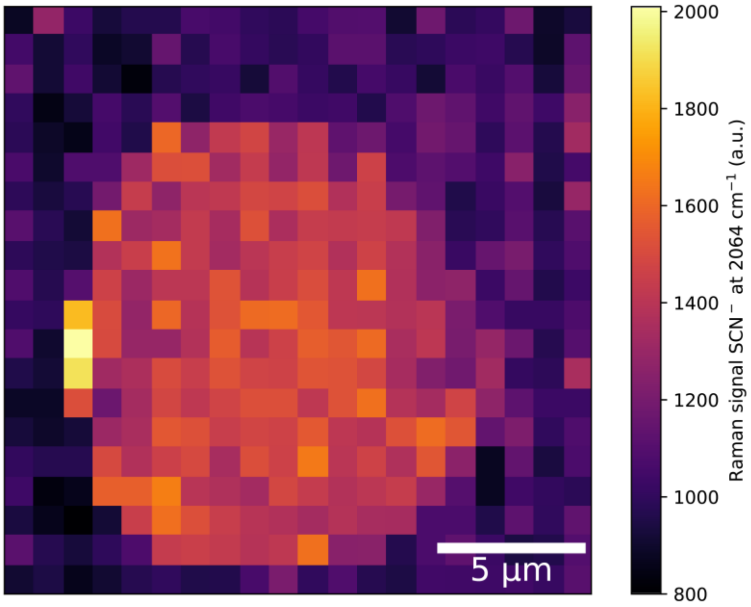

**Supplementary Figure 49:** $SCN^-$ distribution in the protamine/ATP condensate as measured by Raman microscopy, showing that $SCN^-$ is evenly distributed inside the condensate. The peak at $\tilde{\nu}$ = 2064 $cm^{-1}$ was fit with a Gaussian and integrated. Using this procedure, the $K_P$ is determined to be 1.64.

#### 3.5. Supplementary data 'Selective ion binding remodels condensate phase diagrams'

##### 3.5.1. Phase diagrams

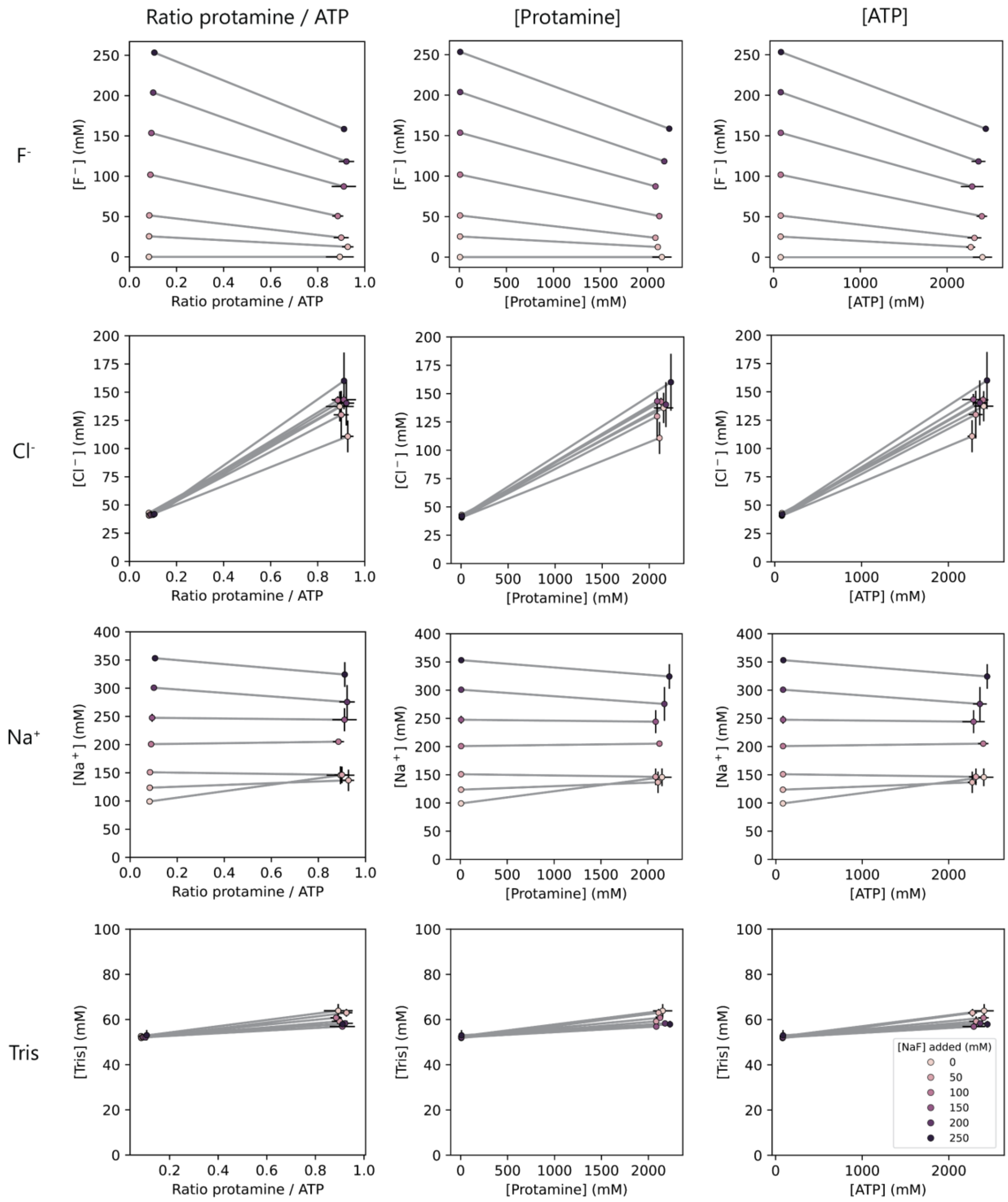

**Supplementary Figure 50:** Phase diagrams for 1 mM protamine / 25 mM ATP condensates with increasing concentrations of NaF. The plots are different representations of the same 6-dimensional dataset. Concentrations represent the charge concentrations of each species. The color of the markers represents the total concentration of NaF that was added to the sample. According to Henderson-Hasselbalch, 30.9% of the Tris is protonated $TrisH^+$ at pH 8.5. Inside the condensate phase this protonation state may be shifted.

**Supplementary Table 16:** Measured charge-based concentrations in the condensate phase for increasing concentrations of NaF.

| [NaF] added (mM) | $[\text{Protamine}]_{cond}$ (mM) | $[\text{ATP}]_{cond}$ (mM) | $[\text{Tris}]_{cond}$ (mM) | $[\text{Na}^+]_{cond}$ (mM) | $[\text{Cl}^-]_{cond}$ (mM) | $[\text{F}^-]_{cond}$ (mM) |
|---|---|---|---|---|---|---|
| 0 | 2152.0 ± 102.5 | 2407.3 ± 111.0 | 63.9 ± 3.0 | 145.7 ± 15.7 | 137.4 ± 13.3 | 0.0 ± 0.0 |
| 25 | 2108.7 ± 27.4 | 2273.1 ± 52.0 | 63.0 ± 1.7 | 136.7 ± 19.0 | 110.8 ± 14.1 | 12.4 ± 1.2 |
| 50 | 2083.5 ± 18.1 | 2315.2 ± 79.9 | 59.2 ± 2.2 | 146.3 ± 14.7 | 129.9 ± 21.1 | 23.8 ± 0.4 |
| 100 | 2124.7 ± 17.6 | 2400.2 ± 60.5 | 60.8 ± 1.3 | 205.2 ± 2.7 | 142.9 ± 3.4 | 50.6 ± 2.7 |
| 150 | 2084.4 ± 6.8 | 2288.2 ± 129.7 | 56.9 ± 0.7 | 244.1 ± 20.4 | 143.3 ± 4.9 | 87.3 ± 0.9 |
| 200 | 2177.3 ± 23.2 | 2361.9 ± 80.2 | 58.3 ± 0.5 | 275.5 ± 30.1 | 140.3 ± 19.8 | 118.3 ± 1.7 |
| 250 | 2230.1 ± 12.7 | 2445.7 ± 22.9 | 57.9 ± 1.5 | 324.3 ± 21.9 | 160.1 ± 25.2 | 158.6 ± 2.2 |

**Supplementary Table 17:** Measured charge-based concentrations in the dilute phase for increasing concentrations of NaF.

| [NaF] added (mM) | $[\text{Protamine}]_{dil}$ (mM) | $[\text{ATP}]_{dil}$ (mM) | $[\text{Tris}]_{dil}$ (mM) | $[\text{Na}^+]_{dil}$ (mM) | $[\text{Cl}^-]_{dil}$ (mM) | $[\text{F}^-]_{dil}$ (mM) |
|---|---|---|---|---|---|---|
| 0 | 6.8 ± 0.08 | 81.8 ± 0.4 | 52.6 ± 0.4 | 99.2 ± 0.4 | 43.0 ± 0.5 | 0.0 ± 0.0 |
| 25 | 6.8 ± 0.04 | 81.3 ± 0.6 | 51.8 ± 0.1 | 123.5 ± 0.7 | 40.7 ± 1.3 | 25.5 ± 0.03 |
| 50 | 6.9 ± 0.02 | 82.4 ± 1.1 | 51.9 ± 0.2 | 150.9 ± 1.9 | 40.8 ± 0.9 | 51.4 ± 0.1 |
| 100 | 7.4 ± 0.04 | 82.6 ± 0.2 | 52.3 ± 0.2 | 200.8 ± 1.8 | 41.6 ± 0.6 | 101.8 ± 0.2 |
| 150 | 7.8 ± 0.04 | 83.7 ± 1.1 | 52.1 ± 0.2 | 247.5 ± 7.1 | 41.2 ± 0.4 | 153.6 ± 0.4 |
| 200 | 8.4 ± 0.07 | 83.3 ± 0.7 | 52.1 ± 0.2 | 300.8 ± 5.2 | 41.3 ± 1.0 | 203.7 ± 0.3 |
| 250 | 9.1 ± 0.4 | 86.2 ± 4.0 | 53.1 ± 2.3 | 353.2 ± 0.9 | 41.9 ± 0.6 | 253.3 ± 1.0 |

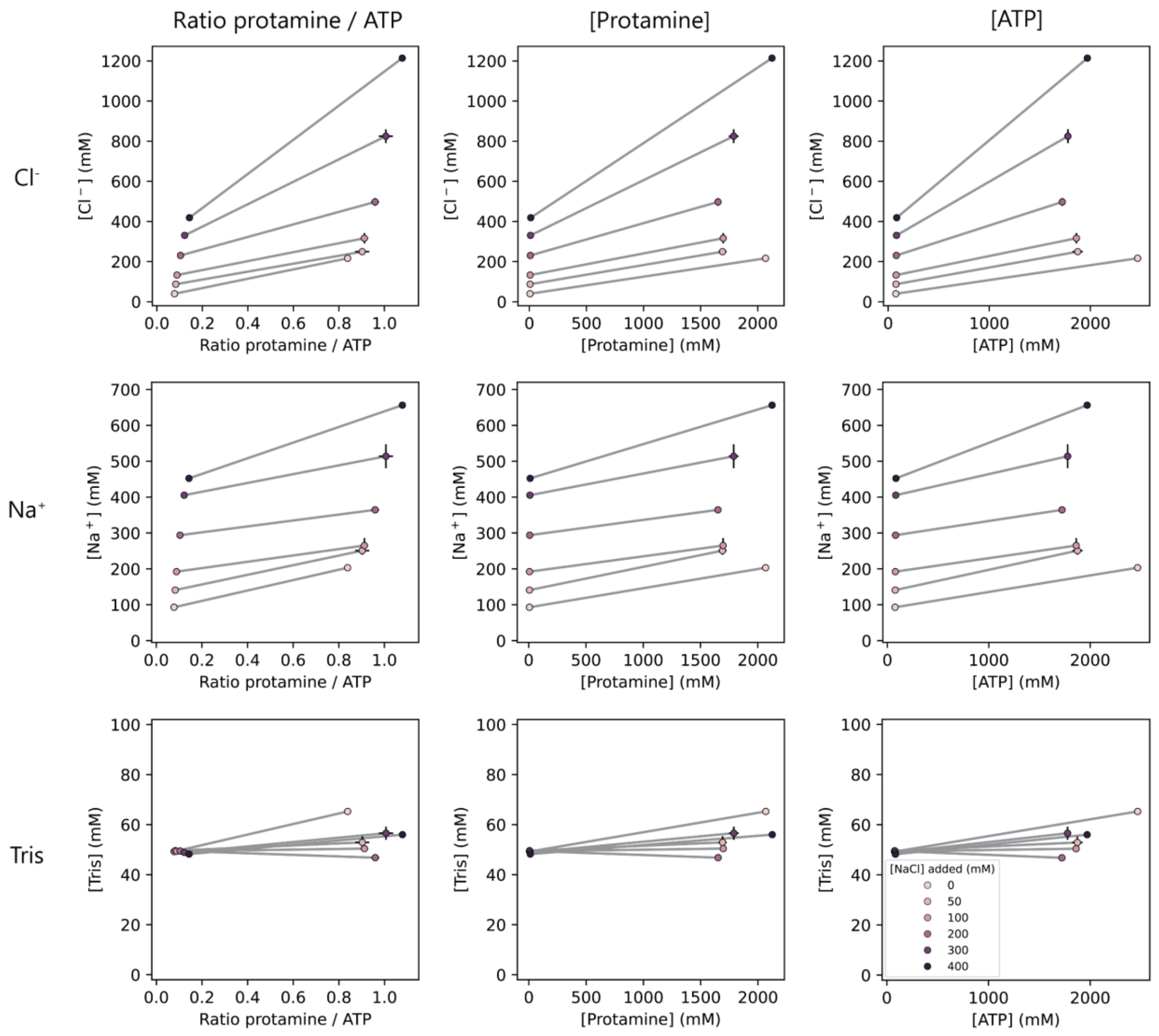

**Supplementary Figure 51:** Phase diagrams for 1 mM protamine / 25 mM ATP condensates with increasing concentrations of NaCl. The plots are different representations of the same 5-dimensional dataset. Concentrations represent the charge concentrations of each species. The color of the markers represents the total concentration of NaCl that was added to the sample. According to Henderson-Hasselbalch, 30.9% of the Tris is protonated $TrisH^+$ at pH 8.5. Inside the condensate phase this protonation state may be shifted.

**Supplementary Table 18:** Measured charge-based concentrations in the condensate phase for increasing concentrations of NaCl.

| [NaCl] added (mM) | $[Protamine]_{cond}$ (mM) | $[ATP]_{cond}$ (mM) | $[Tris]_{cond}$ (mM) | $[Na^+]_{cond}$ (mM) | $[Cl^-]_{cond}$ (mM) |
|---|---|---|---|---|---|
| 0 | 2068.8 ± 9.0 | 2467.2 ± 35.9 | 65.3 ± 0.5 | 203.3 ± 6.5 | 216.4 ± 10.1 |
| 50 | 1691.9 ± 36.0 | 1874.6 ± 51.0 | 52.9 ± 2.4 | 250.9 ± 11.7 | 249.4 ± 3.9 |
| 100 | 1697.7 ± 20.8 | 1861.7 ± 24.1 | 50.4 ± 1.0 | 265.0 ± 20.9 | 316.4 ± 25.6 |
| 200 | 1652.3 ± 6.1 | 1722.4 ± 30.0 | 46.8 ± 0.1 | 364.5 ± 2.1 | 497.4 ± 19.0 |
| 300 | 1789.8 ± 42.2 | 1778.0 ± 35.3 | 56.6 ± 2.6 | 513.9 ± 33.4 | 825.1 ± 34.3 |
| 400 | 2125.1 | 1970.3 | 56.0 | 656.3 | 1214.0 |

**Supplementary Table 19:** Measured charge-based concentrations in the dilute phase for increasing concentrations of NaCl.

| [NaCl] added (mM) | $[Protamine]_{dil}$ (mM) | $[ATP]_{dil}$ (mM) | $[Tris]_{dil}$ (mM) | $[Na^+]_{dil}$ (mM) | $[Cl^-]_{dil}$ (mM) |
|---|---|---|---|---|---|
| 0 | 5.9 ± 0.05 | 75.9 ± 1.8 | 49.2 ± 0.3 | 93.0 ± 0.8 | 39.6 ± 0.8 |
| 50 | 6.4 ± 0.03 | 77.4 ± 1.0 | 49.6 ± 0.03 | 140.9 ± 3.5 | 87.2 ± 1.6 |
| 100 | 7.0 ± 0.05 | 78.8 ± 0.4 | 49.3 ± 0.03 | 192.3 ± 8.0 | 132.9 ± 4.9 |
| 200 | 8.4 ± 0.1 | 80.6 ± 0.8 | 49.5 ± 0.2 | 293.9 ± 6.1 | 230.5 ± 8.6 |
| 300 | 10.1 ± 0.04 | 82.5 ± 0.3 | 48.9 ± 0.2 | 405.4 ± 0.9 | 330.7 ± 0.02 |
| 400 | 12.1 ± 0.05 | 84.4 ± 0.1 | 48.2 ± 0.2 | 452.3 ± 7.5 | 418.7 ± 1.5 |

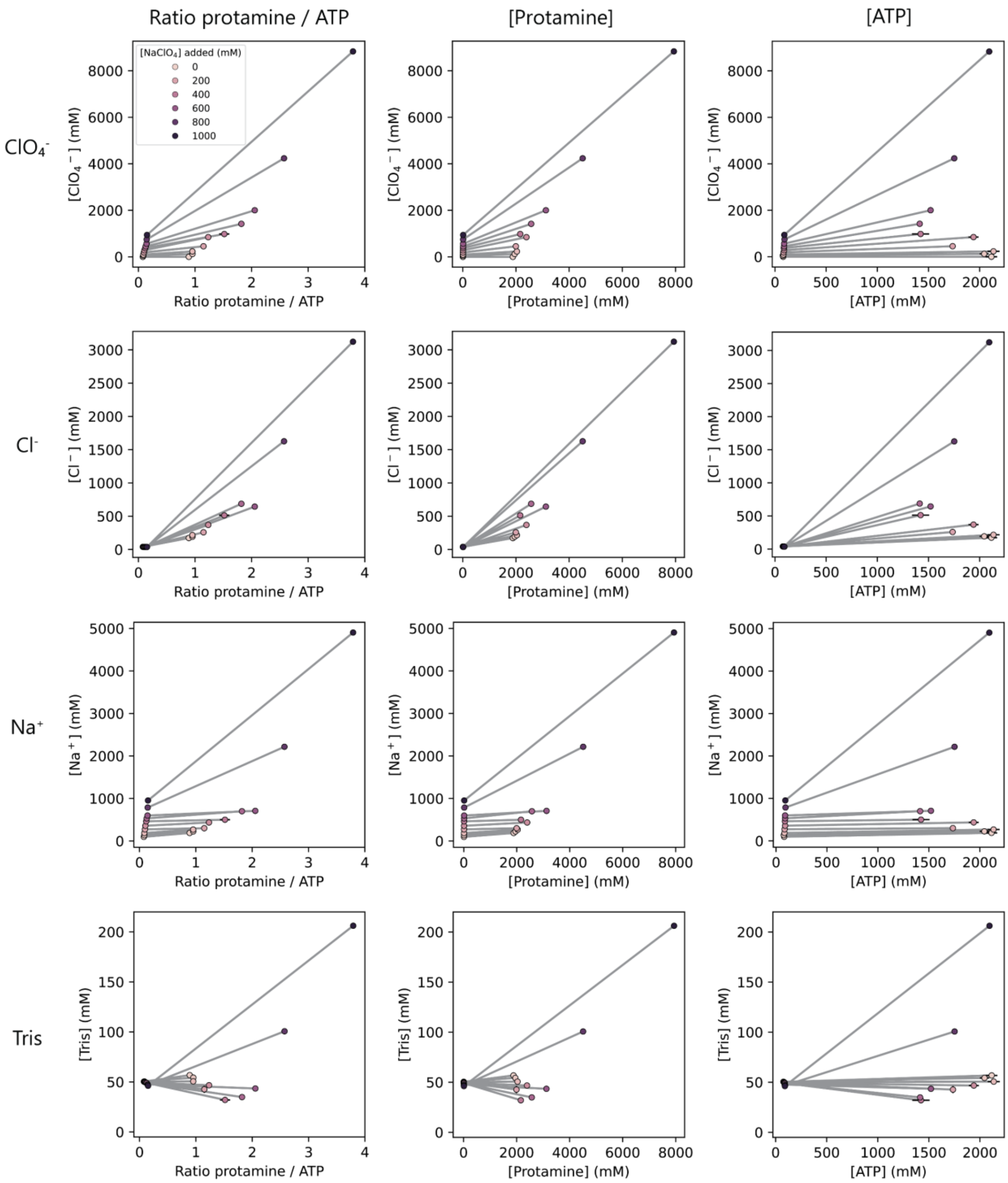

**Supplementary Figure 52:** Phase diagrams for 1 mM protamine / 25 mM ATP condensates with increasing concentrations of $NaClO_4$. The plots are different representations of the same 6-dimensional dataset. Concentrations represent the charge concentrations of each species. The color of the markers represents the total concentration of $NaClO_4$ that was added to the sample. According to Henderson-Hasselbalch, 30.9% of the Tris is protonated $TrisH^+$ at pH 8.5. Inside the condensate phase this protonation state may be shifted.

**Supplementary Table 20:** Measured charge-based concentrations in the condensate phase for increasing concentrations of $NaClO_4$.

| $[NaClO_4]$ added (mM) | $[Protamine]_{cond}$ (mM) | $[ATP]_{cond}$ (mM) | $[Tris]_{cond}$ (mM) | $[Na^+]_{cond}$ (mM) | $[Cl^-]_{cond}$ (mM) | $[ClO_4^-]_{cond}$ (mM) |
|---|---|---|---|---|---|---|
| 0 | 1883.1 ± 3.7 | 2114.7 ± 56.2 | 56.8 ± 0.5 | 188.2 ± 3.1 | 172.3 ± 12.2 | 0.0 ± 0.0 |
| 50 | 1952.1 ± 35.0 | 2044.6 ± 45.1 | 54.3 ± 0.9 | 224.5 ± 6.8 | 192.5 ± 18.7 | 124.1 ± 0.3 |
| 100 | 2034.9 ± 16.6 | 2135.0 ± 59.4 | 50.6 ± 1.7 | 267.2 ± 7.1 | 215.8 ± 9.6 | 232.6 ± 2.9 |
| 200 | 1995.4 ± 4.3 | 1735.8 ± 17.9 | 42.7 ± 3.6 | 300.1 ± 18.4 | 258.3 ± 1.7 | 453.3 ± 14.2 |
| 300 | 2392.7 ± 9.4 | 1939.9 ± 49.3 | 46.7 ± 1.7 | 435.3 ± 14.1 | 369.2 ± 6.3 | 846.2 ± 20.4 |
| 400 | 2159.3 ± 14.4 | 1422.9 ± 84.0 | 32.0 ± 1.2 | 500.0 ± 22.3 | 512.6 ± 11.9 | 980.1 ± 3.1 |
| 500 | 2572.2 ± 27.7 | 1414.1 ± 16.6 | 34.9 ± 0.1 | 699.7 ± 15.6 | 687.2 ± 38.9 | 1418.6 ± 9.5 |
| 600 | 3124.8 | 1520.3 | 43.4 | 708.2 | 644.3 | 2001.9 |
| 800 | 4507.5 | 1751.7 | 100.7 | 2215.9 | 1626.6 | 4236.3 |
| 1000 | 7937.5 | 2093.3 | 206.2 | 4904.5 | 3122.9 | 8831.0 |

**Supplementary Table 21:** Measured charge-based concentrations in the dilute phase for increasing concentrations of $NaClO_4$.

| $[NaClO_4]$ added (mM) | $[Protamine]_{dil}$ (mM) | $[ATP]_{dil}$ (mM) | $[Tris]_{dil}$ (mM) | $[Na^+]_{dil}$ (mM) | $[Cl^-]_{dil}$ (mM) | $[ClO_4^-]_{dil}$ (mM) |
|---|---|---|---|---|---|---|
| 0 | 6.2 ± 0.02 | 79.7 ± 0.9 | 50.3 ± 0.03 | 96.6 ± 4.6 | 39.2 ± 0.5 | 0.0 ± 0.0 |
| 50 | 6.6 ± 0.09 | 76.2 ± 0.1 | 50.5 ± 0.3 | 137.3 ± 2.9 | 39.5 ± 0.2 | 48.9 ± 1.2 |
| 100 | 7.1 ± 0.03 | 82.3 ± 0.6 | 50.1 ± 0.4 | 186.2 ± 2.5 | 39.3 ± 0.2 | 95.7 ± 0.4 |
| 200 | 8.2 ± 0.02 | 83.9 ± 0.9 | 49.9 ± 0.01 | 272.8 ± 7.3 | 37.8 ± 1.3 | 188.5 ± 5.5 |
| 300 | 9.3 ± 0.00 | 88.9 ± 1.7 | 49.5 ± 0.07 | 357.6 ± 8.2 | 37.3 ± 1.4 | 281.5 ± 7.1 |
| 400 | 10.4 ± 0.06 | 86.9 ± 0.2 | 49.2 ± 0.07 | 461.7 ± 16.2 | 38.5 ± 1.0 | 375.4 ± 3.3 |
| 500 | 11.4 ± 0.01 | 88.7 ± 2.4 | 48.9 ± 0.4 | 530.6 ± 24.6 | 37.5 ± 2.1 | 451.1 ± 17.1 |
| 600 | 12.7 ± 0.03 | 88.9 ± 2.0 | 47.7 ± 0.3 | 598.1 ± 32.2 | 36.8 ± 1.1 | 561.8 ± 12.9 |
| 800 | 13.5 ± 0.16 | 92.7 ± 4.2 | 46.9 ± 0.2 | 785.7 ± 34.2 | 36.2 ± 0.9 | 746.7 ± 24.4 |
| 1000 | 13.7 ± 0.16 | 90.9 ± 1.5 | 46.2 ± 0.06 | 952.6 ± 10.2 | 37.1 ± 0.4 | 939.1 ± 0.5 |

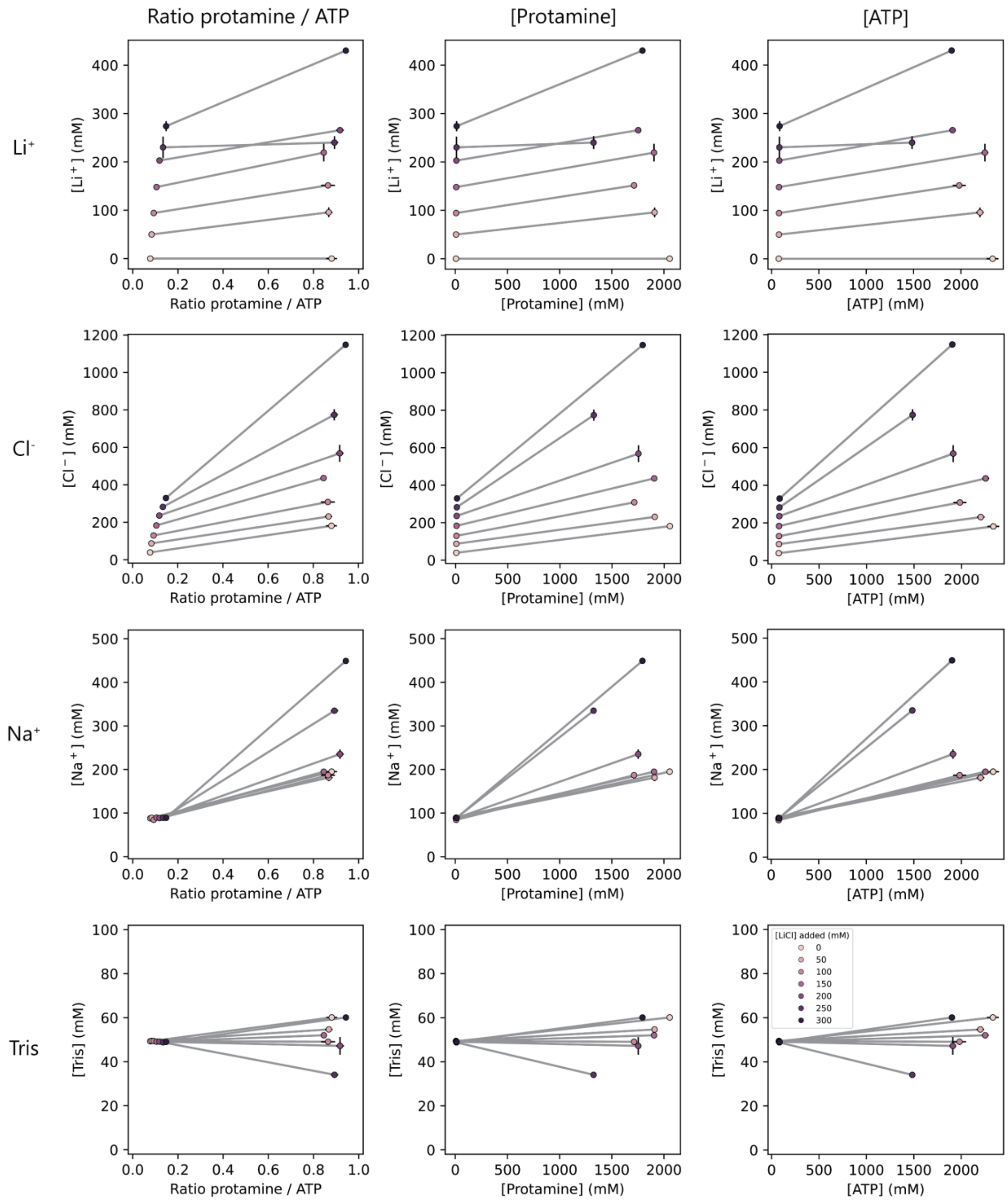

**Supplementary Figure 53:** Phase diagrams for 1 mM protamine / 25 mM ATP condensates with increasing concentrations of LiCl. The plots are different representations of the same 6-dimensional dataset. Concentrations represent the charge concentrations of each species. The color of the markers represents the total concentration of LiCl that was added to the sample. According to Henderson-Hasselbalch, 30.9% of the Tris is protonated $TrisH^+$ at pH 8.5. Inside the condensate phase this protonation state may be shifted.

**Supplementary Table 22:** Measured charge-based concentrations in the condensate phase for increasing concentrations of LiCl.

| [LiCl] added (mM) | $[\text{Protamine}]_{cond}$ (mM) | $[\text{ATP}]_{cond}$ (mM) | $[\text{Tris}]_{cond}$ (mM) | $[\text{Na}^+]_{cond}$ (mM) | $[\text{Cl}^-]_{cond}$ (mM) | $[\text{Li}^+]_{cond}$ (mM) |
|---|---|---|---|---|---|---|
| 0 | 2054.9 ± 10.8 | 2333.9 ± 62.3 | 60.1 ± 0.1 | 195.0 ± 7.4 | 181.4 ± 6.9 | 0.0 ± 0.0 |
| 50 | 1910.3 ± 9.9 | 2201.6 ± 40.5 | 54.7 ± 1.0 | 181.4 ± 8.3 | 230.9 ± 8.9 | 95.7 ± 10.1 |
| 100 | 1713.6 ± 15.8 | 1982.4 ± 69.0 | 49.1 ± 0.6 | 186.9 ± 7.2 | 308.4 ± 2.0 | 151.4 ± 1.3 |
| 150 | 1904.9 ± 4.1 | 2252.8 ± 37.0 | 52.0 ± 1.3 | 194.9 ± 3.2 | 436.3 ± 12.7 | 219.2 ± 18.2 |
| 200 | 1753.6 ± 0.4 | 1911.5 ± 34.8 | 47.2 ± 4.0 | 235.3 ± 10.8 | 568.2 ± 45.1 | 265.6 ± 0.2 |
| 250 | 1325.4 ± 5.3 | 1484.2 ± 26.8 | 34.1 ± 0.3 | 334.9 ± 3.0 | 773.9 ± 29.9 | 239.9 ± 13.2 |
| 300 | 1794.4 | 1902.1 | 60.1 | 449.1 | 1147.8 | 430.1 |

**Supplementary Table 23:** Measured charge-based concentrations in the dilute phase for increasing concentrations of LiCl.

| [LiCl] added (mM) | $[\text{Protamine}]_{dil}$ (mM) | $[\text{ATP}]_{dil}$ (mM) | $[\text{Tris}]_{dil}$ (mM) | $[\text{Na}^+]_{dil}$ (mM) | $[\text{Cl}^-]_{dil}$ (mM) | $[\text{Li}^+]_{dil}$ (mM) |
|---|---|---|---|---|---|---|
| 0 | 5.9 ± 0.02 | 76.4 ± 0.2 | 49.3 ± 0.1 | 87.9 ± 2.8 | 39.1 ± 1.4 | 0.0 ± 0.0 |
| 50 | 6.5 ± 0.04 | 78.1 ± 0.2 | 49.5 ± 0.1 | 89.1 ± 0.5 | 87.4 ± 0.8 | 49.9 ± 2.4 |
| 100 | 7.3 ± 0.05 | 78.9 ± 0.1 | 49.4 ± 0.2 | 84.5 ± 1.1 | 129.7 ± 0.7 | 94.3 ± 0.2 |
| 150 | 8.3 ± 0.03 | 79.1 ± 0.2 | 49.1 ± 0.2 | 89.6 ± 0.6 | 183.2 ± 1.6 | 147.9 ± 0.8 |
| 200 | 9.7 ± 0.14 | 82.1 ± 0.6 | 49.1 ± 0.4 | 88.5 ± 1.0 | 236.0 ± 0.8 | 202.8 ± 0.7 |
| 250 | 11.1 ± 0.03 | 82.8 ± 0.2 | 48.8 ± 0.1 | 89.0 ± 1.2 | 282.1 ± 2.2 | 230.1 ± 22.3 |
| 300 | 12.6 ± 0.05 | 85.3 ± 0.7 | 49.0 ± 0.3 | 89.0 ± 1.8 | 329.5 ± 0.2 | 273.9 ± 10.5 |

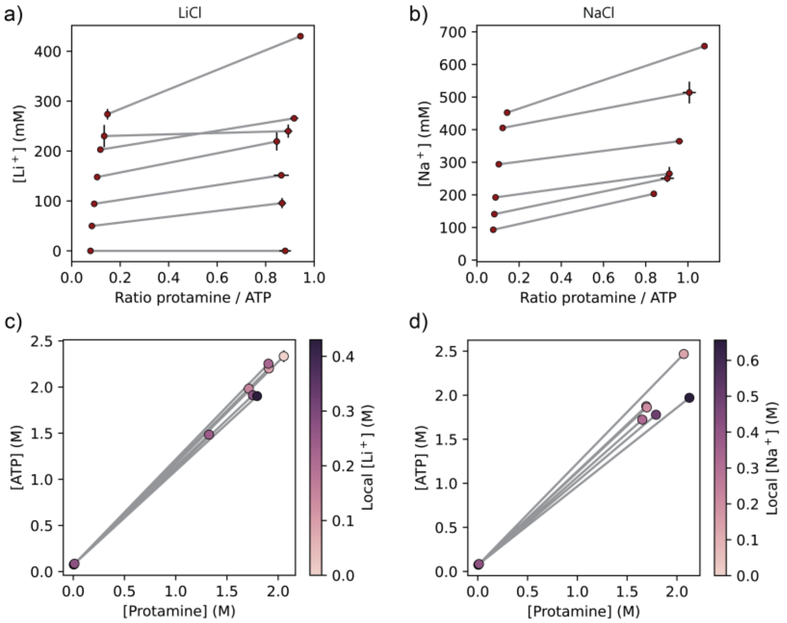

**Supplementary Figure 54:** Phase diagrams for 1 mM protamine / 25 mM ATP condensates with increasing concentrations of LiCl (a, c) or NaCl (b, d). **a)** $Li^+$ is localized to the condensates and the partitioning stays approximately equal for increasing concentrations of $Li^+$. **b)** $Na^+$ is localized to the condensates and the partitioning stays approximately equal for increasing concentrations of $Na^+$. **c)** Addition of LiCl does not change the ratio of protamine : ATP inside the condensates. **d)** Addition of NaCl mildly changes the ratio of protamine : ATP, shifting the ratio towards more protamine at high NaCl concentrations.

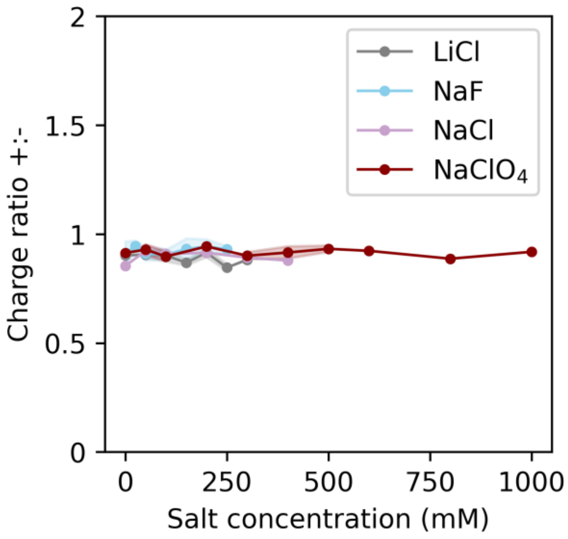

**Supplementary Figure 55:** The overall charge ratio of the protamine/ATP condensates stays equal for different concentrations of LiCl, NaF, NaCl and $NaClO_4$.

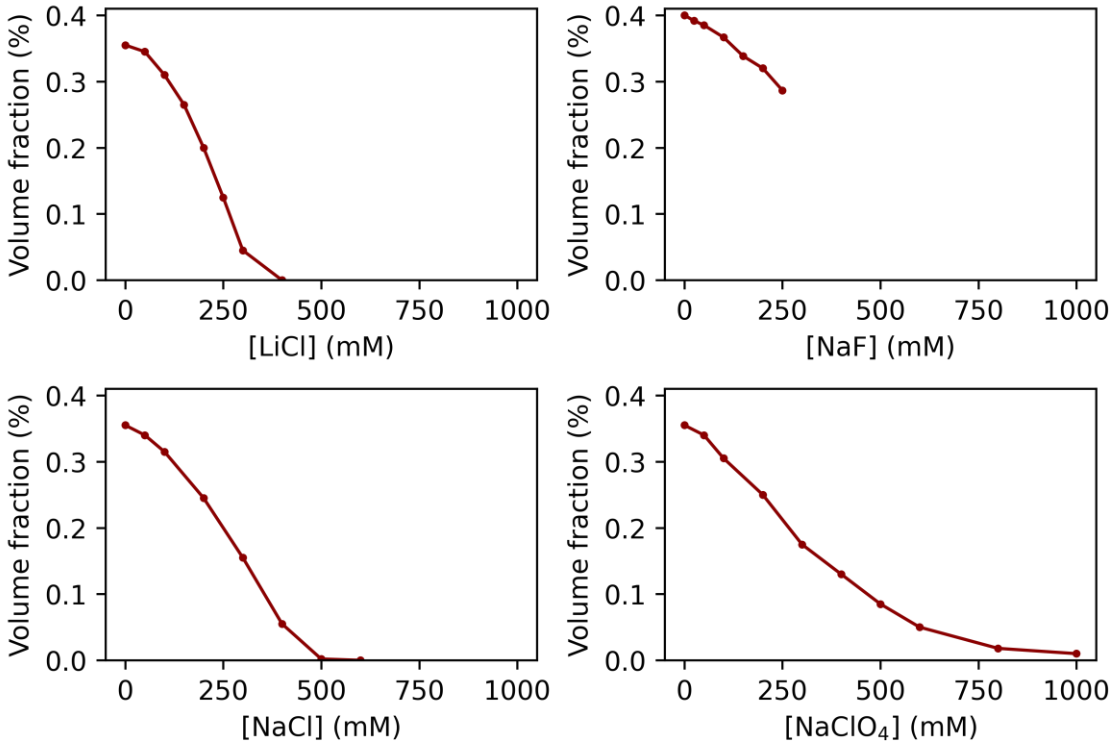

**Supplementary Figure 56:** Changes in condensate volume fraction as a function of salt concentration for LiCl, NaF, NaCl and $NaClO_4$. Condensates dissolve more readily for LiCl and NaCl than for $NaClO_4$.

##### 3.5.2. [ATP]$_{crit}$ protamine/ATP

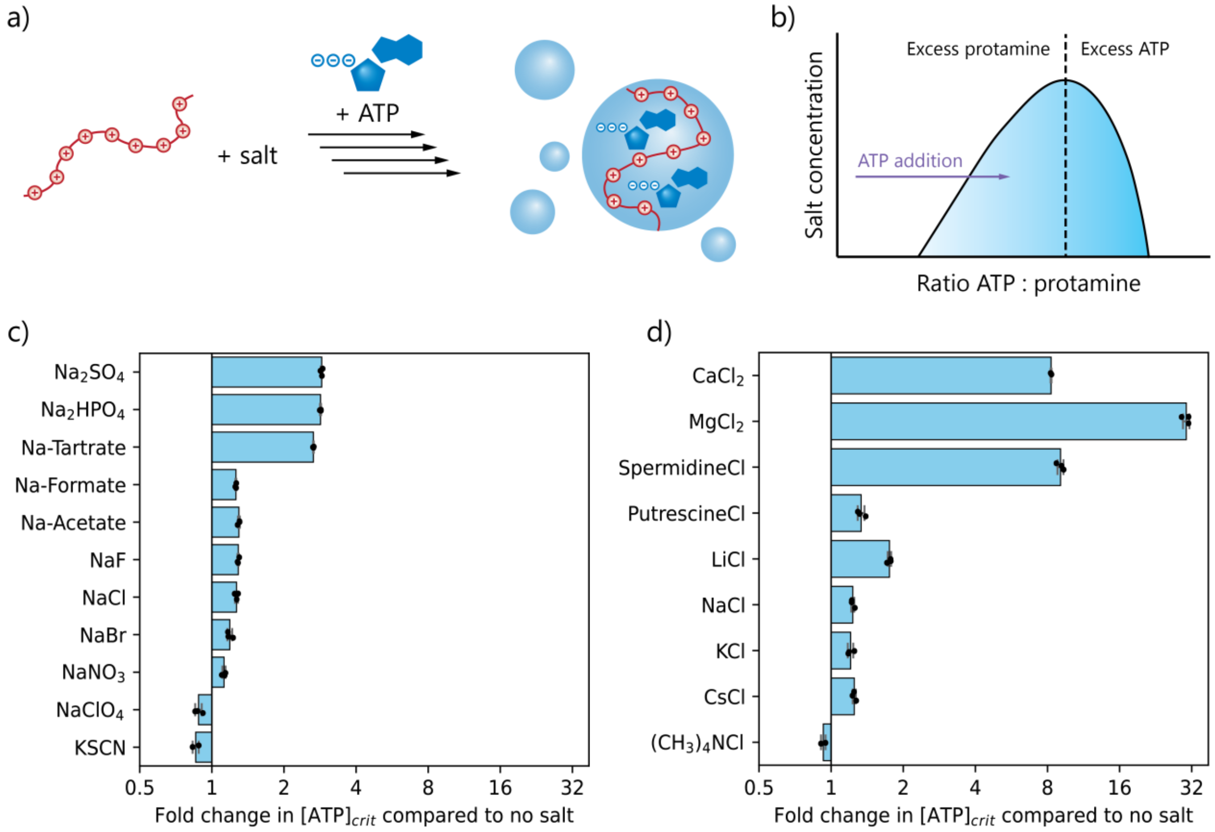

**Supplementary Figure 57:** Fold-change in critical concentration of ATP ([ATP]$_{crit}$) required for formation of protamine/ATP coacervates. 0.5 mM protamine was mixed with 100 mM salt and ATP was titrated in. The onset of turbidity is reported as [ATP]$_{crit}$. **a)** Anions that bind to protamine lower the [ATP]$_{crit}$, because they shield the positive charge on protamine and thereby lower the excess of positive charge. **b)** Cations that bind to ATP significantly increase the [ATP]$_{crit}$, because they shield the negative charge on ATP and thereby increase the excess of positive charge. Ions that do not bind specifically increase the [ATP]$_{crit}$ because of aspecific charge-screening effects. Interestingly, tetramethylammonia, which is highly chaotropic and thus binds ATP weakly lowers the [ATP]$_{crit}$, most likely due to its high concentration it outcompetes the stronger binding of ATP's counterion sodium. It can also clearly be seen that the effect of multivalent ions is significantly stronger than that of monovalent ions, and all multivalent ions increase the [ATP]$_{crit}$, even phosphate which does bind to protamine.

##### 3.5.3. [Polyion]$_{crit}$ of other heterotypic condensates

To assess whether our observations for the effect of ions on protamine/ATP condensate stability are general, we determined the [polyion]$_{crit}$ for different types of heterotypic condensates (protamine/polyA, protamine/$D_{30}$ and $K_{30}$/$D_{30}$), by titrating in either the polycation or the polyanion (Supplementary Figure 58 & 59). We observe that for many of these condensate systems, addition of 100 mM of any salt always lowers the [polyion]$_{crit}$, indicating that for these condensates made with longer polymers, a small amount of salt helps to lower charge-repulsion within polymer chains and favors phase separation. Still, we observe a similar trend between different ions as for protamine/ATP: salt ions that bind specifically to the polyion in excess lower the critical concentration of the oppositely charged polyion, while ions that bind to the polyion that is titrated in increase the [polyion]$_{crit}$. These observations show that the effect of specific ion binding on condensate stability is a general effect for different heterotypic condensates.

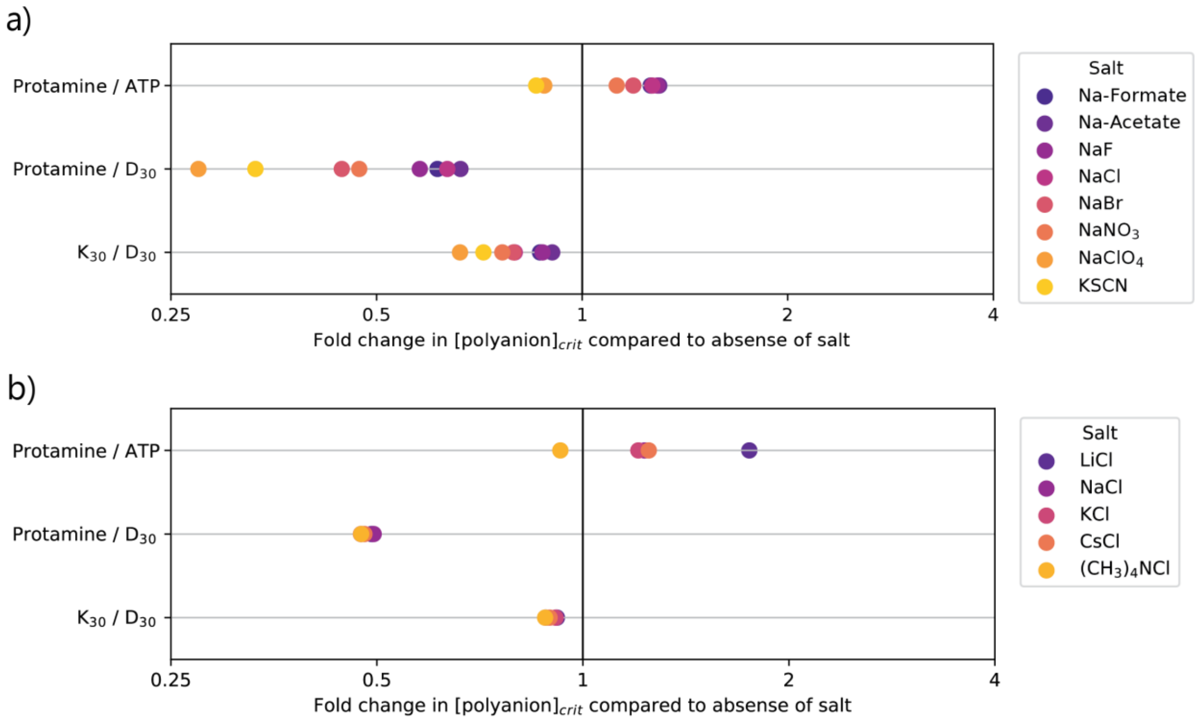

**Supplementary Figure 58:** Fold-change in critical polyanion concentration ($[\text{polyanion}]_{\text{crit}}$) for formation of different charge-based condensates for 100 mM of different monovalent salts, with respect to the critical concentration in absence of salt. All systems were studied at pH 8.5. Anions that bind to the polycation have a stronger tendency to lower the $[\text{polyanion}]_{\text{crit}}$, while cations that bind to the polyanion have a stronger tendency to increase the $[\text{polyanion}]_{\text{crit}}$.

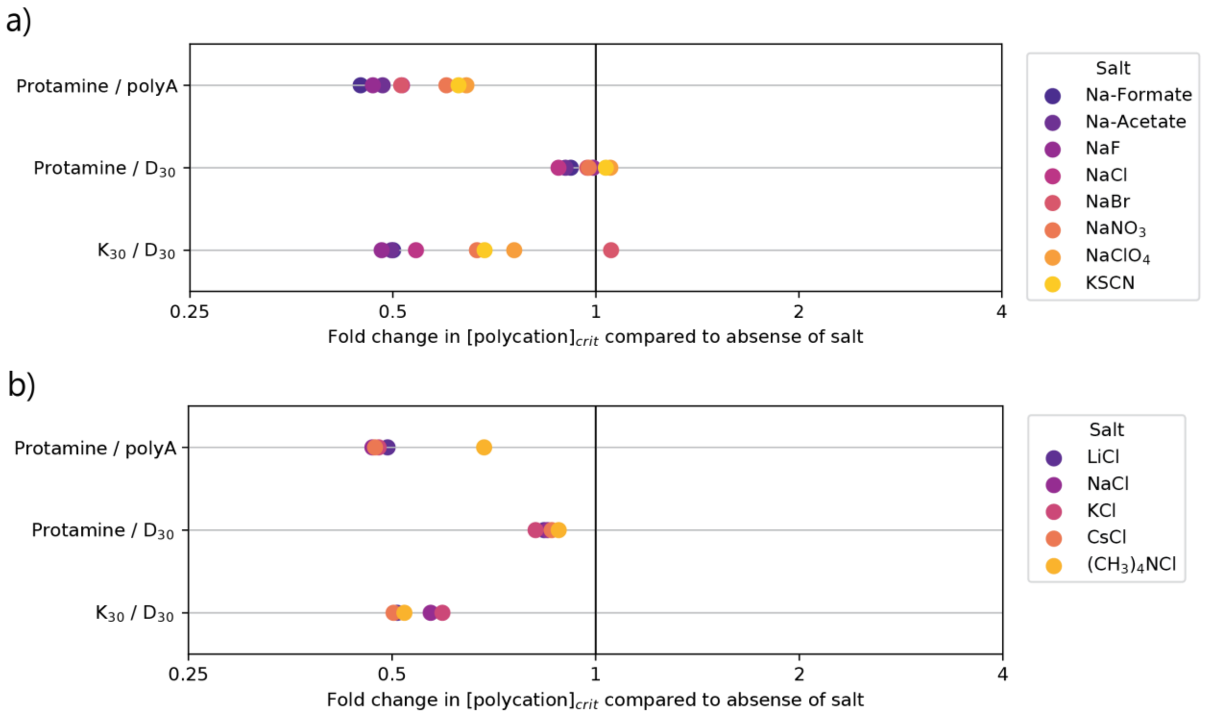

**Supplementary Figure 59:** Fold-change in critical polycation concentration ($[\text{polycation}]_{\text{crit}}$) for formation of different charge-based condensates for 100 mM of different monovalent salts, with respect to the critical concentration in absence of salt. All systems were studied at pH 8.5. Anions that bind to the polycation have a stronger tendency to increase the $[\text{polycation}]_{\text{crit}}$, while cations that bind to the polyanion have a stronger tendency to lower the $[\text{polyanion}]_{\text{crit}}$.

3.5.4. Salt effects on a homotypic hydrophobic condensate

To understand the effect of different ions on homotypic, hydrophobic condensates, we investigated condensates formed by the hydrophobic peptide derivative LLssLL[1] and observe that in this case, both specific ion binding and salting out effects play a role (Supplementary Figure 60). The peptide derivative has two free N-termini which are positively charged at pH 8.5, and can therefore bind chaotropic anions. We observe a reduction in LCST in presence of perchlorate and thiocyanate, in line with binding to and subsequent neutralization of the N-termini resulting in a larger hydrophobicity and increased phase separation propensity. However, we also observe a reduction in LCST for phosphate and fluoride, indicating that also general salting-out effects play a role.

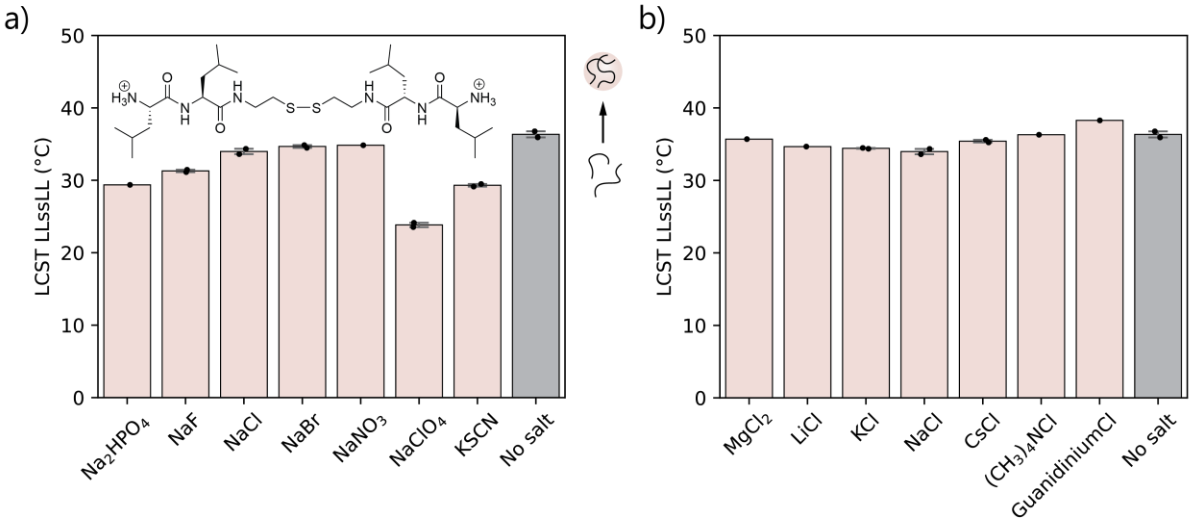

**Supplementary Figure 60:** Lower critical solution temperature (LCST) of hydrophobic condensates of 3 mg/mL peptide derivative LLssLL with 100 mM salt in 50 mM Tris pH 8.5. **a)** The LCST is lowered, i.e. condensates are stabilized, for both chaotropic and kosmotropic anions, via neutralization of the N-termini by selective chaotrope binding and the salting-out effect, respectively. **b)** Cations have a less pronounced effect on the LCST of LLssLL. Chaotropic cations slightly disfavor phase separation because they solubilize the hydrophobic peptide.

##### 3.6. Supplementary data 'Ion binding changes viscosity and interface potential of condensates'

**Supplementary Table 24:** Diffusion coefficients and viscosities for protamine/ATP condensates prepared with different salts, as measured by RICS.

| **Sample** | ***D*** **(µm/s²)** | $\eta_d$ **(mPa·s)** |
|---|---|---|
| No salt | 0.144 ± 0.006 | 1.49 ± 0.06 |
| 50 mM $MgCl_2$ | 0.124 ± 0.010 | 1.74 ± 0.13 |
| 100 mM LiCl | 0.173 ± 0.017 | 1.25 ± 0.12 |
| 100 mM NaCl | 0.171 ± 0.014 | 1.26 ± 0.11 |
| 100 mM CsCl | 0.182 ± 0.011 | 1.19 ± 0.07 |
| 50 mM $Na_2HPO_4$ | 0.177 ± 0.009 | 1.22 ± 0.06 |
| 100 mM NaF | 0.153 ± 0.015 | 1.41 ± 0.14 |
| 100 mM $NaClO_4$ | 0.158 ± 0.009 | 1.37 ± 0.08 |
| 200 mM LiCl | 0.253 ± 0.054 | 0.88 ± 0.18 |
| 200 mM NaCl | 0.232 ± 0.024 | 0.93 ± 0.09 |
| 200 mM CsCl | 0.254 ± 0.022 | 0.85 ± 0.07 |
| 200 mM NaF | 0.168 ± 0.024 | 1.30 ± 0.21 |
| 200 mM $NaClO_4$ | 0.101 ± 0.008 | 2.13 ± 0.17 |

**Supplementary Table 25:** Parameters used to calculate ζ-potentials of pLys/pGlu condensates. $c_{ion}$ is the ionic strength of the buffer (determined via Henderson-Hasselbalch), the added salt and the counterions of the protamine and ATP. $\eta_d$ was determined by RICS. $E_0$ was determined by testing the lowest electric field strength at which droplets moved. Other parameters used to calculate the ζ-potential are T = 298 K and water viscosity of 0.891 mPa·s.

| **Sample** | $c_{ion}$ **(mM)** | $\eta_d$ **(mPa·s)** | ***E (V)*** | $E_0$ **(V)** | **ζ (mV)** |
|---|---|---|---|---|---|
| No salt | 102.3 | 1.49 ± 0.06 | 8 | 1.9 | -4.34 ± 1.71 |
| 50 mM $MgCl_2$ | 252.8 | 1.74 ± 0.13 | 5 | 2.5 | 3.49 ± 2.61 |
| 100 mM LiCl | 202.6 | 1.25 ± 0.12 | 5 & 12 | 2.5 | -0.40 ± 0.21 |
| 100 mM NaCl | 202.6 | 1.26 ± 0.11 | 5 & 8 | 2.5 | -1.07 ± 0.73 |
| 100 mM CsCl | 203.3 | 1.19 ± 0.07 | 5 & 8 | 2.5 | -0.81 ± 0.54 |
| 50 mM $Na_2HPO_4$ | 249.1 | 1.22 ± 0.06 | 5 | 1.5 | -1.91 ± 0.57 |
| 100 mM NaF | 203.2 | 1.41 ± 0.14 | 5 & 8 | 2.5 | -2.17 ± 0.96 |
| 100 mM $NaClO_4$ | 202.3 | 1.37 ± 0.08 | 5 & 12 | 2.5 | -1.19 ± 0.31 |
| 200 mM LiCl | 302.9 | 0.88 ± 0.18 | 5 | 2.5 | 0.96 ± 0.50 |
| 200 mM NaCl | 302.8 | 0.93 ± 0.09 | 5 | 2.5 | -0.68 ± 0.24 |
| 200 mM CsCl | 304.2 | 0.85 ± 0.07 | 5 & 8 | 2.5 | -0.31 ± 0.08 |
| 200 mM NaF | 304.1 | 1.30 ± 0.21 | 5 & 8 | 2.5 | -1.46 ± 0.11 |
| 200 mM $NaClO_4$ | 302.3 | 2.13 ± 0.17 | 5 & 8 | 2.5 | 0.02 ± 0.11 |

##### 3.7. Supplementary data ‘Ion binding alters RNA duplex stability in condensates’

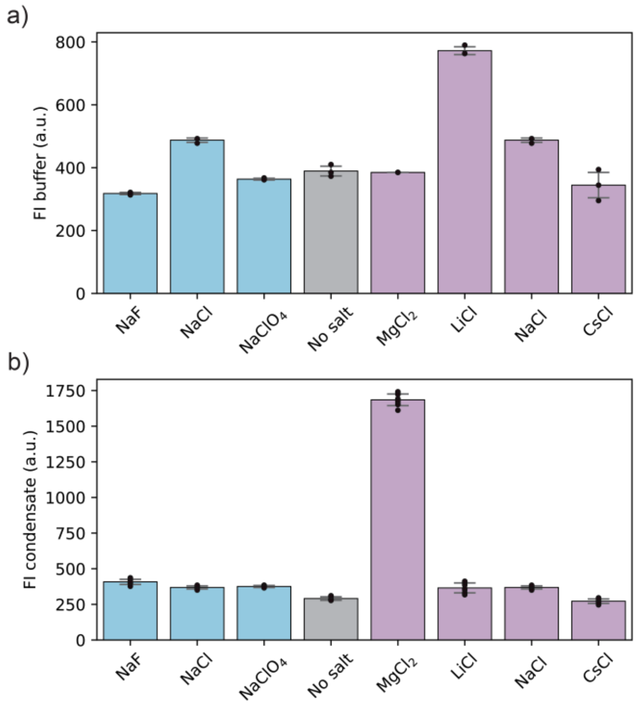

**Supplementary Figure 61:** Cy3-Fluorescence intensity (FI) of the RNA FRET pair in buffer and in protamine/ATP condensates. **a)** In buffer all fluorescence intensities are in the same order of magnitude. **b)** Inside the condensate, however, the fluorescence intensity is significantly higher in presence of $MgCl_2$, indicating that the presence of $MgCl_2$ enhances the partitioning of RNA into the condensates.

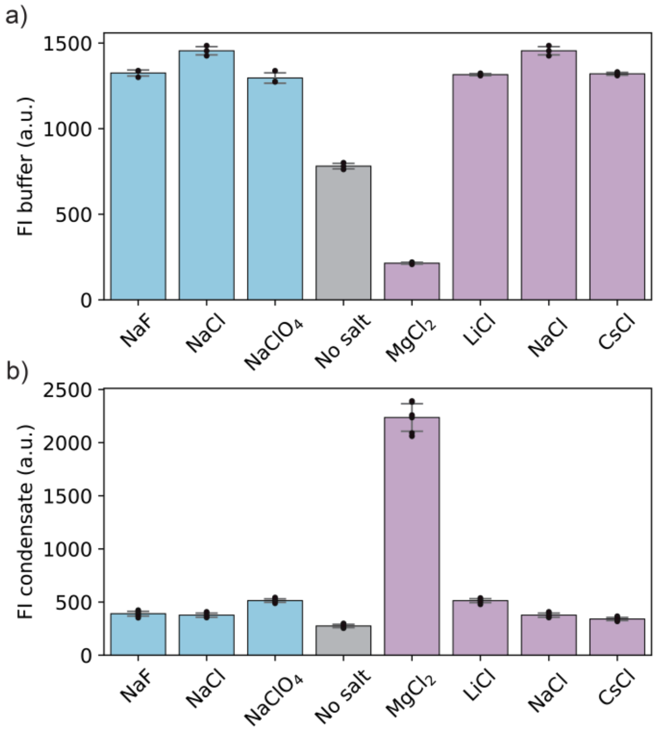

**Supplementary Figure 62:** Cy3-Fluorescence intensity (FI) of the DNA FRET pair in buffer and in protamine/ATP condensates. **a)** In buffer all fluorescence intensities are in the same order of magnitude, except for $MgCl_2$, which seems to lower the fluorescence intensity. **b)** Inside the condensate, however, the fluorescence intensity is significantly higher in presence of $MgCl_2$, indicating that the presence of $MgCl_2$ enhances the partitioning of DNA into the condensates. Either quenching by $Mg^{2+}$ is less pronounced in the condensates, or the increased partitioning of DNA in presence of $MgCl_2$ is still an underestimation.

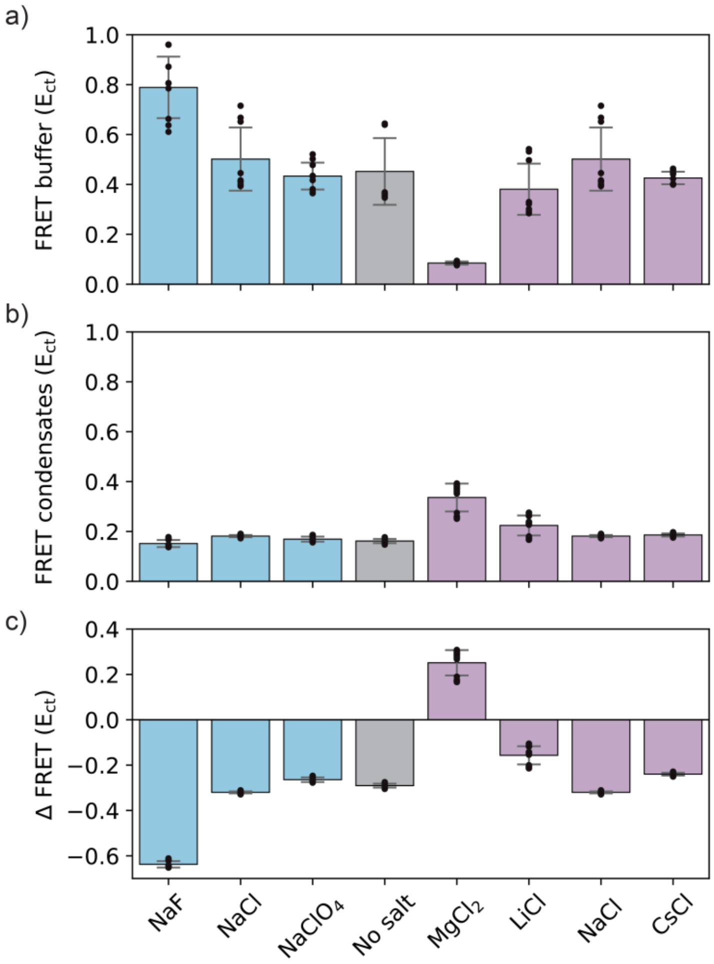

**Supplementary Figure 63:** FRET intensities for the Cy3/Cy5-labelled DNA duplex in buffer and in protamine/ATP condensates. **a)** FRET efficiency in buffer, **b)** FRET efficiency in protamine/ATP condensates, **c)** difference in FRET efficiency between condensates and buffer (ΔFRET = FRET efficiency condensate – FRET efficiency buffer). A similar trend is observed as for RNA: kosmotropic anions stabilize the duplex in buffer, while the kosmotropic cations $Mg^{2+}$ and $Li^{+}$ destabilize the duplex. In condensates, the trend for the cations is reversed compared to buffer, similar to what we observed for RNA. The duplex stabilization by $F^{-}$, however, disappears for DNA inside condensates.

### 4. Extended Discussion: rethinking charge-charge interactions

In this paper, we observed that charge-charge interactions between condensate components or between condensate components and salt ions follow the law of matching water affinities, resulting in strong binding between two chaotropes or two kosmotropes, but weak binding between ions with very different hydration strengths. These observations show that the entropic rearrangement of water molecules during ion complexation has an important contribution to charge-charge interactions, and therefore charge-charge interactions should not be considered as purely enthalpic, as is the naïve interpretation. This prompts us to rethink charge-charge interactions in the context of biomolecular condensates and may help to explain the thermodynamic properties of charge-based condensate formation.

Because all organic anions are chaotropic and all organic cations are kosmotropic, interaction between polymeric condensate components is inherently unfavorable in terms of ion hydration according to the LMWA, as it involves association of two ions with different water affinities. Interactions with typical counterions (chloride, sodium) are more favorable due to more similar water affinity. This matches well with observations from literature that complexation of polyelectrolytes and complex coacervation are typically entropy-driven[18–23] and sometimes even enthalpically unfavorable.[18,21,23]

Condensate formation is therefore often explained in terms of counterion release,[18–21] although counterion release has both an entropic and enthalpic factor, which will depend on the counterion involved. If counterion-polyelectrolyte side chain interactions are stronger than side chain-side chain interactions between polyelectrolytes, then strong counterion-side chain bonds have to be broken during polyelectrolyte complexation and counterion release, explaining the observed positive or mildly negative enthalpy. Interestingly, Yang *et al.* observed that the enthalpy of polyelectrolyte complex formation is highly dependent on the anionic counterion involved, with the kosmotropic (and thus weakly binding) acetate giving a $\Delta H_{PEC}$ of -6.0 kJ/mol, while the chaotropic (and thus strongly binding) perchlorate gives $\Delta H_{PEC}$ = 12.8 kJ/mol,[24] indicating that strong counterion-polyelectrolyte interaction can make polyelectrolyte complexation enthalpically unfavorable, and is an important factor to consider in the thermodynamics of condensate formation. The observation that the presence of strongly binding ions can make condensate formation less thermodynamically favorable, matches with our observed lower $T_{crit}$ for protamine/ATP with strongly binding ions in Figure 4i.

In addition to enthalpy, ion type is also expected to influence the entropy of condensate formation. Counterion release increases the entropy of the counterions, but also involves rearrangement of water molecules in the ion's hydration shell, and, therefore, the entropy associated with counterion release will depend on the hydration strength of the ion.[25]

Altogether, it is clear that charge-based condensate formation is not just driven by enthalpy-driven electrostatic interactions, and a more nuanced description of the process is required, involving the important role of counterions and hydration water. Our quantitative analysis of interaction strengths between ions and condensate components helps elucidate and refine the role of salt and counterions in condensate formation.

### 5. Data repository

Raw data can be found on the Radboud Data repository.

### 6. Supplementary references

(1) Abbas, M.; Lipiński, W. P.; Nakashima, K. K.; Huck, W. T. S.; Spruijt, E. A Short Peptide Synthon for Liquid–Liquid Phase Separation. *Nat. Chem.* **2021**, *13* (11), 1046–1054. https://doi.org/10.1038/s41557-021-00788-x.

(2) Rembert, K. B.; Paterová, J.; Heyda, J.; Hilty, C.; Jungwirth, P.; Cremer, P. S. Molecular Mechanisms of Ion-Specific Effects on Proteins. *J. Am. Chem. Soc.* **2012**, *134* (24), 10039–10046. https://doi.org/10.1021/ja301297g.

(3) Abraham, M. J.; Murtola, T.; Schulz, R.; Páll, S.; Smith, J. C.; Hess, B.; Lindah, E. Gromacs: High Performance Molecular Simulations through Multi-Level Parallelism from Laptops to Supercomputers. *SoftwareX* **2015**, *1–2*, 19–25. https://doi.org/10.1016/j.softx.2015.06.001.

(4) Piana, S.; Robustelli, P.; Tan, D.; Chen, S.; Shaw, D. E. Development of a Force Field for the Simulation of Single-Chain Proteins and Protein-Protein Complexes. *J. Chem. Theory Comput.* **2020**, *16* (4), 2494–2507. https://doi.org/10.1021/acs.jctc.9b00251.

(5) Bussi, G.; Donadio, D.; Parrinello, M. Canonical Sampling through Velocity Rescaling. *J. Chem. Phys.* **2007**, *126* (1), 14101. https://doi.org/10.1063/1.2408420.

(6) Parrinello, M.; Rahman, A. Polymorphic Transitions in Single Crystals: A New Molecular Dynamics Method. *J. Appl. Phys.* **1981**, *52* (12), 7182–7190. https://doi.org/10.1063/1.328693.

(7) Breßler, I.; Kohlbrecher, J.; Thünemann, A. F. SASfit: A Tool for Small-Angle Scattering Data Analysis Using a Library of Analytical Expressions. *J. Appl. Crystallogr.* **2015**, *48* (5), 1587–1598. https://doi.org/10.1107/S1600576715016544.

(8) Hammouda, B. Small-Angle Scattering from Branched Polymers. *Macromol. Theory Simulations* **2012**, *21* (6), 372–381. https://doi.org/10.1002/mats.201100111.

(9) Silva, E. R.; Listik, E.; Han, S. W.; Alves, W. A.; Soares, B. M.; Reza, M.; Ruokolainen, J.; Hamley, I. W. Sequence Length Dependence in Arginine/Phenylalanine Oligopeptides: Implications for Self-Assembly and Cytotoxicity. *Biophys. Chem.* **2018**, *233*, 1–12. https://doi.org/10.1016/j.bpc.2017.11.005.

(10) De Mello, L. R.; Hamley, I. W.; Castelletto, V.; Garcia, B. B. M.; Han, S. W.; De Oliveira, C. L. P.; Da Silva, E. R. Nanoscopic Structure of Complexes Formed between DNA and the Cell-Penetrating Peptide Penetratin. *J. Phys. Chem. B* **2019**, *123* (42), 8861–8871. https://doi.org/10.1021/acs.jpcb.9b05512.

(11) Smokers, I. B. A.; Spruijt, E. Quantification of Biomolecular Condensate Volume Reveals Network Swelling and Dissolution Regimes during Phase Transition. *Biomacromolecules* **2024**. https://doi.org/10.1021/acs.biomac.4c01201.

(12) Jones, G.; Dole, M. The Viscosity of Aqueous Solutions of Strong Electrolytes with Special Reference to Barium Chloride. *J. Am. Chem. Soc.* **1929**, *51* (10), 2950–2964. https://doi.org/10.1021/ja01385a012.

(13) Collins, K. D. The Behavior of Ions in Water Is Controlled by Their Water Affinity. *Q. Rev. Biophys.* **2019**, *52*, e11. https://doi.org/10.1017/S0033583519000106.

(14) Robinson, J. B.; Strottmann, J. M.; Stellwagen, E. Prediction of Neutral Salt Elution Profiles for Affinity Chromatography. *Proc. Natl. Acad. Sci. U. S. A.* **1981**, *78* (4), 2287–2291. https://doi.org/10.1073/pnas.78.4.2287.

(15) Petrášek, Z.; Schwille, P. Precise Measurement of Diffusion Coefficients Using Scanning Fluorescence Correlation Spectroscopy. *Biophys. J.* **2008**, *94* (4), 1437–1448. https://doi.org/10.1529/biophysj.107.108811.

(16) Schrimpf, W.; Barth, A.; Hendrix, J.; Lamb, D. C. PAM: A Framework for Integrated Analysis of Imaging, Single-Molecule, and Ensemble Fluorescence Data. *Biophys. J.* **2018**, *114* (7), 1518–1528. https://doi.org/10.1016/j.bpj.2018.02.035.

(17) Van Haren, M. H. I.; Visser, B. S.; Spruijt, E. Probing the Surface Charge of Condensates Using Microelectrophoresis. *Nat. Commun.* **2024**, *15* (1), 1–10. https://doi.org/10.1038/s41467-024-47885-2.

(18) Chang, L. W.; Lytle, T. K.; Radhakrishna, M.; Madinya, J. J.; Vélez, J.; Sing, C. E.; Perry, S. L. Sequence and Entropy-Based Control of Complex Coacervates. *Nat. Commun.* **2017**, *8* (1), 1–8. https://doi.org/10.1038/s41467-017-01249-1.

(19) Kayitmazer, A. B. Thermodynamics of Complex Coacervation. *Adv. Colloid Interface Sci.* **2017**, *239*, 169–177. https://doi.org/10.1016/j.cis.2016.07.006.

(20) Chen, S.; Wang, Z. G. Driving Force and Pathway in Polyelectrolyte Complex Coacervation. *Proc. Natl. Acad. Sci. U. S. A.* **2022**, *119* (36), e2209975119. https://doi.org/10.1073/pnas.2209975119.

(21) Priftis, D.; Laugel, N.; Tirrell, M. Thermodynamic Characterization of Polypeptide Complex Coacervation. *Langmuir* **2012**, *28* (45), 15947–15957. https://doi.org/10.1021/la302729r.

(22) Fu, J.; Schlenoff, J. B. Driving Forces for Oppositely Charged Polyion Association in Aqueous Solutions: Enthalpic, Entropic, but Not Electrostatic. *J. Am. Chem. Soc.* **2016**, *138* (3), 980–990. https://doi.org/10.1021/jacs.5b11878.

(23) Chowdhury, A.; Borgia, A.; Ghosh, S.; Sottini, A.; Mitra, S.; Eapen, R. S.; Borgia, M. B.; Yang, T.; Galvanetto, N.; Ivanović, M. T.; Łukijańczuk, P.; Zhu, R.; Nettels, D.; Kundagrami, A.; Schuler, B. Driving Forces of the Complex Formation between Highly Charged Disordered Proteins. *Proc. Natl. Acad. Sci. U. S. A.* **2023**, *120* (41). https://doi.org/10.1073/pnas.2304036120.

(24) Yang, M.; Digby, Z. A.; Schlenoff, J. B. Precision Doping of Polyelectrolyte Complexes: Insight on the Role of Ions. *Macromolecules* **2020**, *53* (13), 5465–5474. https://doi.org/10.1021/acs.macromol.0c00965.

(25) Shen, Y.; Li, S.; Jiang, J.; Sun, F.; Zhao, Y.; Qiao, F.; Qin, B. Ions Effect on Tunable Coacervate and Its Relevance to the Hofmeister Series. *Colloids Surfaces A Physicochem. Eng. Asp.* **2024**, *702*, 134597. https://doi.org/10.1016/j.colsurfa.2024.134597.
